# Biospeckle laser digital image processing for quantitative and statistical evaluation of the activity of Ciprofloxacin on *Escherichia coli* K-12

**DOI:** 10.1101/479345

**Authors:** Hilda Cristina Grassi, Ana Velásquez, Olga Mercedes Belandria, María Lorena Lobo-Sulbarán, Jesús E. Andrades-Grassi, Humberto Cabrera, Efrén D. J. Andrades

## Abstract

Antibiotic susceptibility testing is a necessary step prior to the treatment of clinical infections. A major concern is the time required to obtain a fast and reliable result. The aim of this work is to use Biospeckle laser in a 15min assay for an antimicrobial susceptibility test of Ciprofloxacin in serial two-fold dilutions on *Escherichia coli* K-12 using Venereal Disease Research Laboratory (VDRL) plates. Analysis of images by video edition is performed on a quantitatively selected region of interest, and processed with ImageJ-ImageDP; and by the construction of time series and analysis with either statistical diagnostics tests or Autoregressive Integrated Moving Average (ARIMA) models. Antimicrobial susceptibility tests are also performed for the purpose of quantitative comparison, showing a profile that is comparable to the result obtained with ImageJ-ImageDP processing after 15min of antibiotic action. Only the time series of the least affected bacteria (low Ciprofloxacin concentration) behaves in an expected manner, being non-independent and mainly non-linear, non-normal, and heteroscedastic. The most affected bacteria (higher Ciprofloxacin concentration) are non-independent and tend to be linear, normal and heteroscedastic. Adjustment to a linear regression identifies both, the culture medium without bacteria and the most affected bacteria, normality identifies the most affected bacteria and heteroscedasticity-homoscedasticity distinguishes the presence-absence of bacteria, respectively. ARIMA models (1,1,1)(1,0,1)_11_ and (4,1,1)(1,1,1)_11_ fit the time series of the most affected bacteria while the latter also fits the culture medium without bacteria. The time series of the least affected bacteria are identified by a (7,1,2)(1,0,1)_11_ model. The non-linear, non-normal and heteroscedastic behavior of this group is probably responsible for its adjustment to a model with a relatively high parameter. The four methods: diagnostic statistical tests, fitting of ARIMA models, ImageJ-ImageDP and antimicrobial susceptibility tests, show similar results, being able to distinguish among the groups of assays with bacteria and Ciprofloxacin below and above the Minimal Inhibitory Concentration.

**Author Summary:** Biospeckle laser patterns occur when a dynamic surface is illuminated. This research describes its application to the activity of *Escherichia coli* bacteria and the effect of 18 different concentrations of an antibiotic (Ciprofloxacin) in a 15min assay, using VDRL plates where the sample has a relatively small volume and is flat shaped. The assay is performed on an anti-vibration table in a dark room with a laser that sequentially illuminates each of the wells of the plate. A camera takes short 30sec videos with approximately 750 frames and sends them to a computer where image processing takes place. In order to select a segment of 80 successive frames to analyze, the region with the higher variation was identified, punched out and edited as a “flip-book animation” with a program named ImageJ and processed with another program named ImageDP that takes the difference between successive frames and is able to describe the speed with which the Biospeckle dots move, expecting to show that the antibiotic affects the bacteria by changing the speed with which they move. Also in each video, within the region of higher differential activity, a pixel was selected to construct a time series which is the successive value of that pixel in 253 frames, representing a recording of 8sec. This was analyzed with two statistical methods: diagnostic statistics tests and ARIMA models, both of which try to demonstrate how the results are organized. All the results, the speed of the dots in the “flip-book animation” and the structure of the data of the time series, were comparable to those obtained with traditional antimicrobial susceptibility tests with the same bacteria and antibiotic.

## Introduction

Antibiotic susceptibility testing is a necessary step prior to the treatment of clinical infections. A major concern is the time required to obtain a fast and reliable result especially in the case of acute bacterial infections for which it is necessary to determine the appropriate antibiotic that must be used. Established assays such as broth and agar dilution, and disk diffusion are the accepted techniques [1] but new approaches are needed in order to reduce the required time and the sample volume. Another issue is to search for the possibility of testing direct clinical samples reducing the assay requirements for handling and treatment of the microbial contaminated fluid as well as bacterial isolation and enrichment. Fast antimicrobial susceptibility testing has been achieved by diverse methods, among these are the molecular techniques [2,3] biosensors [4] fluorescence detection [5], as well as the adaptation of new techniques such as Raman Spectroscopy [6]. Therefore it is desirable to perform an exploratory assay which could eventually set the basis for a fast antibiogram.

Biospeckle Laser is a non-invasive technique and refers to a pattern that occurs when a laser beam illuminates a dynamic surface [7], such as a microorganism contaminated sample which has a movement of its surface that causes the wave fronts to interfere and produce a pattern of moving dots [8]. This technique has been used in different fields of research as well as at macroscopic and microscopic levels for a variety of applications [9, 10,11]. Usually this dynamic phenomenon is caught in a video for image processing. Diverse approaches and descriptors have been proposed that enable the information of the videos to be transformed in a number that represents the activity of the sample [12]. However, this is a multivariate phenomenon that is affected by the size of the organism to be detected, the speed and the direction of its movement [13], the type of medium or environment in which the organism is contained, etc.

The aim of the present work is to adapt the Biospeckle laser technique to antimicrobial susceptibility testing for Ciprofloxacin on *Escherichia coli* K-12. Ciprofloxacin is a broad-spectrum antimicrobial carboxyfluoroquinoline which selectively poisons preferentially the GyrA subunit of gyrase in Gram-negative bacteria and the ParC subunit of Topo IV in Gram-positive bacteria, with generation of chromosome breaks, which culminate in cell death [14,15]. An assay using VDRL plates that had already been used for testing Benznidazol on *Trypanosoma cruzi* [16], was adapted to *E*. *coli*. However, due to the type of microorganism, its size, motility and cell density, the processing of the images was performed on a quantitatively selected region of interest (ROI) which was analyzed through different approaches, such as video edition-processing and descriptive statistics; and time series construction and analysis with either statistical diagnostics tests or ARIMA models. A time series data set consists of observations on a variable or several variables over time which can be used for repeated-measures analysis or model adjustment [17]. Interpretation of Biospeckle images by time series analysis has already been proposed [18]. The selection of a ROI resulted in an important step because it allowed the consideration of different activity-level regions, as previously reported [19,20]. In this work a low Biospeckle intensity (*I*) region with a high differential bacterial activity was identified which was suitable for further analysis. Antimicrobial susceptibility tests were also performed for the purpose of quantitative and statistic comparison.

## Materials and Methods

### Antibiotic, Bacteria and culture conditions

*Escherichia coli* K-12 (*HfrH b*_*1*_^*-*^) [21] designated as strain 322 in the Laboratory Collection, was used in all the assays. For the Biospeckle and microbiological techniques, bacteria were grown overnight in Mueller-Hinton Broth (MHB) Himedia and prior to use it was adjusted to 0.5 McFarland standard which corresponds to approximately 1.5E+08 bacteria/mL (Absorbance 0.08-0.1 at λ_625nm_ in a Spectronic 20) [22]. The antibiotic used was Ciprofloxacin (BACIPRO^TM^ BIOGALENIC), i.v. infusion, 200mg/100mL. The leaflet indicates that it contains NaCl 0.9% (354 mg Na/100mL). This antibiotic was used in two-fold serial dilutions in sterile NaCl 0.9%.

### VDRL Plate Assay design for Biospeckle laser

The assay was designed as previously described [16]. Ciprofloxacin was serially diluted in saline (0.9% sterile NaCl). 95 μL of either Mueller-Hinton Broth (18 odd numbered wells) or bacterial suspension in Mueller-Hinton Broth (18 even numbered wells) were distributed in an alternating manner in wells of VDRL plates. At t=0, five μL of the antibiotic dilution were added to two contiguous wells, one without bacteria and one with bacteria, so that only the even numbered wells contained bacteria and both contained antibiotic at the same concentration. The plates were left still to avoid vibrations and to allow 15 minutes for the action of the drug at room temperature (20-25°C). Also 4 control assays were performed.

Table 1 shows the composition of each well. After 15 minutes, each well of the VDRL plate was processed under the experimental setup to obtain a video for each well. Four control wells (videos) were performed: Control A (95μL MHB + 5μL NaCl 0.9%); Control B (95μL with 1.43E+07 bacteria in MHB + 5μL NaCl 0.9%); Control C (100μL MHB); Control D (100μL with 1.50E+07 bacteria in MHB). The dimensions of the VDRL Plates were as specified by the supplier and previously reported; the mean internal diameter of each well was 12.7+/-0.1 mm and the height of 100μL in the well, was 0.75mm [16]. As it is described further on, the laser beam has a diameter of 10mm on the surface. With these dimensions the calculated volume of the illuminated liquid is 60μL and the number of bacteria present in this volume is approximately 8.6E+06, which is an appropriate number for evaluating the activity of the Biospeckle pattern.

**Table 1.**
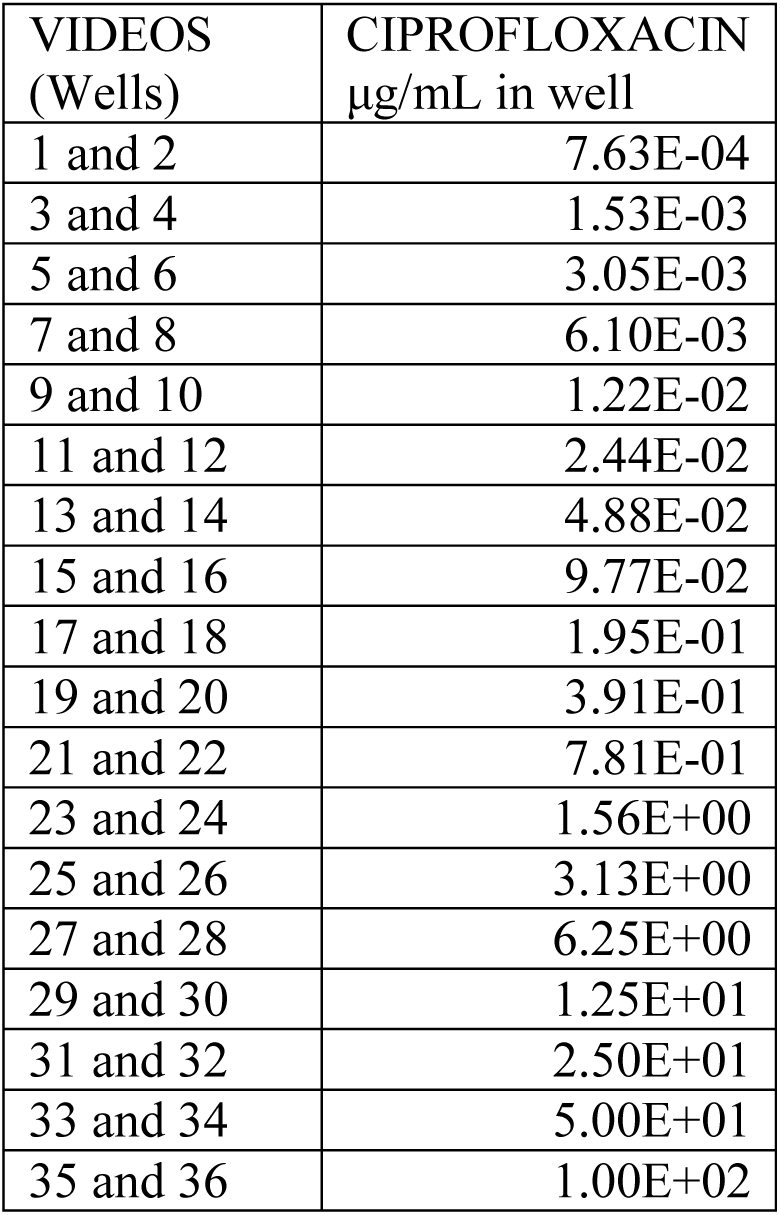
Concentration of Ciprofloxacin in the wells of the VDRL plates for Biospeckle laser. Odd numbered videos contained 95μL of MHB and 5μL of the antibiotic. Even numbered wells contained 95μL of bacteria (approximately 1.43E+07 total cells) in MHB and 5μL of the antibiotic. The same number was assigned to the well and to the video.

| VIDEOS<br>(Wells) | CIPROFLOXACIN<br>$\mu\text{g/mL}$ in well |
| --- | --- |
| 1 and 2 | 7.63E-04 |
| 3 and 4 | 1.53E-03 |
| 5 and 6 | 3.05E-03 |
| 7 and 8 | 6.10E-03 |
| 9 and 10 | 1.22E-02 |
| 11 and 12 | 2.44E-02 |
| 13 and 14 | 4.88E-02 |
| 15 and 16 | 9.77E-02 |
| 17 and 18 | 1.95E-01 |
| 19 and 20 | 3.91E-01 |
| 21 and 22 | 7.81E-01 |
| 23 and 24 | 1.56E+00 |
| 25 and 26 | 3.13E+00 |
| 27 and 28 | 6.25E+00 |
| 29 and 30 | 1.25E+01 |
| 31 and 32 | 2.50E+01 |
| 33 and 34 | 5.00E+01 |
| 35 and 36 | 1.00E+02 |

### Experimental setup

The experimental setup was the same as previously described [16] and is shown in Fig 1. Briefly, the setup rests on top of an anti-vibration table. The Biospeckle laser imaging system has a 1mW He-Ne unpolarized laser operating at 632.8nm. The laser which is located at a distance of 50cm is coupled to a convex divergent lens and the beam forms a 10mm spot in the individual wells of the VDRL plate, with an angle of incidence of 72°. Since the wells have an internal diameter of 12.7+/− 0.1mm, the beam illuminates the center of the well of the VDRL plate which rests on an opaque black cardboard, avoiding sparkles and edge effects. A CCD camera (Thorlabs USB.2, 30fps) is located at 30cm from the sample and is connected to a PC which records 0.5-1min videos. The resolution of the camera is 320 × 240, pixel size 6.45μm, speckle size aprox. 21.6μm. The speckle video data is sent to the computer for video and image processing. In order to visualize a significant number of bacteria per pixel, a relatively low resolution was chosen for the camera. Thus it was possible to apply statistical diagnostic tests and models on the time series of a selected pixel and descriptive statistics on the region of interest (ROI).

**Fig 1.**
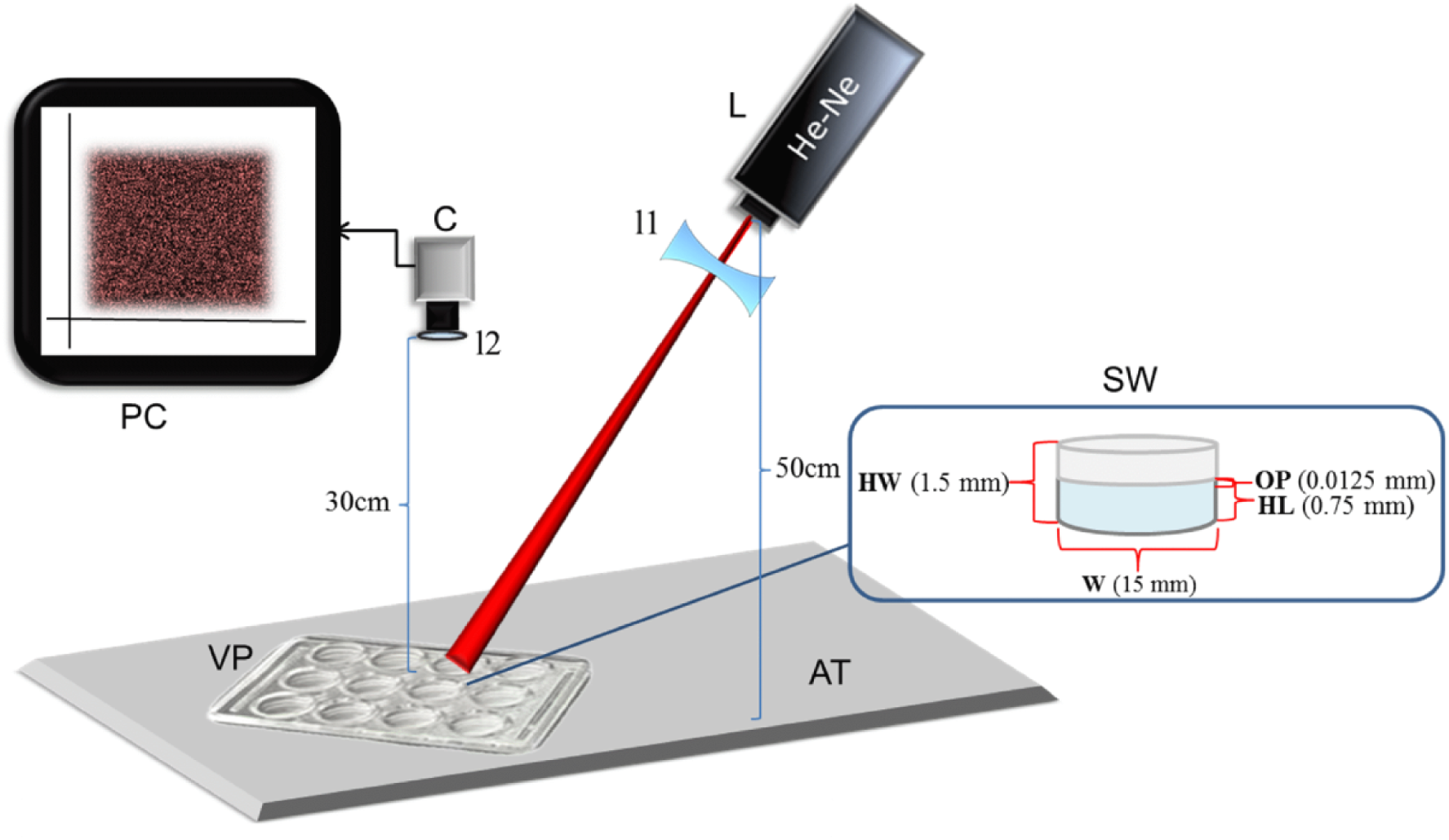
Schematic diagram of the speckle experimental setup. Experimental setup, with data of the setup and the VDRL well, L laser; C CCD camera; PC computer; l1 lens of the laser; l2 lens of the camera; AT anti-vibration table; VP VDRL plate; SW sample well; OP object’s plane;W width; HL height of the liquid; HW height of the well. The diagram is not drawn to scale. [16].

### Acquisition and handling of image data

#### Videos

40 videos were obtained. For each video in gray scale, the frames were downloaded using a frame-by-frame mode in the image processing unit. Videos contained over 750 frames (a recording of 30-60 seconds), each one with dimensions x=320, y=240 pixels, and each frame having an exposure time of approximately 0.033 sec. Regardless of the length of the video, the processed frames were always the same for each experiment and were taken starting on frame 100 which corresponds to 3.33 sec after the beginning of the video for repeated measures and time series and at frame 500 which corresponds to 16.5 sec, for video edition and processing. There are 4 control videos without antibiotic and 36 videos, each one having a dilution of the antibiotic, 18 of them are assays without bacteria and 18 of them are assays with bacteria. Although each video was processed individually, the results are sometimes combined in groups of low, intermediate and high antibiotic concentration. The Minimal Inhibitory Concentration (MIC) is ≤1 μg/mL according to CLSI [1], and 0.488 μg/mL as obtained in this work, which is close to the concentration of videos 19 and 20 and is usually included in the intermediate concentration group so as to differentiate the antibiotic effect of the high and low concentration groups.

#### Selection of the Region of Interest (ROI)

Video 1, which was taken from the well that contained culture medium with the smallest amount of added antibiotic, without bacteria, and Video 2 which was taken from the well that contained culture medium, bacteria and the same amount of added antibiotic, were used to construct a spatial-temporal speckle matrix that would be a guide to find the region with the maximal differential activity and would be selected as the region of interest (ROI). In each video, using the program ImageJ and starting with the raster images, each one being a systematic division of the illuminated space, a segment of 11 lines was taken (*y*=115 to *y*=125, both inclusive) from 11 successive frames which include a temporal resolution of 0.37 seconds (from 3.33sec to 3.70 sec of each video). Each line spanned from *x*=0 to *x*=319. Since this was performed in 11 successive speckle images (frames 100-110), for each “*x*”, there were 121 values (11 “*y*” values from 11 frames). These 121 values were used to obtain the Standard Deviation (SD) for each one of the 320 “*x*” values of Video 1 and of Video 2. Fig 2 shows a diagram of the selection of the ROI. Then, the 320 SD values of each video were used to obtain 320 values of ***Qx*_*i*_**, as follows:

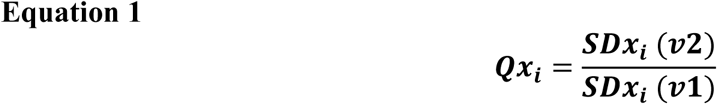

**Fig 2.**
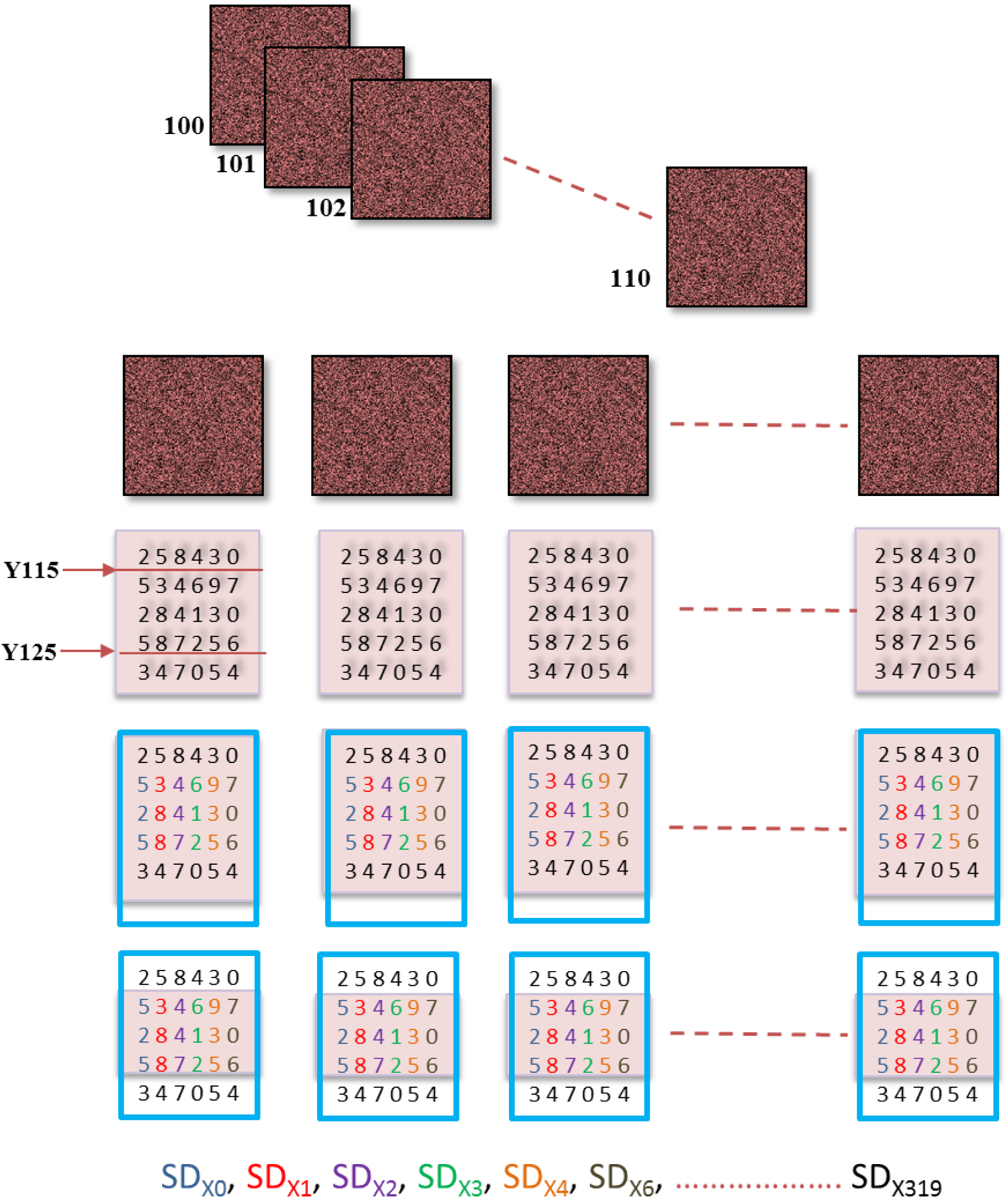
Selection of the region of interest (ROI). Schematic representation of the quantitative selection of the ROI.

Where ***SDx*_*i*_** is the Standard Deviation in each “*x”, i* spans from 0 to 319, and *v1* and *v2* refer to Video 1 and Video 2, respectively. This yields 320 values for ***Qx*_*i*_**, each one derived from a spatial temporal speckle matrix of 121 values from each video.

The region with the highest values for ***Qx*_*i*_**, was further examined in order to select the ROI.

#### Image-ROI video edition and processing by the method of Generalized Differences

In each of the 40 videos, the region of interest (ROI) was extracted from 80 successive frames. A region with coordinates *x*282-*x*311; *y*105-*y*135, was extracted which resulted in a square matrix with 30×30 pixels, from the 80 frames. Then, using the program ImageJ (a public domain Java image processing program developed at NIH, USA), a new video was constructed with the 80 selected regions, generating an uncompressed digital “flip book animation” (Fig 3), at a frame rate of 7fps. Each new edited video was processed with ImageDP, according to previous studies [16], obtaining a value of *A* which is a measure of the Biospeckle activity of the sample, as captured in each video. Briefly, Eq (2) was developed as an algorithm of averages that analyzed all the frames in one video sequence. *A* is a measure of mean intensity (*I*), being expressed as an averaged mean intensity by the method of Generalized Differences:

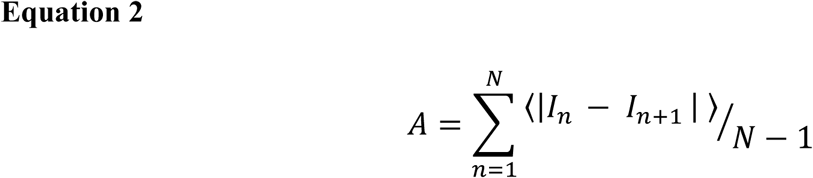

where |.| is the absolute value, <.> is the mean value, *I* is intensity in gray scale and N represents the total number of frames analyzed, which in this case is 80. The mean intensity of the absolute value of a subtraction pixel by pixel of whole successive images is averaged to obtain *A*, when a video has 80 images, there will be 79 mean differences (N-1). With this algorithm, successive images that were constructed with a square matrix of 30×30 pixels are subtracted, transformed to the absolute value of the difference, the mean for the absolute values of the intensities of the differences is obtained and finally the means are averaged. It should be noted that with this algorithm, the final number corresponds to the average of means. In the case of Eq (2), this was achieved with a free software program, Image Delta Processor (named as ImageDP), written for this purpose in Java. ImageDP is available on line at http://bigjocker.com/biospeckle/ and it can be used according to the instructions of the ImageDP Manual (S1 Text). Thus for each assay, a video was constructed with ImageJ and processed with ImageDP. The 40 values of *A* obtained were used to fit a linear regression and also combined in groups of low, intermediate and high antibiotic content as described above.

**Fig 3.**
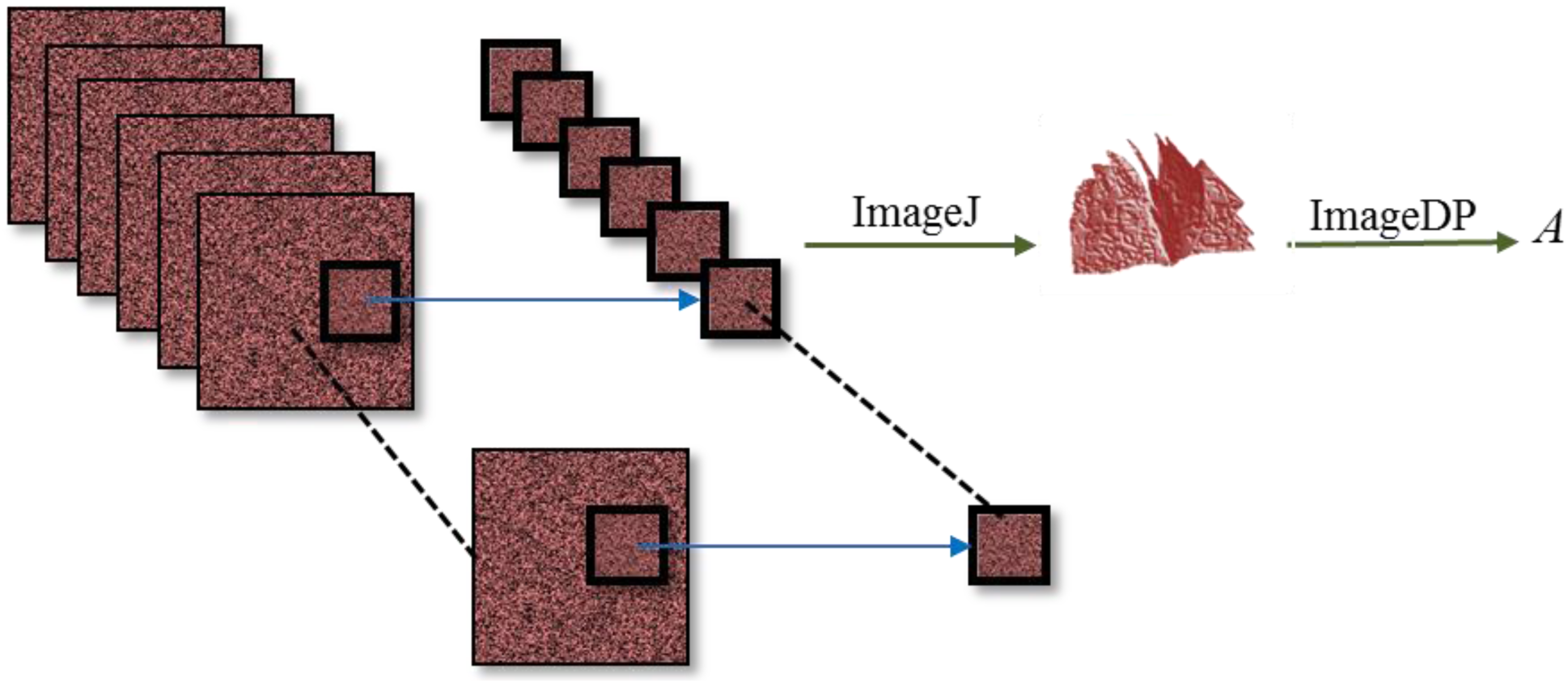
Image-ROI video edition and processing. . Schematic representation of the selection and punching out of a ROI, construction of a new video with ImageJ and processing with ImageDP.

#### Selection of the pixel for the time series

The region with the highest values for ***Qx*_*i*_**, was further examined in order to select a value of “*x”*, which would define one of the coordinates for the pixel to be selected. The other coordinate was chosen as *y*=120 because it represents the middle of the *y*-scale of the frame and also of the constructed matrix (*y*=115 to *y*=125). The chosen pixel was (*x*=303, *y*=120) and it was used to develop the time series for each of the 40 videos. In each one of the 40 videos, this pixel lays within the region with the least intensity (*I*), making it possible to detect the ***Qx*_*303*_**, as one of the highest among all the ***Qx*_*i*_** values.

#### Construction of the time series

SAGA (System for Automated Geoscientific Analysis) was used for the construction of a time series of each of the 40 videos. The “Shapes” Module was used (Shapes points, Convert table to points and Shapes Grid, Add grid values to points) of the Coordinate (*x*303,*y*120). Each time series had a length of 253 frames which corresponded with a time period of approximately 8 seconds. The time series were processed individually with the statistics tools.

### Statistics tools applied on the time series for repeated-measures analysis

#### Normality test: Shapiro Wilk

For the Shapiro Wilk Normality test, the H_0_ states that the population is normally distributed [23]. If the p-value is less than 0.05, the H_0_ is rejected and there is evidence that the data are not from a normally distributed population, accepting the alternative hypothesis (H_1_). This test was performed in R and R commander. Since this test is sensitive to the number of values it was performed with the first 50 values of the time series.

#### Linear Regression

In this case the H_0_ states that there is no significant linear correlation between the dependent and the independent variable, p-value greater than 0.05. If the p-value is less than 0.05, H_0_ is rejected; hence there is a significant relationship between the variables in the linear regression model. However, linear regressions are based on assumptions of homoscedasticity and independence that will be tested below. This test was performed in R and R commander.

#### Homoscedasticity / Heteroscedasticity: Breusch-Pagan

In this test the H_0_ states that there is homoscedasticity, p-value greater than 0.05; therefore, a p-value less than 0.05 is indicative of heteroscedasticity [24]. The test was performed in R and R commander.

#### Independence / Autocorrelation: Durbin-Watson test

The H_0_ (p-value greater than 0.05) states that the errors are serially uncorrelated against the H_1_ (p-values less than 0.05) that they follow a first order autoregressive process. Furthermore, for the D statistic “small” values or values close to 0, are indicative of positive correlation [25,26]. This test was performed in R and R commander.

#### Autocorrelation: ACF and PACF

For a given time series, the ACF (Autocorrelation Function), the PACF (Partial Autocorrelation Function) as well as the run sequence plot of the data as a function of time, were obtained [27]. This test was performed in R and R commander.

#### Auto Regressive Integrated Moving Average (ARIMA) Models

The (*p,d,q*)(P,D,Q)_*m*_ terms of the ARIMA models were selected according to the Box and Jenkins methodology [27]. Briefly, for a given time series, the run sequence plot is examined to see if it is stationary and if it has trend and seasonality. The ACF and PACF are examined to evaluate if the stochastic process is random and independent (white noise). If the process is not white noise, the ACF and PACF functions are evaluated in order to identify the type of temporal autocorrelation, as follows: the ACF is examined to see if the data is random, if there is autocorrelation and auto regression and if it does not represent white noise, also the shape of the decay of the ACF is indicative of the type of model that should be chosen; the PACF is examined to see the lags that show statistical significance. The first-difference trend and seasonal transformations are performed in order to obtain second order stationarity. Usually for a time series that has trend and seasonality the first-difference run sequence plot will have a mean of the differenced data around zero. Also, the significant lags of the PACF of the original data will indicate the “AR” (*p*) term while the significant lags of the ACF of the differenced data will indicate the “MA” (*q*) term. Trend was included starting with the lowest possible value and seasonality (P,D,Q) _*m*_ was selected by a quantitative approach, with the sub index term “_*m*_” (the number of periods in each season) being 11. Although each video was processed individually, the tendency of the results is sometimes analyzed in groups as described above in the *Videos* epigraph and in reference to the MIC.

### Antimicrobial susceptibility testing

Antimicrobial testing methods were performed in order to confirm the susceptibility of the strain to the concentration range of Ciprofloxacin, under the conditions used. The MIC value was recorded as the lowest concentration of the assayed antimicrobial agent that inhibits the growth of the microorganism.

#### Broth Macro-dilution

This test was prepared in tubes with 1mL MHB containing serial two-fold dilutions of Ciprofloxacin and 1mL of MHB containing bacteria at 0.5 Mc Farland Standard. The microorganism was allowed to grow for 24 hours at 36°C then the Absorbance at 625nm was taken.

#### Agar dilution spot test

The bacterial strain was grown in MHB up to 0.5 Mc Farland Standard and plated on the surface of Mueller-Hinton Agar (MHA), Becton, Dickinson and Company. Spots of 5μL serial two-fold dilutions of Ciprofloxacin were placed on the surface [28]. The plates were incubated for 24 hours at 36°C and inhibition of growth was evaluated, measuring the diameter of the halo under the magnifying microscope.

## RESULTS

The present paper aims at applying the Biospeckle methodology already designed [16] to a bacterial culture in the wells of VDRL plates, with the purpose of performing an exploratory assay which could eventually set the basis for a fast antibiogram.

### Selection of the Region of Interest (ROI)

Developing the Biospeckle methodology with bacteria has to take into account the size and the density of the microorganism, as well as the motility and Brownian motion. Moreover, the assumption of homogeneity cannot be applied directly to Biospeckle laser monitoring of microorganisms in a medium [29]. Since there are variations of homogeneity in the illuminated sample, particularly in a culture medium with cells, it was necessary to select a region of the frames where the activity of the microorganism and the effect of the antibiotic are properly revealed. It should be noted that the laser illuminated the plate with an angle of incidence of 72° in order to obtain different intensity regions within each image. The same illumination was applied to all the samples.

A line at the middle of the frame (*y*120) was selected for all the values of “*x*” (*x*0-*x*319). Fig 4a and Fig 4b show the intensity (*I*) for all the pixels in that line, for frame “100”, taken from both, Video 1 (culture medium with the lowest added antibiotic concentration, without bacteria) and Video 2 (culture medium with the lowest added antibiotic concentration, with bacteria) respectively, with both showing a similar profile and dispersion pattern, the double arrow points at the region *x*282-*x*311 at the middle of frame “100” (*y*120), which was selected as a high activity region as is shown later. The usual Biospeckle practice is to take the region with the mean intensity for analysis. However, as Fig 4a and Fig 4b show, in this case the mean intensity region also has the highest slope, implying that it is a region with a high variation. Nevertheless, all the regions (from pixel *x*=0 to pixel *x*=319) were evaluated in order to find the pixels where the presence of bacteria was best revealed. Fig 5a shows the surface plot for the segment *x*0-*x*319;*y*115-*y*125 of Video 1 (without bacteria) and Fig 5b shows the same area of Video 2 (with bacteria).

**Fig 4.**
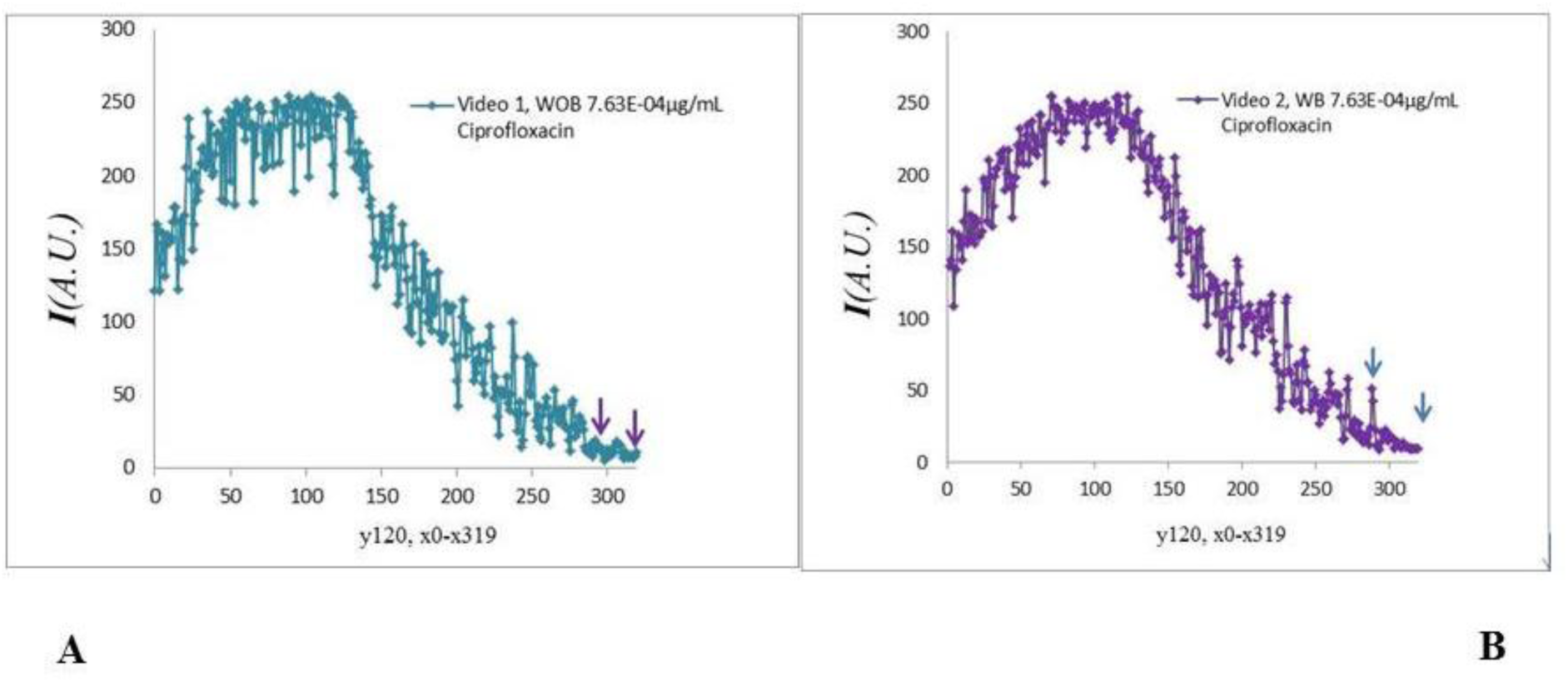
Intensity (*I*) of pixels at line *y*120 of Videos 1 and 2. Intensity of pixels at line *x*0-319 *y*120 of A. Video 1 which was taken from an assay that contained culture medium (MHB) without bacteria (WOB) and the least amount of Ciprofloxacin (7.63E-04 μg/mL in the well), and B. Video 2 which was taken from an assay that contained culture medium (MHB) with bacteria (WB) and the least amount of Ciprofloxacin (7.63E-04 μg/mL in the well).

**Fig 5.**
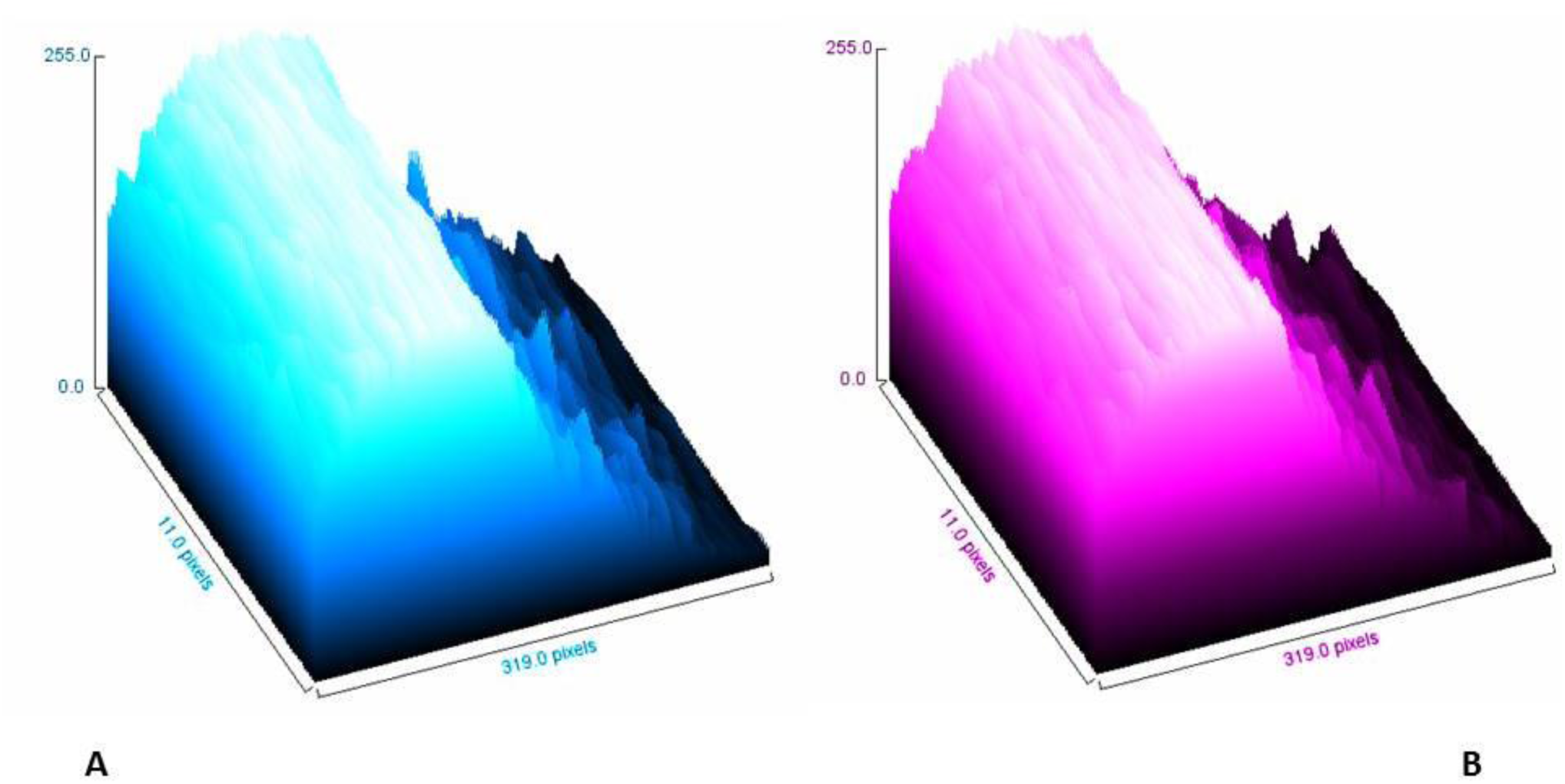
Surface plot of a segment of a Video. Surface plot of the segment *x*0-*x*319 *y*115-*y*125 of A. Video 1 (culture medium (MHB) without bacteria and the least amount of Ciprofloxacin, 7.63E-04 μg/mL in the well) and B. Video 2 (culture medium (MHB) with bacteria and the least amount of Ciprofloxacin, 7.63E-04 μg/mL in the well).

For the selection of the ROI, the data of Video 1 and Video 2 were used applying Equation 1, as described in Materials and Methods (*Selection of the Region of Interest (ROI)* and Fig 2). Fig 6 shows the result of this spatial-temporal analysis in a plot with the x-axis crossing at a value of 1 in order to distinguish the areas where the SD of Video 2 (with bacteria) is higher than the SD of Video 1 (without bacteria). A region of higher differential Biospeckle activity with coordinates *x*282-*x*311;*y*115-*y*125 is shown between the arrows. The purpose of this spatial-temporal analysis was to select the ROI through a quantitative method, avoiding qualitative approaches.

**Fig 6.**
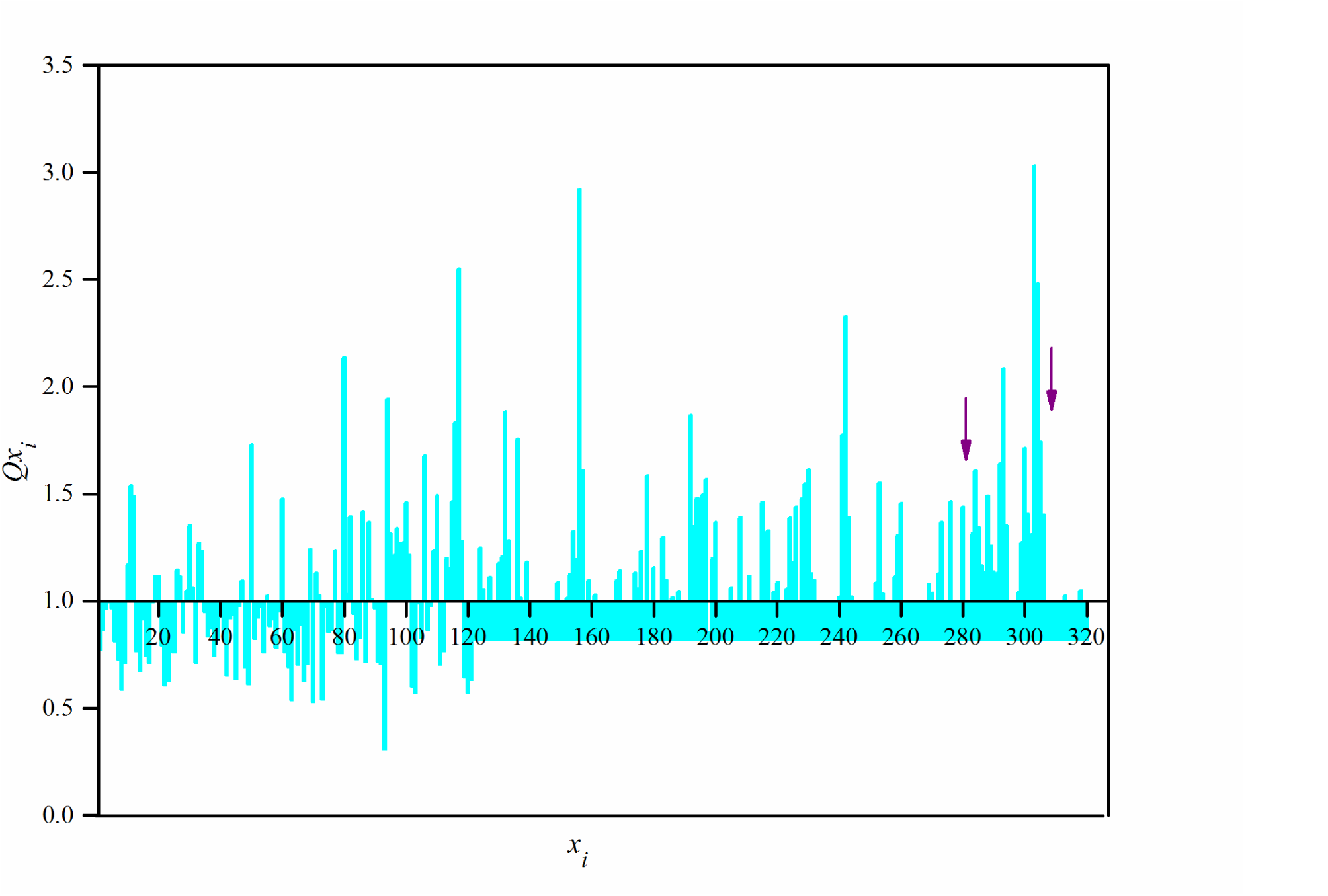
Spatial temporal analysis of the SD. Application of Equation 1 to the data of Video 1 (MHB without bacteria and Ciprofloxacin 7.63E-04 μg/mL) and Video 2 (MHB with bacteria and Ciprofloxacin 7.63E-04 μg/mL). The SD was taken for each column of pixels in “*x*” for *y*120-115 in 11 frames. Equation 1 was applied and the quotient of Video2/Video1 is plotted with the x-axis crossing at a value of 1 in order to distinguish the areas where the SD of Video 2 (with bacteria) is higher than the SD of Video 1 (without bacteria).

### Generalized differences as descriptor of Biospeckle activity of the edited videos

Having identified the area of high differential activity, a segment of 30×30 pixels (*x*282-*x*311; *y*105-135) was extracted from each video. Figs 7 and Fig 8 show the selected areas for Videos 1 and 2, respectively. It should be noted that although this selected area has the lowest intensity (*I*) value within the frame, it has a higher differential SD, suggesting that it is in this region that the presence of bacteria and the effect of the antibiotic could be detected. Hence in each case (Videos 1-36) a new edited video was constructed as shown in Fig 3, with the segment of maximal differential activity. For this purpose, the region of interest (ROI) was extracted from 80 successive frames, a video was constructed and each new video was processed with ImageDP, according to previous studies [16], obtaining a value of *A* which is a measure of the Biospeckle activity of the sample, as captured in the edited video. S1 Data shows the values of *A* for the videos with and without bacteria, however videos 31/32 to 35/36 were not included due to a distortion caused by the higher antibiotic concentrations. This type of behavior was also observed in [16]. Fig 9 shows the linear regression for the values of *A* taken from S1 Data, as a function of the logarithm of the concentration of Ciprofloxacine. Although the curve with bacteria shows a clear tendency to decrease as the antibiotic concentration rises, it also shows a low determination coefficient with a high dispersion of the points.

**Fig 7.**
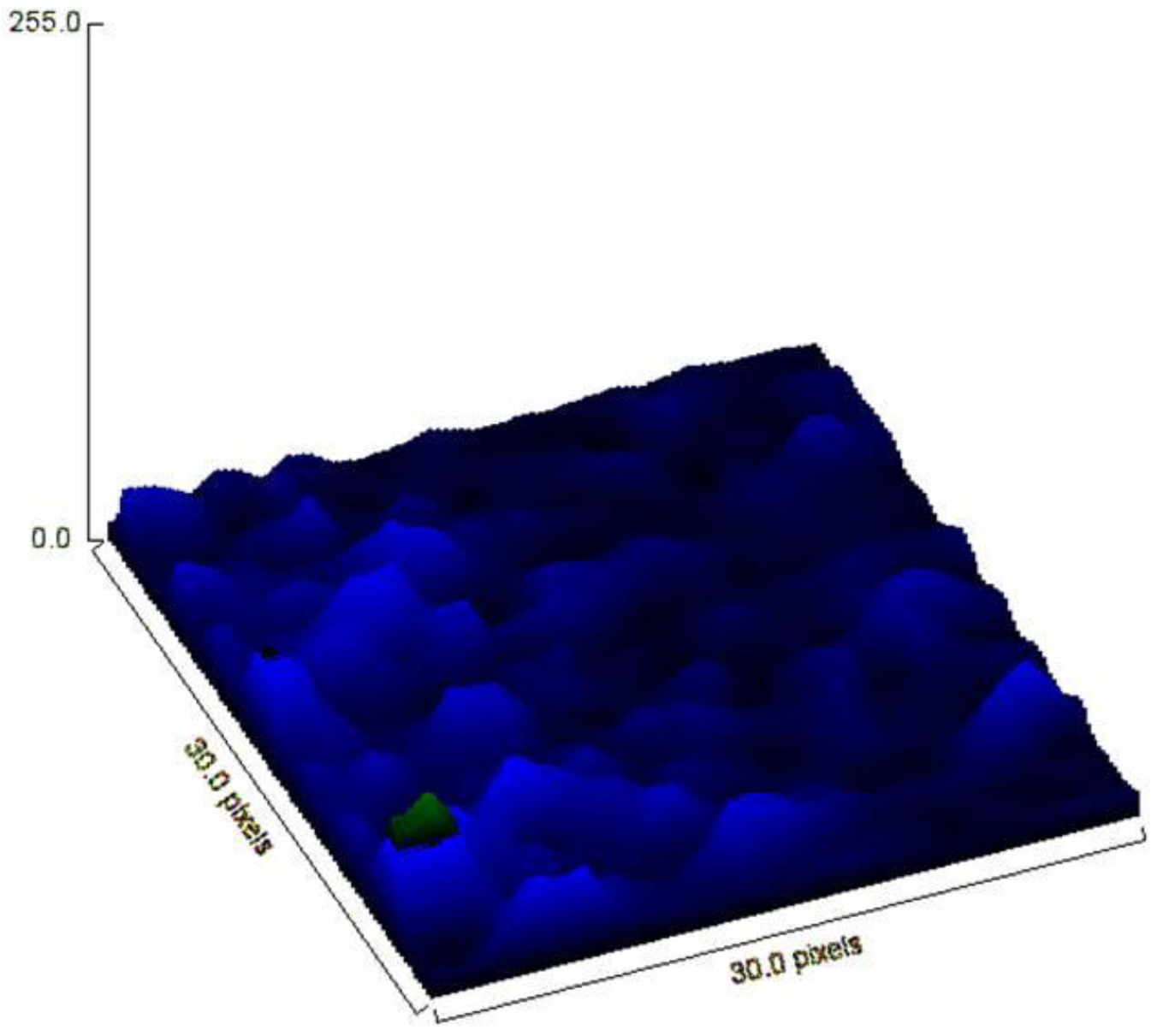
Surface plot of the selected ROI of Video 1. Surface plot of the segment *x*282*-x*311 *y*115*-y*125of Video 1 (culture medium (MHB) without bacteria and the least amount of Ciprofloxacin, 7.63E-04 μg/mL in the well).

**Fig 8.**
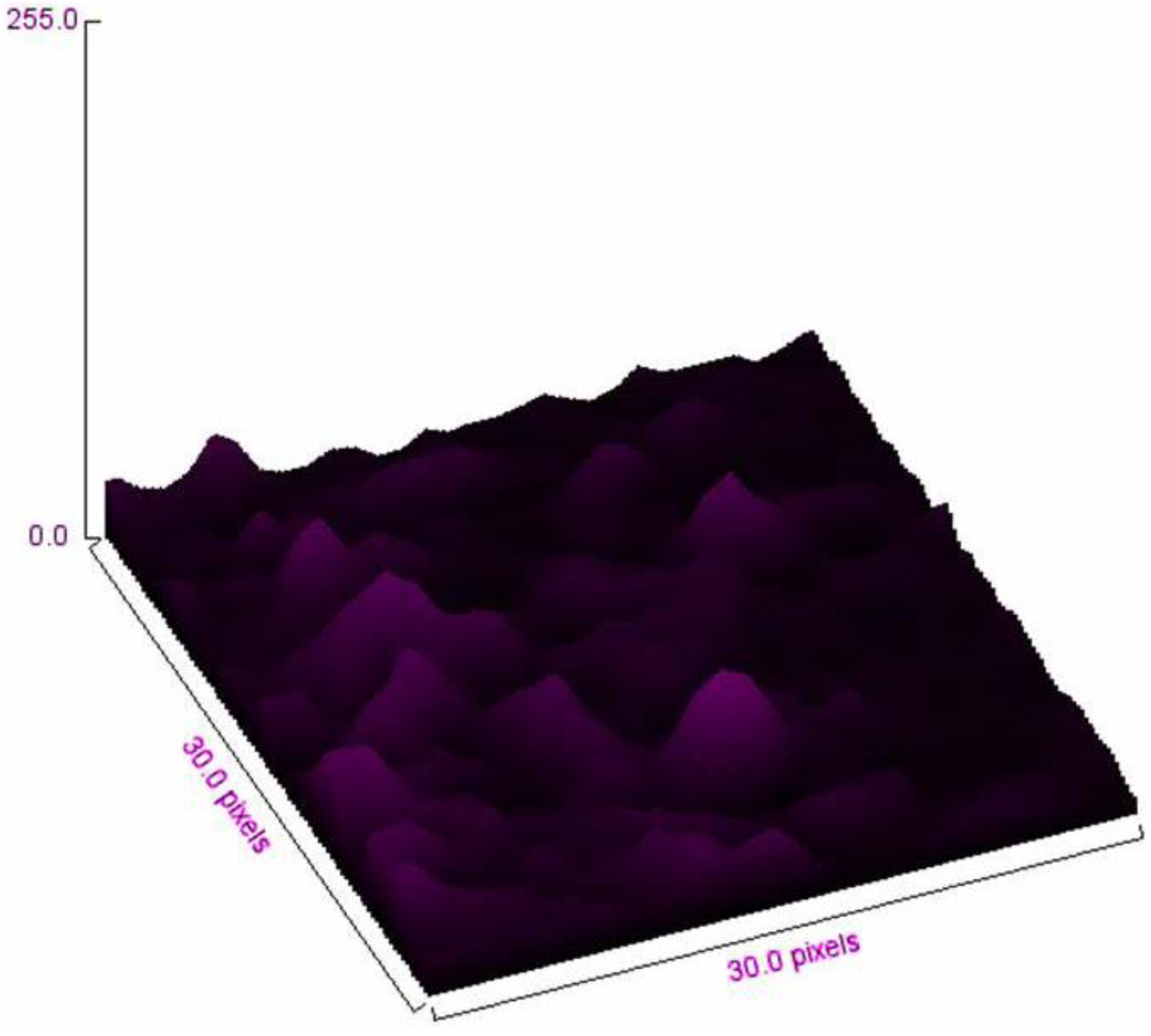
Surface plot of the selected ROI of Video 2. Surface plot of the segment *x*282-*x*311 *y*115-*y*125of Video 2 (culture medium (MHB) without bacteria and the least amount of Ciprofloxacin, 7.63E-04 μg/mL in the well).

**Fig 9.**
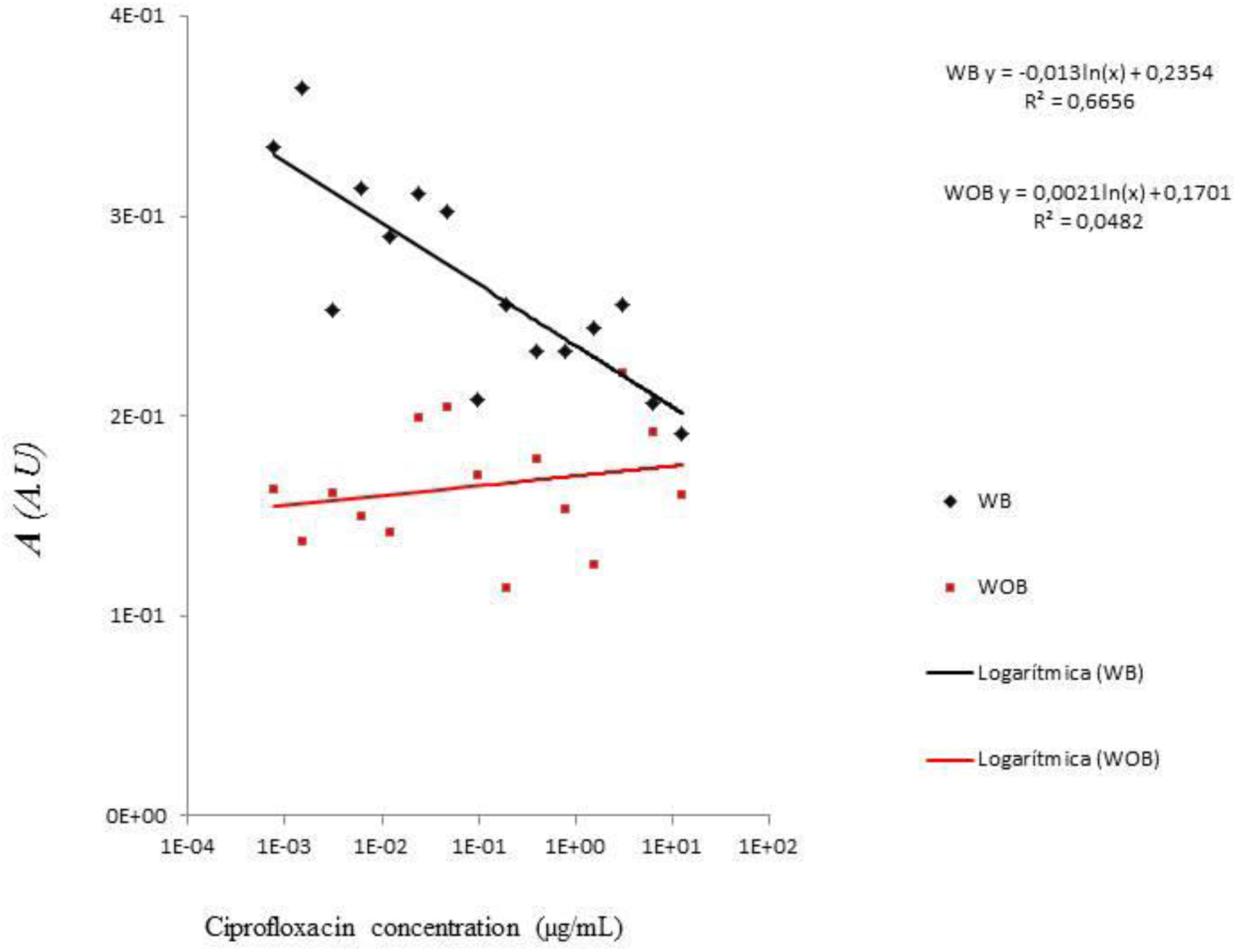
Linear regression for the values of *A* as a function of the logarithm of the concentration of Ciprofloxacin. WB, with bacteria; WOB without bacteria.

Due to the dispersion of the data, the videos with and without bacteria, were organized in three groups, low (7.6E-04 to 1.2E-02 µg/mL), intermediate (2.4E-02 to 3.9E-01 µg/mL) and high concentration (7.8E-01 to 1.25E+01 µg/mL) of Ciprofloxacin. It should be noted that for the purpose of comparison, all groups have the same number of videos (5 videos each group). However, within each group the range of the antibiotic concentration may have up to two orders of magnitude. According to CLSI, the MIC for Ciprofloxacin on *E*. *coli* is ≤1μg/mL [1]. In this work the MIC of Ciprofloxacin on the used strain was determined at 0.488 μg/mL by two methods, Broth Macrodilution and Agar Dilution as shown below. This is close to the antibiotic concentration of videos 19 and 20 (0.391 μg/mL), both included in the intermediate group. S1 Data contains the the data of the edited videos. Odd and even numbered videos 31 to 36 were excluded from this analysis because the high antibiotic concentration caused a distortion in the Biospeckle pattern. This effect had already been described when benznidazol was the test drug on *Trypanosoma cruzi* [16]. In each group, the Biospeckle activity was obtained with ImageDP as an *A* value, then the the mean and the SD of the groups, were calculated and are shown in Fig. 10. As Fig 10 shows, the Biospeckle activity of the videos without bacteria is similar in the three groups, with a slight tendency to increase at the higher antibiotic concentrations. In the group of videos with bacteria, the Biospeckle activity is higher at the lower antibiotic concentrations and drops as the concentration increases. The Biospeckle activity of the bacteria at the highest concentrations of antibiotic approaches the Biospeckle activity of the culture medium without bacteria and the same antibiotic concentrations.

**Fig 10.**
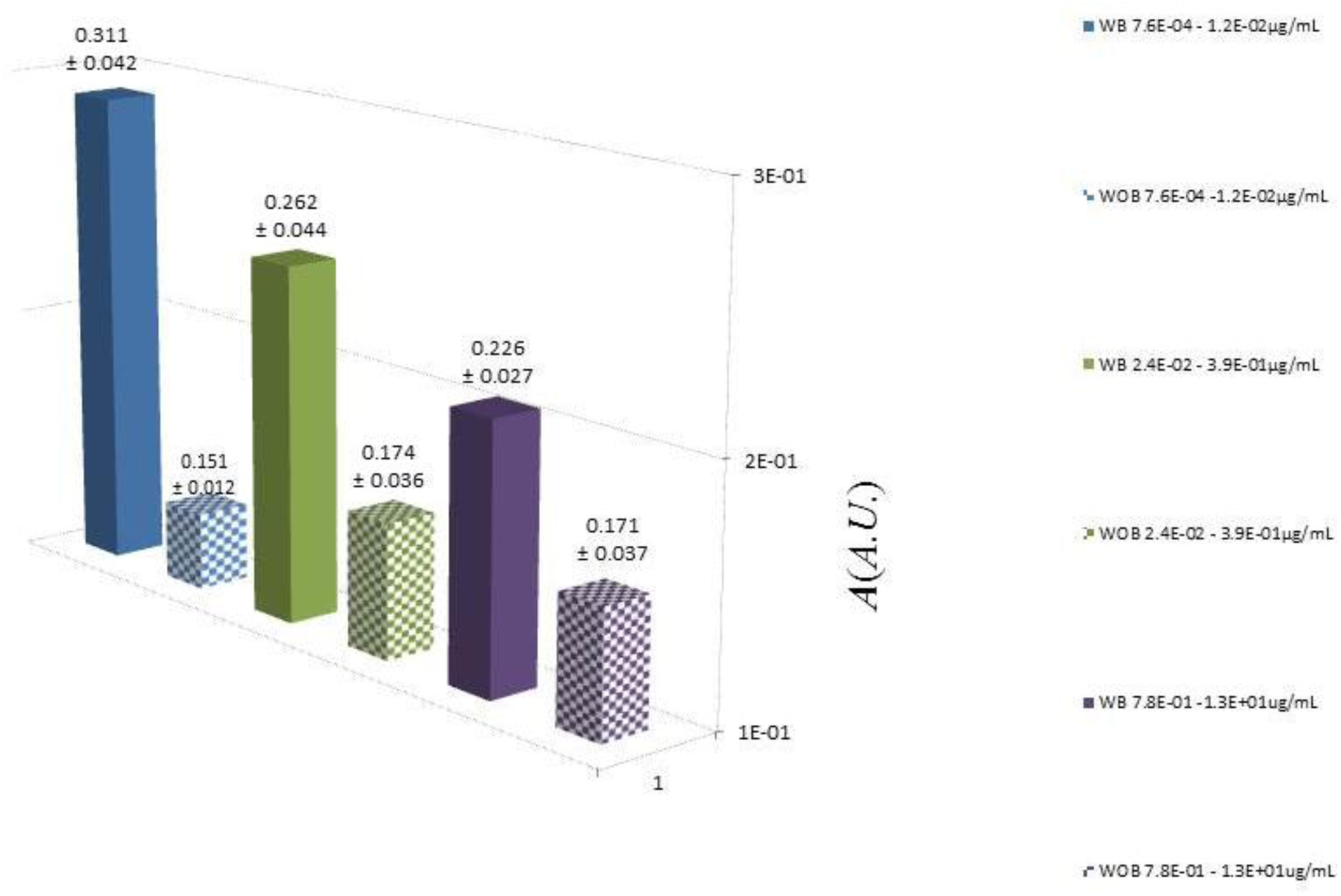
Activity of the Biospeckle pattern of the edited videos. Each video was edited by punching out the ROI and constructing a new video with ImageJ, which was processed with ImageDP, obtaining in each case, a value of *A* which represents a measure of the activity of the Biospeckle pattern. The results are organized in groups according to the concentration of Ciprofloxacin, as follows, low (7.6E-04 to 1.2E-02 µg/mL), intermediate (2.4E-02 to 3.9E-01 µg/mL) and high (7.8E-01 to 1.25E+01 µg/mL) concentration of Ciprofloxacin; WOB without bacteria, WB with bacteria.

All this indicates that, if the ROI is properly selected, and new videos are constructed with ImageJ and analyzed with ImageDP, it is possible to distinguish the videos with bacteria from those without bacteria and the effect of the antibiotic. Moreover, if the videos are analyzed in separate groups, above and below the MIC, they show a difference in the activity of the Biospeckle pattern with different values, when the antibiotic is present at concentrations above and below the MIC.

### Diagnostic statistical tests for repeated-measures analysis (linear regression model, normality, homoscedasticity and independence of the time series of each video)

In an attempt to improve the detection of the presence of bacteria and the effect of the antibiotic, a time series was extracted within the selected ROI of each of 38 videos (odd numbered videos 1-35; even numbered videos 2-36 and control videos A and B). The coordinates were chosen from a high differential activity pixel within the selected ROI. In all cases the time series was extracted at pixel *x*303;*y*120. Statistic tests were performed for each time series.

Figs 11-14 show the results of diagnostic statistical tests, for the time series of each video and two control videos (see S2 Data). In this case each test was performed on the individual time series and the results were considered in groups for the purpose of analysis and discussion.

**Fig 11.**
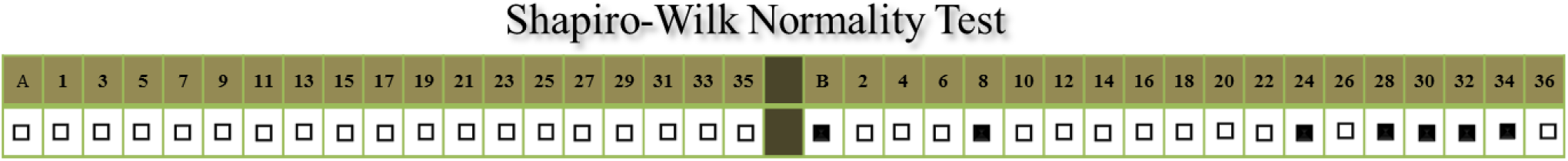
Shapiro-Wilk Normality Test. This test was performed on the time series of all of the videos with the values of pixel *x*303;*y*120 of the first 50 frames (frames 100-150) because it is sensitive to the number of input values. Note: black indicates that the H_0_ is accepted and there is normal distribution of the data; white means that the H_1_ is accepted and the data are not normally distributed.

**Fig 12.**
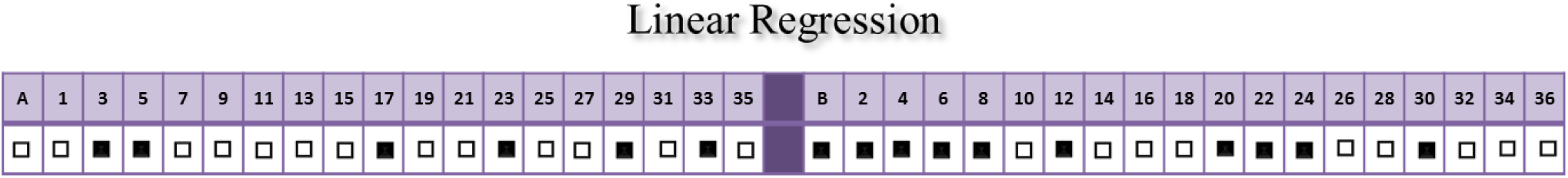
Adjustment to a linear regression. . This test was performed on the time series of all of the videos on the values of pixel *x*303;*y*120 over 253 frames (frames 100-353). Note: black indicates that the H_0_ is accepted and there is no relation among the variables; white means that the H_1_ is accepted and there is a linear relation among the dependent and independent variables.

**Fig 13.**
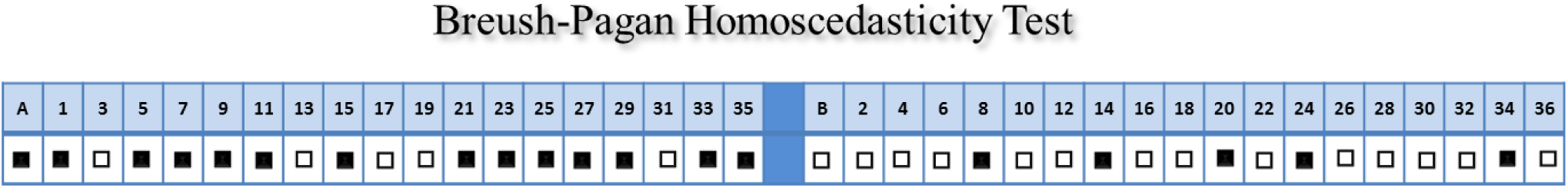
Breusch-Pagan homoscedasticity / heteroscedasticity test. . This test was performed on the time series of all of the videos on the values of pixel *x*303;*y*120 over 253 frames (frames 100-353). Note: black means that the H0 is accepted and there is homoscedasticity; white means the H_1_ is accepted and there is heteroscedasticity.

In the assays with culture medium, bacteria and low antibiotic concentrations (7.6E-04 to 2.4E-02, even numbered videos 2-12) below the MIC, the p-value for a linear regression is greater than 0.05 in 6 out of 7 assays, so there is not a significant relationship between the dependent and independent variables, in a linear regression model; there is non-normality (Shapiro-Wilk test) in 5 out of 7 assays, there is heteroscedasticity (according to the Breusch-Pagan test) in 6 out of 7 assays and positive autocorrelation (Durbin-Watson test) in all of them. Therefore, the presence of bacteria with the least antibiotic effect, promotes a Biospeckle pattern in the time series that has positive autocorrelation and is mainly heteroscedastic, non-linear with a non-normal distribution of the data. In this case, the non-linear regression is compatible with the heteroscedasticity and non-independence (autocorrelation).

In assays with culture medium, bacteria and the antibiotic that approaches the MIC from the lower concentrations (4.88E-02 to 7.81E-01, even numbered videos 14 to 22) and includes it, there is a mixed behavior which suggests a transient condition between the low and the high concentration groups, in the linear regression and homo/heteroscedasticity, with non-normal behavior and positive autocorrelation in all of the assays.

In assays with culture medium, bacteria and the antibiotic at concentrations above the MIC (1.56E+00 to 1E+02, even numbered videos 24 to 36), in 5 out of 7 time series, there is a significant relationship between the dependent and independent variables in a linear regression model, being similar to the culture medium without bacteria and with antibiotic and contrary to the low concentration group (bacteria with low antibiotic concentrations). Furthermore, this high concentration group shows mainly heteroscedasticity (5 out of 7) and a normal behavior (5 out of 7) with all of the time series having positive autocorrelation. This is an unexpected combination of results because the assumptions for a linear regression are not satisfied.

Considering all the assays with culture medium with different concentrations of Ciprofloxacin (7.6E-04 to 1.0E+02, odd numbered videos 1-35) without bacteria, in 13 out of 19 time series the p-value for a linear regression is less than 0.05 so there is a significant relationship between the dependent and independent variables. In 14 of the 19 videos, there is homoscedasticity according to the Breusch-Pagan test and in all of them there is a non-normal distribution as indicated in the Shapiro-Wilk test and positive autocorrelation with the Durbin-Watson test. Therefore, the time series of the culture medium without bacteria and different concentrations of the antibiotic have a Biospeckle pattern that have a non-normal distribution with positive autocorrelation and are mainly homoscedastic and linear. All of the time series that follow a linear regression are expected to be homoscedastic and independent according to the assumptions.

### Autocorrelation Function (ACF), Partial Autocorrelation Function (PACF) and Selection of ARIMA models

Since the results of the Durbin-Watson test (Fig 14) showed that all the time series were non-independent with positive autocorrelation, the ACF and PACF were performed for each one of the 40 time series, confirming the autocorrelation. As an example, the run sequence plot, ACF and PACF for the time series of Video 1 (culture medium with the lowest added antibiotic concentration, without bacteria) and Video 2 (culture medium with the lowest added antibiotic concentration, with bacteria), are shown in S3 Data. Model adjustment was individually made in each of the 40 videos but the discussion is performed considering the groups as above.

**Fig 14.**
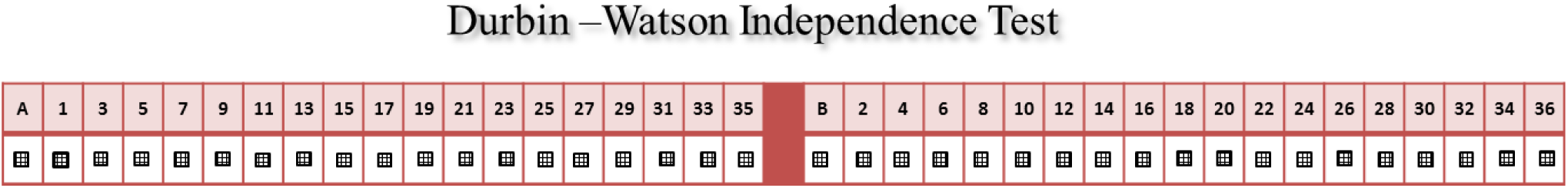
Durbin-Watson test for independence. . This test was performed on the time series of all of the videos on the values of pixel *x*303;*y*120 over 253 frames (frames 100-353). In all cases the p-values are less than 0.05, rejecting the H_0_ which states that the errors are serially uncorrelated and accepting the H_1_ which states that the errors follow a first order autoregressive process. Small values of the D statistic indicate that successive error terms are close in value to one another, or positively correlated. Note: gray means there is positive autocorrelation, which was found in all the time series.

Controls A (culture medium with NaCl 0.9%, without bacteria) and C (culture medium without bacteria and without NaCl) were used as an orientation for model selection. Once it was determined that there is autocorrelation, an ARIMA model was explored for the time series of pixel *x*303;*y*120 of Control A, according to Box-Jenkins approach. A process to obtain the best ARIMA (*p, d, q*)(P,D,Q)_m_ model was worked out as follows.

The time series was examined for Control A. The run sequence plot of the time series shows that it is non-stationary (Fig 15), the ACF (Fig 16) shows that the data does not represent white noise, the data are not random, the observations are related to adjacent observations and there is a strong autocorrelation and auto regression. The PACF function (Fig 17) shows statistical significance for lags 1 and 2.

**Fig 15.**
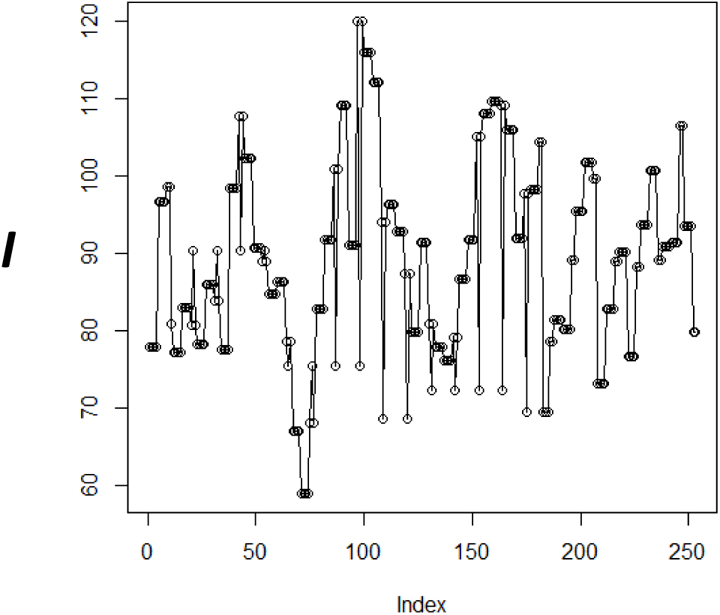
Run sequence plot of Control A. (culture medium with NaCl 0.9%, without bacteria).

**Fig 16.**
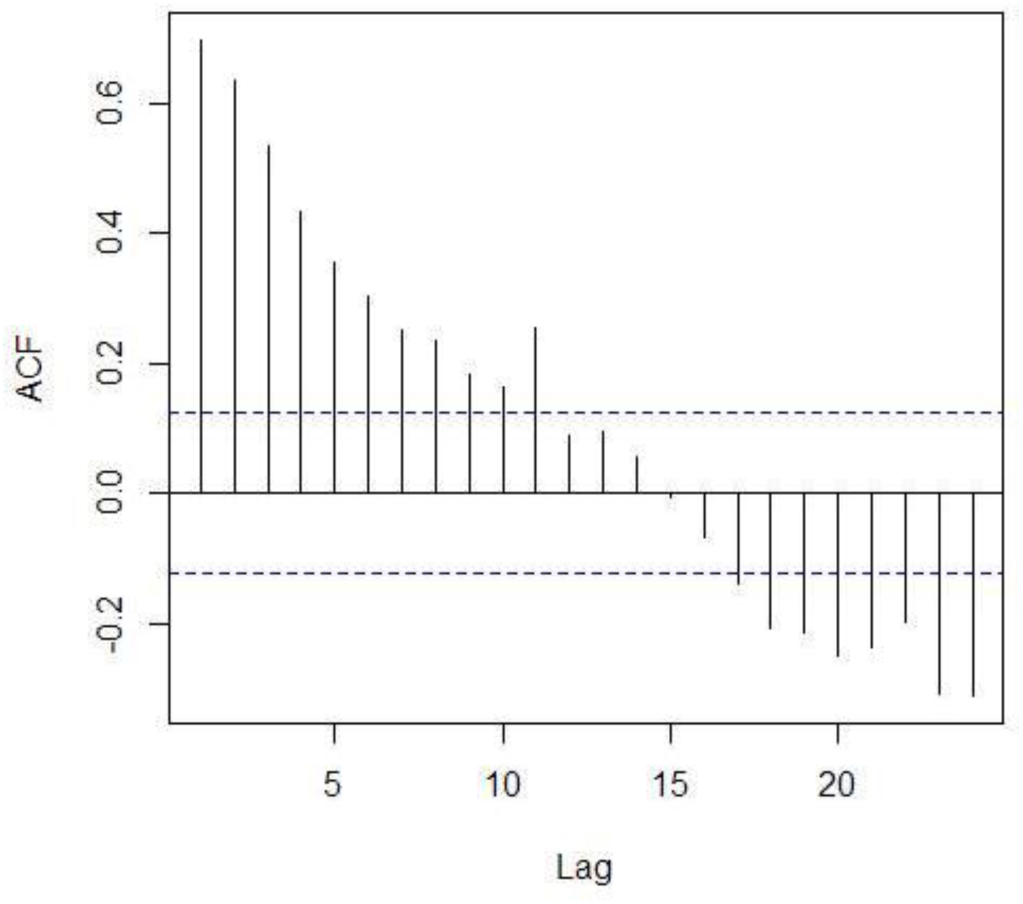
Autocorrelation Function (ACF) of Control A. (culture medium with NaCl 0.9%, without bacteria).

**Fig 17.**
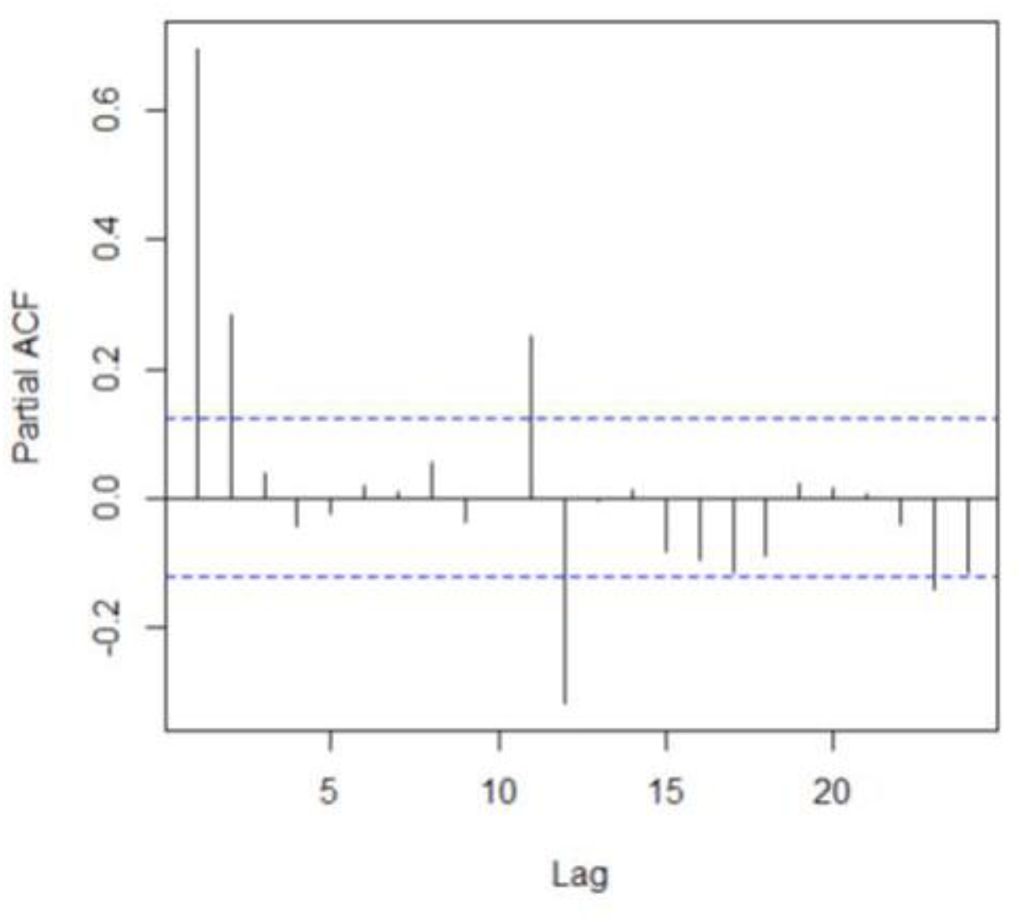
Partial Autocorrelation Function (PACF) of Control A. (culture medium with NaCl 0.9%, without bacteria).

A first-difference trend transformation was performed. The run sequence plot of the differenced data (Fig 18) shows that the mean of the differenced data is around zero, with the differenced result less autocorrelated than the original data. The ACF of the differenced data (Fig 19) and the PACF (Fig 20) both with a 95% confidence band show that only the autocorrelation at lag 1 and possibly at lag 2 are significant, suggesting that the differenced data are stationary.

**Fig 18.**
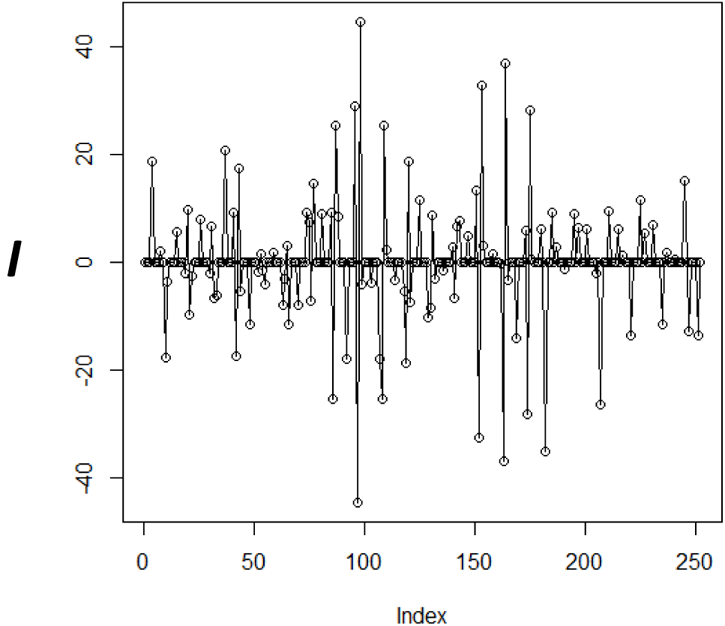
Run sequence plot of first-difference trend transformation of Control A. (culture medium with NaCl 0.9%, without bacteria).

**Fig 19.**
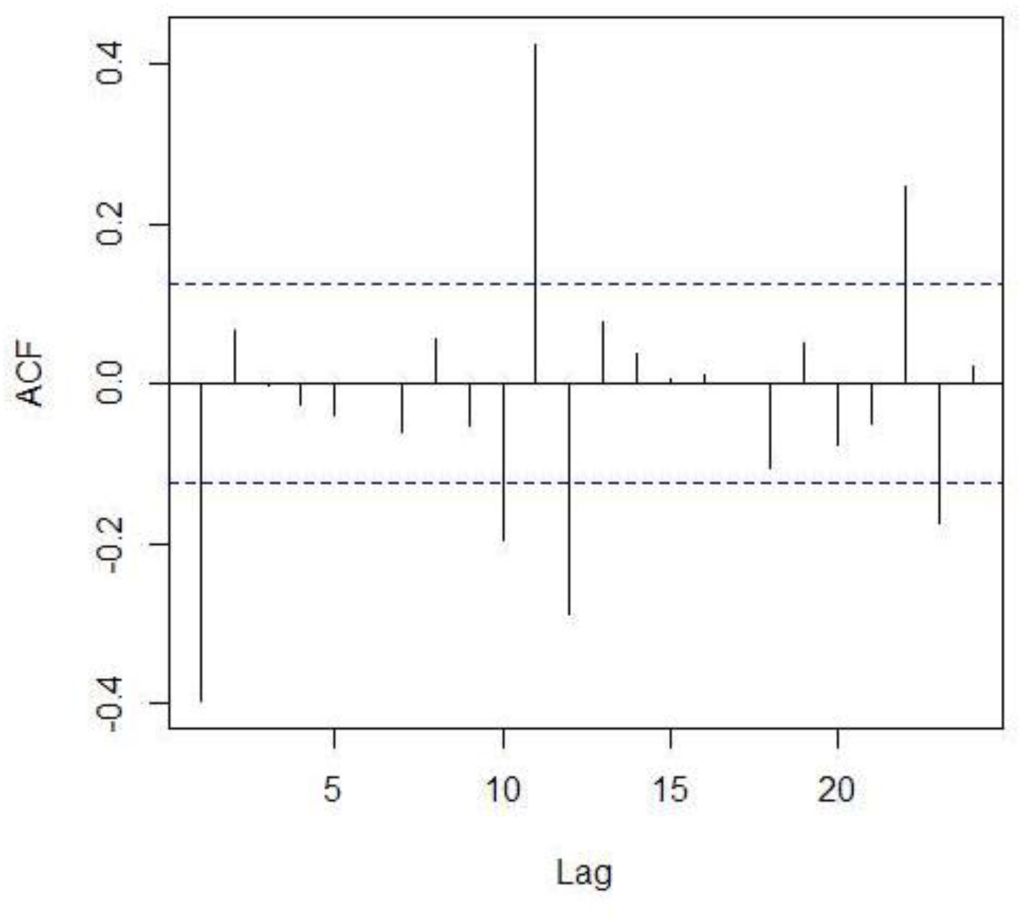
Autocorrelation Function (ACF) of first-difference trend transformation of Control A. (culture medium with NaCl 0.9%, without bacteria).

**Fig 20.**
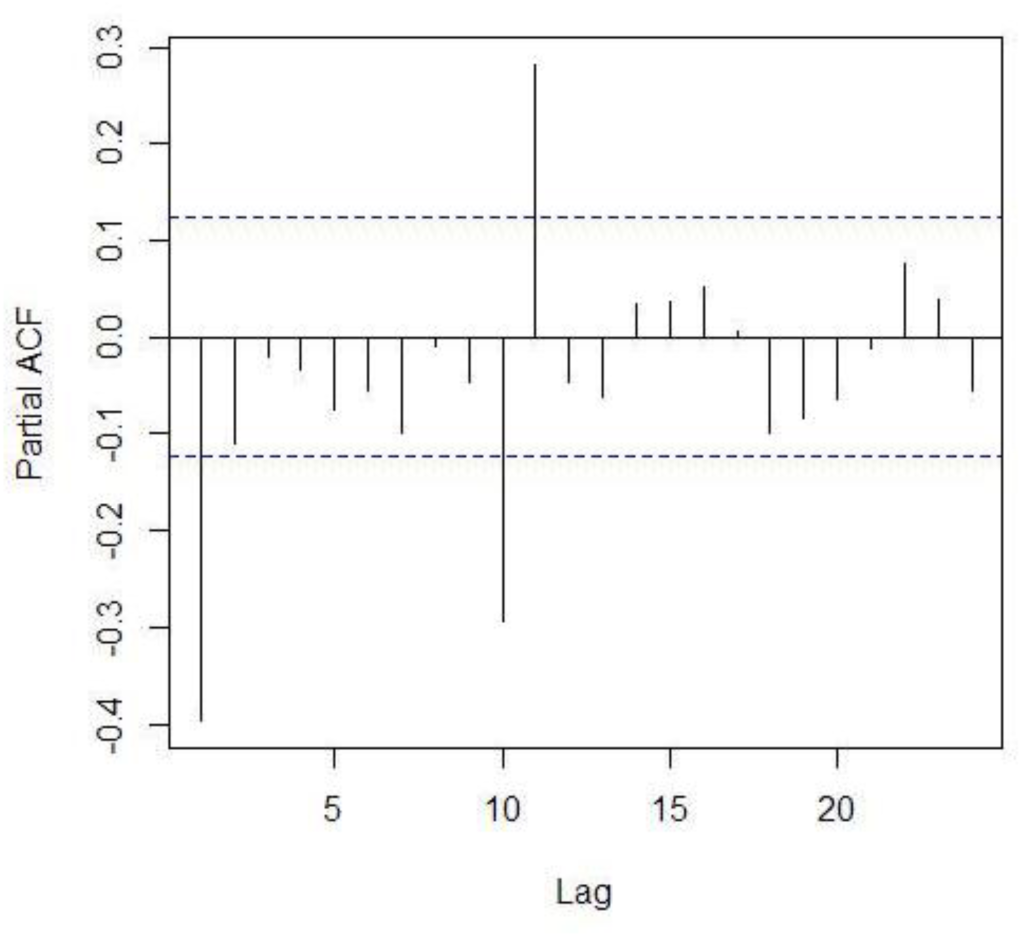
Partial Autocorrelation Function (PACF) of first-difference trend transformation of Control A. (culture medium with NaCl 0.9%, without bacteria).

Similarly, a first-difference seasonal transformation was performed, showing stationarity (Figs 21-23).

**Fig 21.**
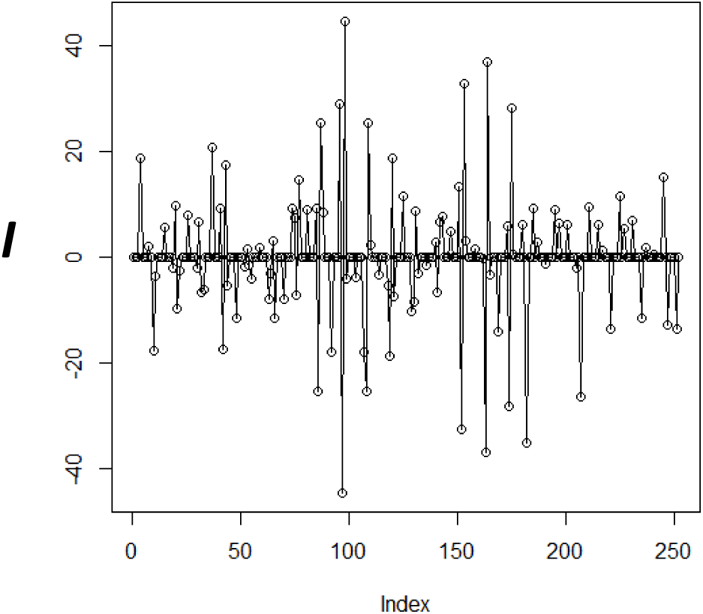
Run sequence plot of first-difference seasonal transformation of Control A. (culture medium with NaCl 0.9%, without bacteria).

**Fig 22.**
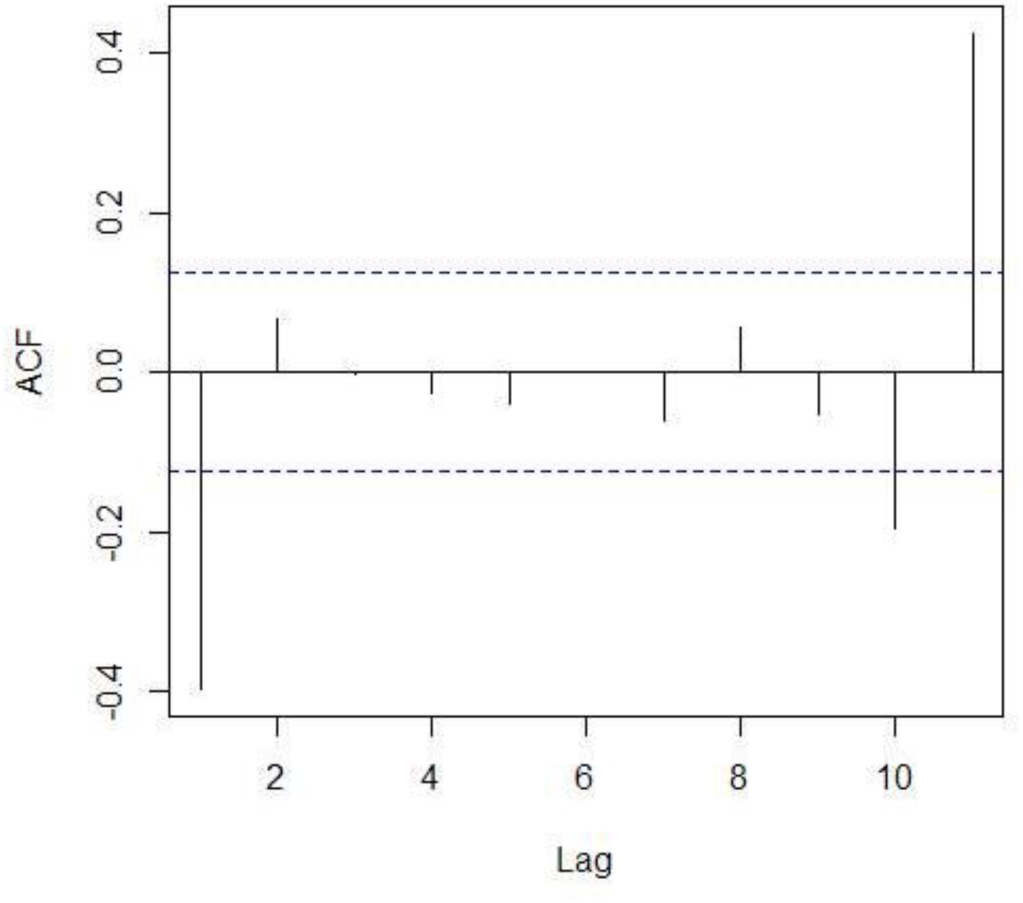
Autocorrelation Function (ACF) of first-difference seasonal transformation of Control A. (culture medium with NaCl 0.9%, without bacteria).

**Fig 23.**
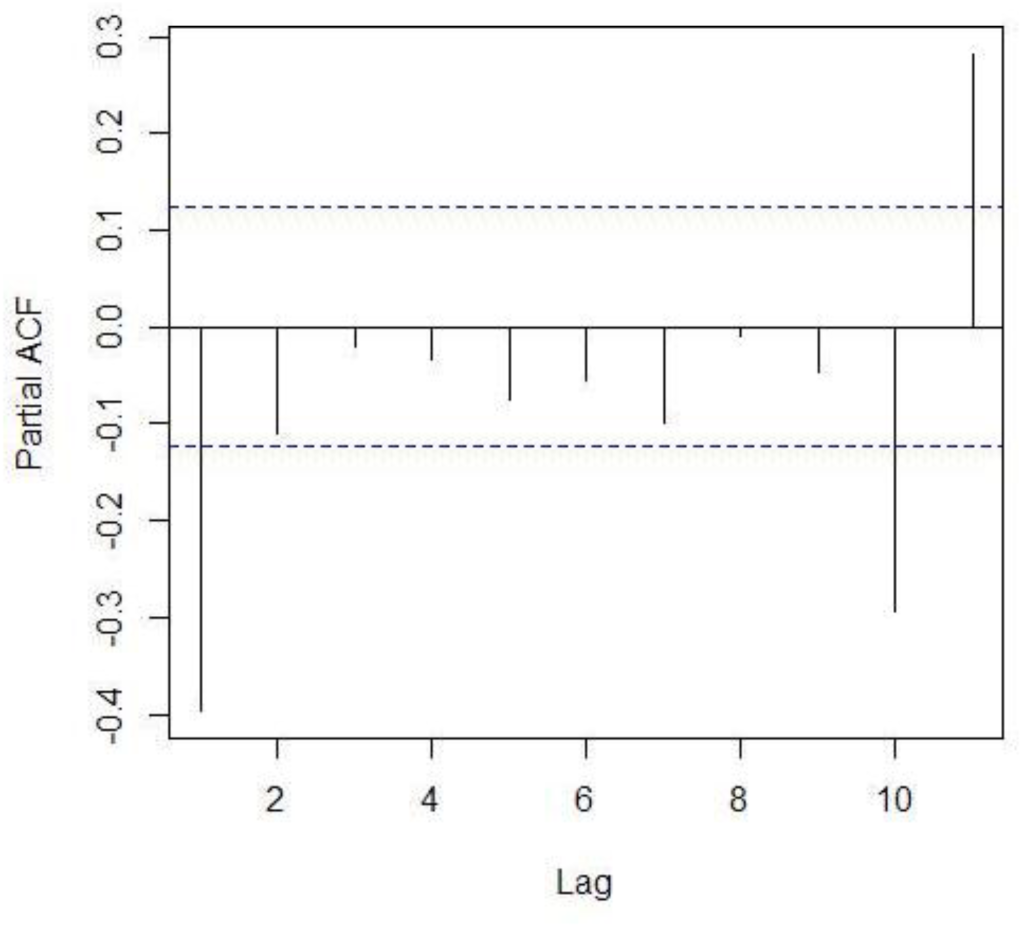
Partial Autocorrelation Function (PACF) of first-difference seasonal transformation of Control A. (culture medium with NaCl 0.9%, without bacteria).

Since the ACF plot of the original data shows that the autocorrelations are very strong and positive and decay very slowly, the time series was identified as an “AR(p) signature”. The PACF shows significance at lags 1 and 2 indicating that this is possibly an Auto-regressive, AR(2) signature. Since the ACF of the differenced data shows significance at lag 1 and possibly at lag 2, a Moving Average, MA(2) model is suggested.

Furthermore, these examinations indicate that the time series has trend and seasonality. Therefore, a term for trend (*d*) as well as the terms for seasonality (P,D,Q)_11_ were included. It should be noted that this rationale was used as a general orientation to select a model that could fit the largest number of videos obtained either with or without bacteria. Therefore, a (2,1,2)(2,1,0)_11_ model was thought to be close enough for this purpose and was chosen as the first selected model which has the further advantage of having been widely used.

Furthermore, a similar analysis of the time series for Videos 1 and 2 give the same orientation (see S3 Data). A similar process was carried out with Control C. The results are shown in Figs 24 to Fig 32. The run sequence plot (Fig 24) shows non-stationarity. The ACF (Fig 25) shows an “AR(p) signature” for which an AR(4) signature was suggested from the PACF (Fig 26), and an MA(1) was suggested from the ACF plot of the differenced data. Since stationarity is achieved with the differenced data, trend and seasonality were included in the model. The AR(4) can also be observed from the PACF of the differenced data. Taking into account that the most suitable ARIMA models are those with the lower number of parameters possible, a term (1,1,1) was assayed for variations in seasonality. The best fit was approached with a model (4,1,1)(1,1,1)_11_. Again, this rational was thought to be close enough for the purpose of having a starting criterion to guide towards finding the most suitable models.

**Fig 24.**
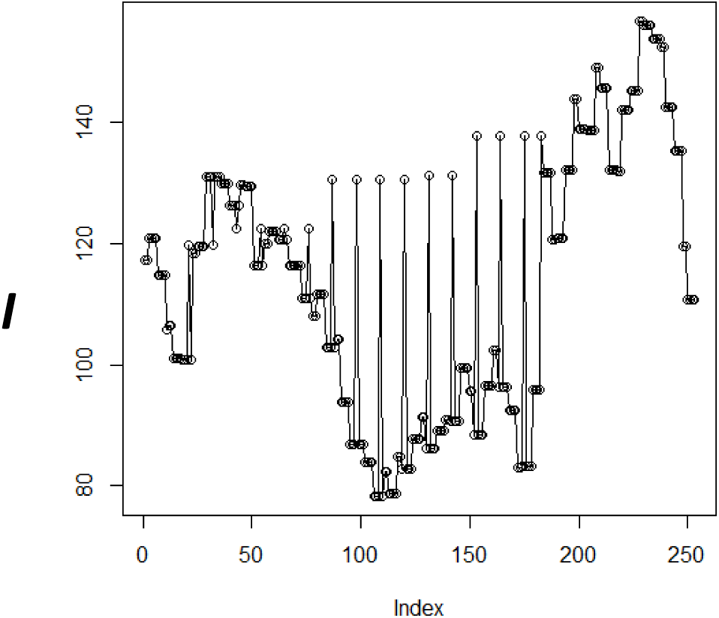
Run sequence plot of Control C. (culture medium without bacteria and without NaCl).

**Fig 25.**
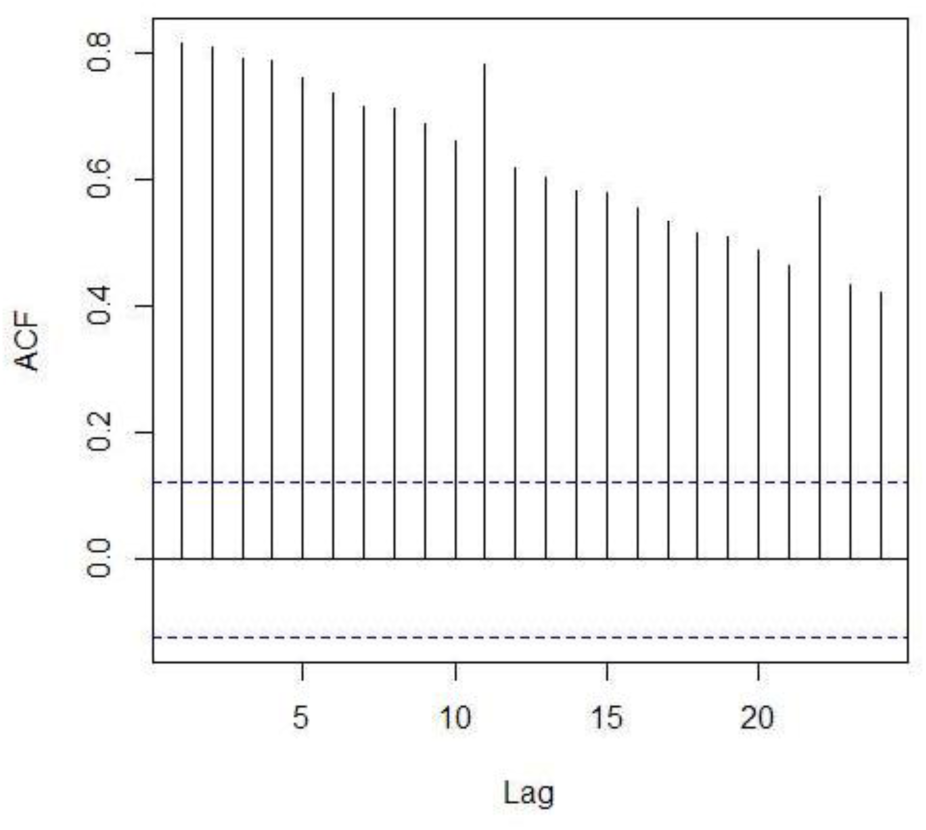
Autocorrelation Function (ACF) of Control C. (culture medium without bacteria and without NaCl).

**Fig 26.**
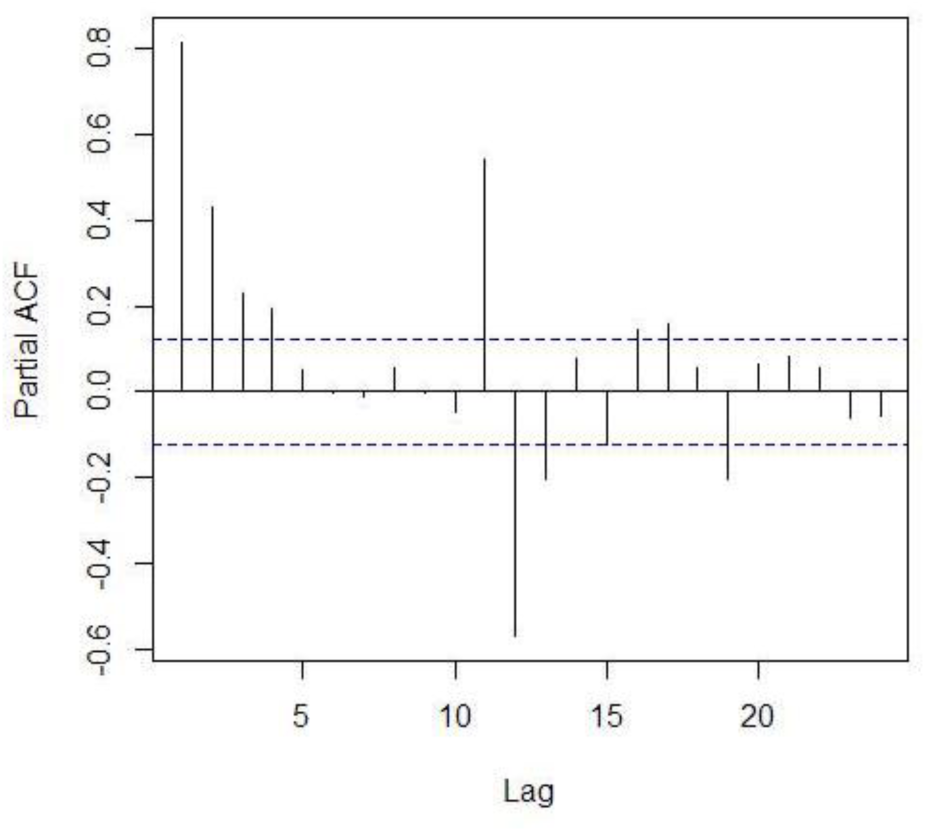
Partial Autocorrelation Function (PACF) of Control C. (culture medium without bacteria and without NaCl).

**Fig 27.**
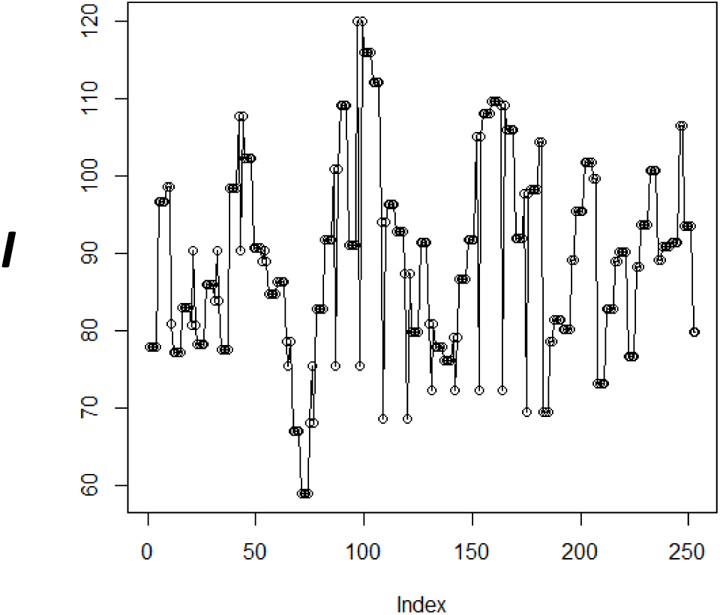
Run sequence plot of first-difference trend transformation of Control C. (culture medium without bacteria and without NaCl).

**Fig 28.**
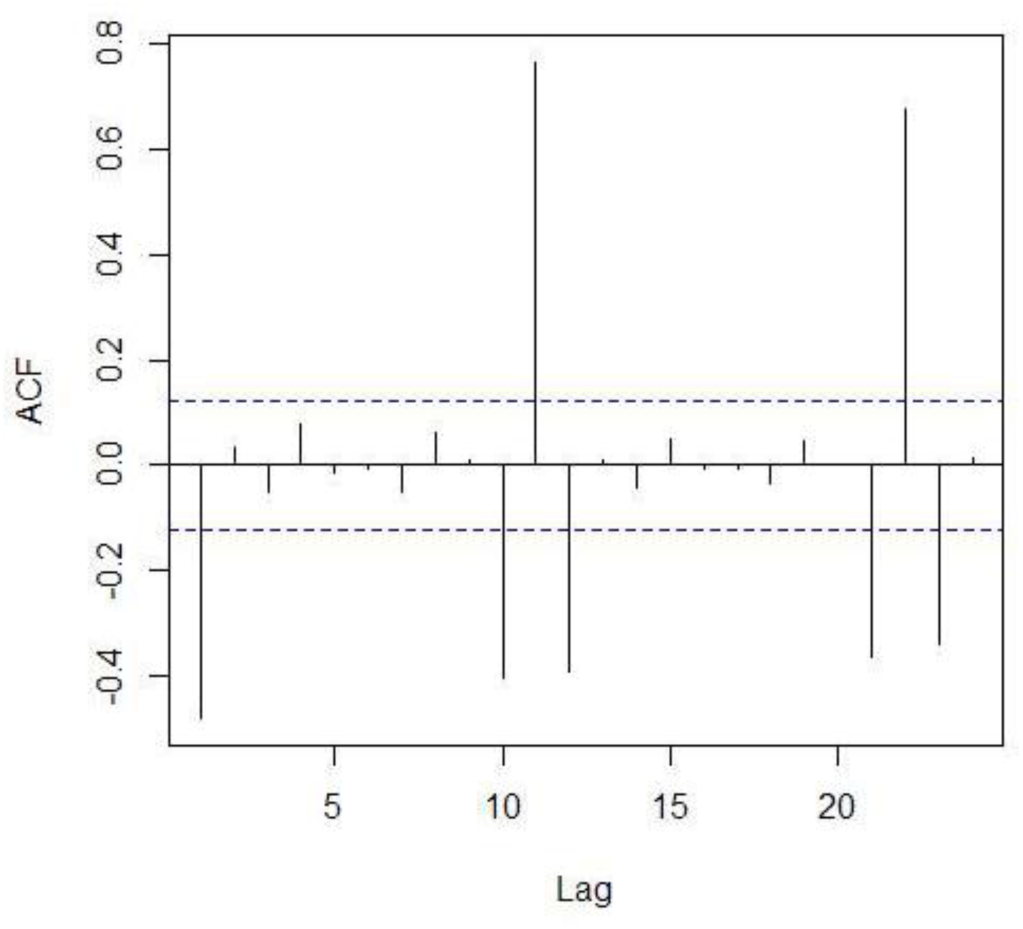
Autocorrelation Function (ACF) of first-difference trend transformation of Control C. (culture medium without bacteria and without NaCl).

**Fig 29.**
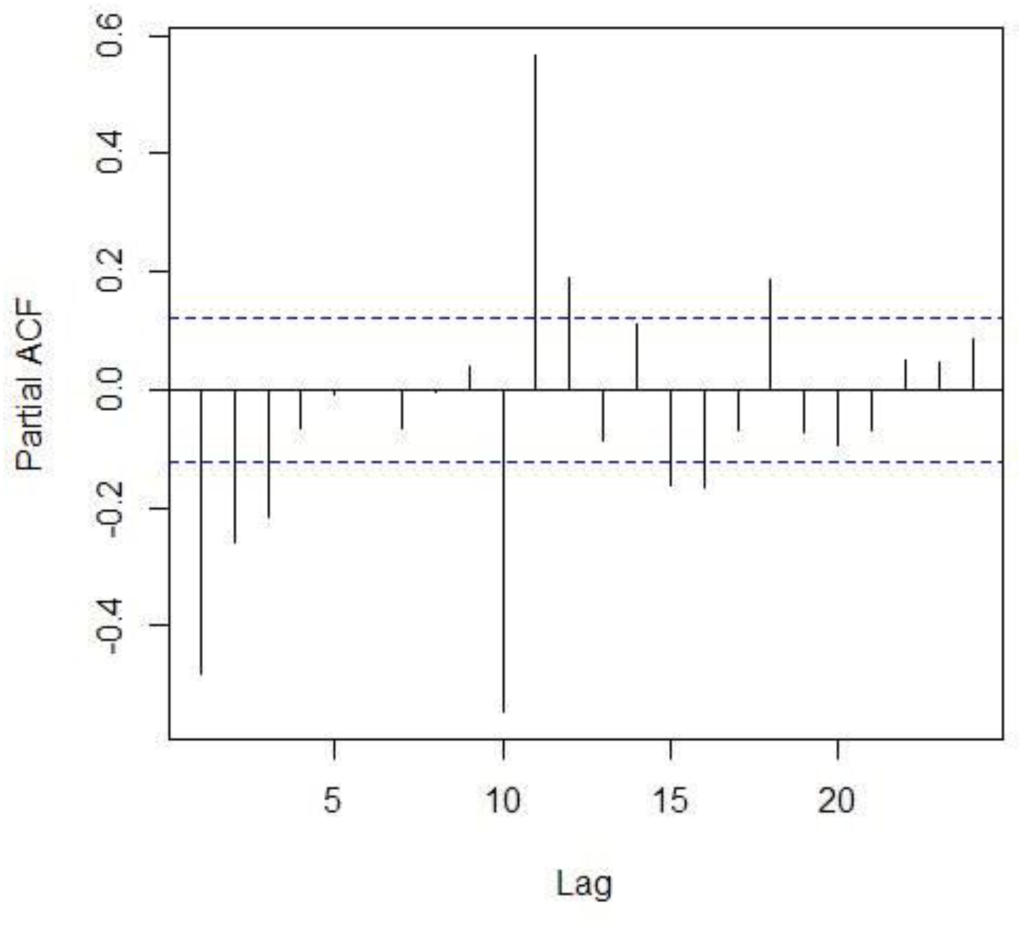
Partial Autocorrelation Function (PACF) of first-difference trend transformation of Control C. (culture medium without bacteria and without NaCl).

**Fig 30.**
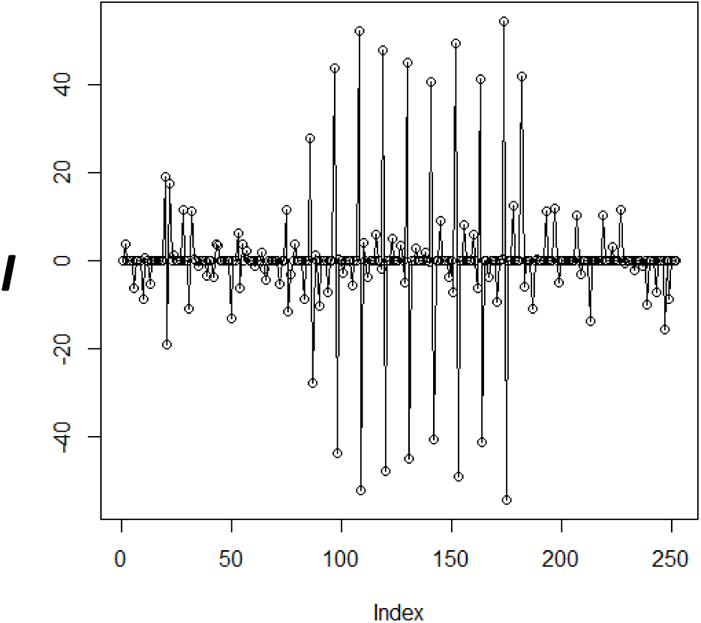
Run sequence plot of first-difference seasonal transformation of Control C. (culture medium without bacteria and without NaCl).

**Fig 31.**
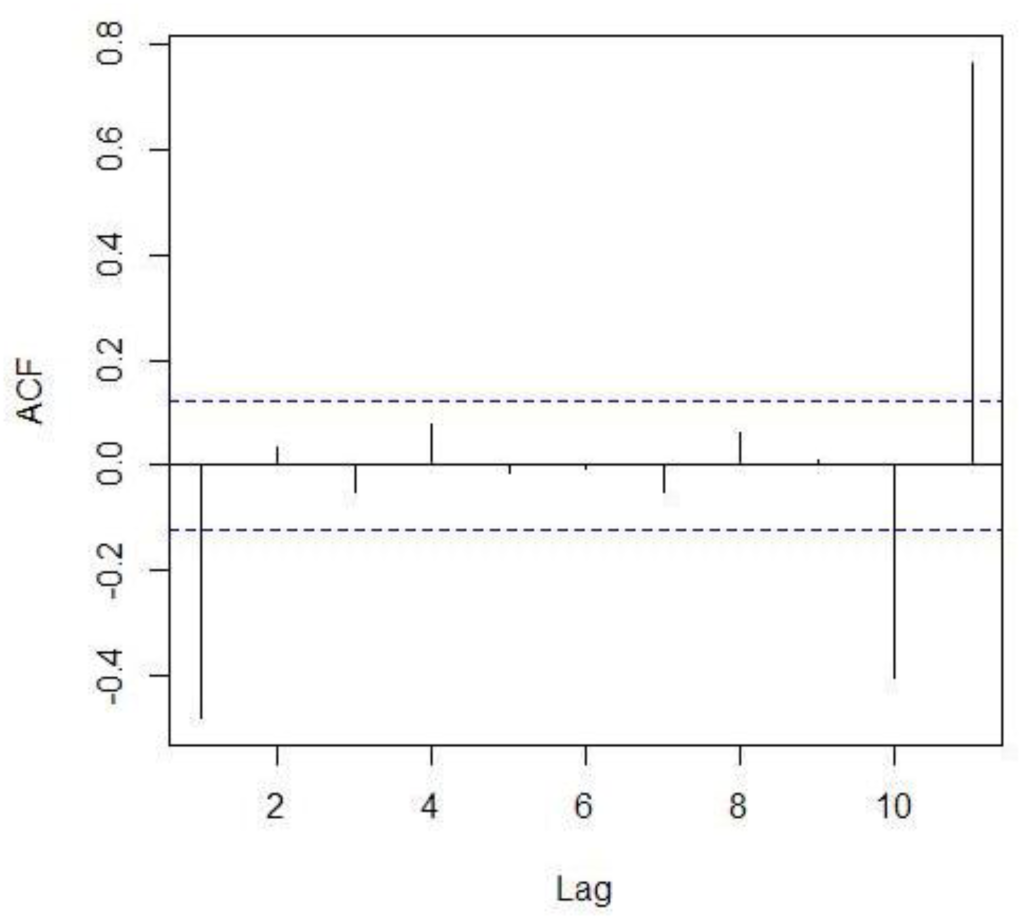
Autocorrelation Function (ACF) of first-difference seasonal transformation of Control C. (culture medium without bacteria and without NaCl).

**Fig 32.**
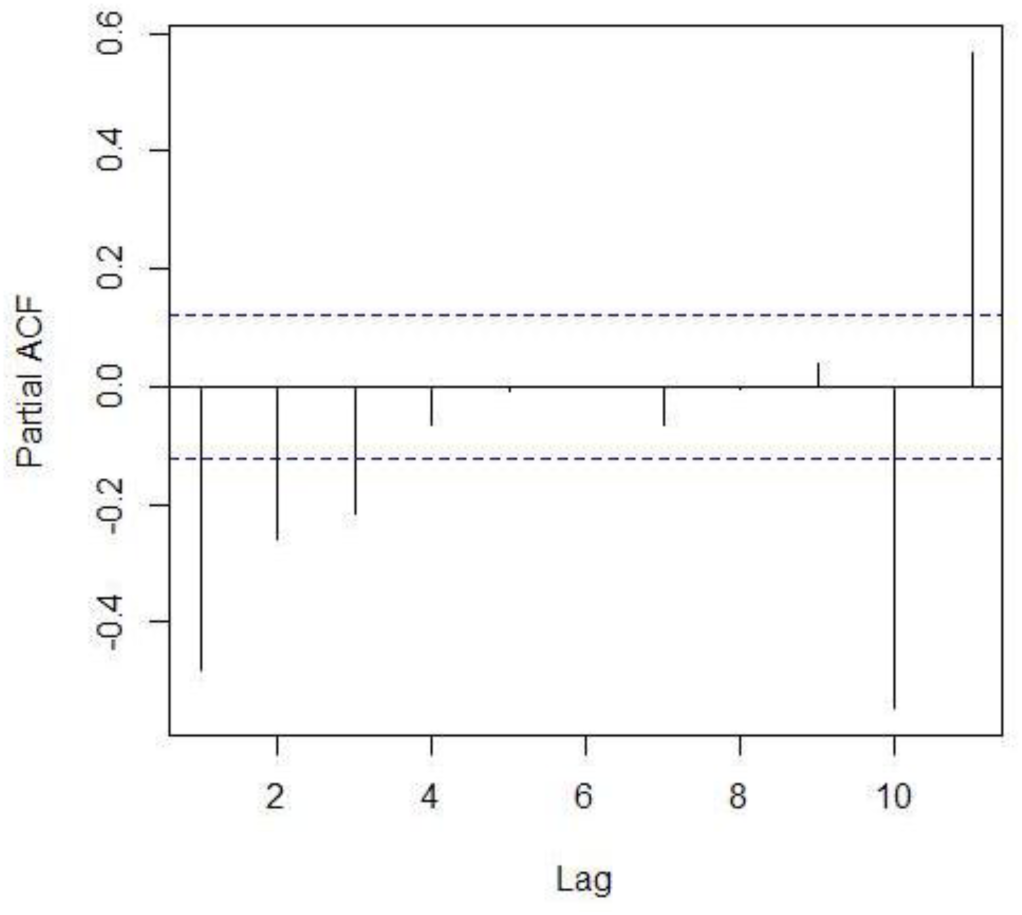
Partial Autocorrelation Function (PACF) of first-difference seasonal transformation of Control C. (culture medium without bacteria and without NaCl).

The different models were evaluated on the individual time series and the results are shown individually and also considered in terms of bacteria in the presence of Ciprofloxacin at low concentration (7.6E-04 to 2.4E-02, even numbered videos 2-12), intermediate concentration (4.88E-02 to 7.81E-01, even numbered videos 14 to 22) and high concentration (1.56E+00 to 1E+02, even numbered videos 24 to 36) of the antibiotic and the culture medium with all the concentrations of Ciprofloxacin (7.6E-04 to 1.0E+02, odd numbered videos 1-35).

Fig 33 shows the results of applying these two models ((2,1,2)(2,1,0)_11_ and (4,1,1)(1,1,1)_11_) to the 38 videos with and without bacteria and varying amounts of Ciprofloxacin. Black boxes symbolize that the model fits the time series, with coefficients that are <1 and with residuals that behave as white noise. Both models tend to fit better the time series for the odd numbered videos (that contain culture medium without bacteria with varying amounts of antibiotic), than the even numbered videos with bacteria and the low and intermediate amounts of antibiotic. Also, the time series from videos with bacteria and the highest amount of antibiotic, adjust to both models as would be expected if the antibiotic tends to affect the bacteria, behaving like the culture medium with antibiotic and without bacteria. Therefore, fitting models (2,1,2)(2,1,0)_11_ and (4,1,1)(1,1,1)_11_ on the time series, it is possible to distinguish among the conditions with and without bacteria and among antibiotic concentrations below and above the MIC.

**Fig 33.**
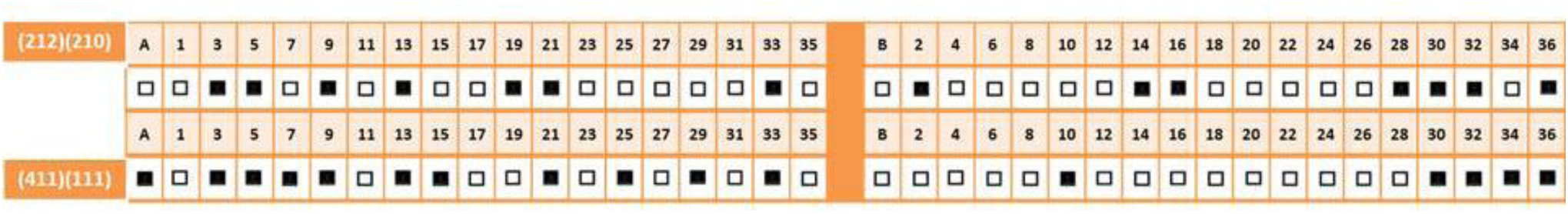
Adjustment of ARIMA models (2,1,2)(2,1,0)_11_ and (4,1,1)(1,1,1)_11_. Each one of these models was performed with each one of the time series that are indicated above the boxes. Black boxes indicates that the model adjusts to the time series with 253 values; white boxes indicates that the model does not adjust to the time series.

In order to be able to further distinguish the groups of assays, several models were tested separately to find the best fit for the culture medium without bacteria, the affected bacteria and the unaffected bacteria, increasing and decreasing the parameters of the ARIMA models.

### Increasing the parameters of the ARIMA models

Since the purpose of this work is to distinguish the presence and absence of bacteria and the effect of the antibiotic, variations of the model were performed so as to obtain the best fit with the different conditions. Therefore, in an attempt to detect the bacteria that have been least affected by the antibiotic and are probably the most active, alternative models were assayed. Since these time series have an “AR(p) signature”, different models with increasing values of (p) were tested, starting from AR(2), MA(2). Fig 34 shows the best fitted model for this purpose, (7,1,2)(1,0,1)_11_, adjusting to the time series of bacteria in the presence of the least amount of antibiotic. It should be noted that in the case of the even numbered videos (with bacteria) a tendency that fades off as the antibiotic concentration rises from low to intermediate is seen, and it is not an “all or nothing” result, again resembling a transient condition (see Figs 12 and 13).

**Fig 34.**
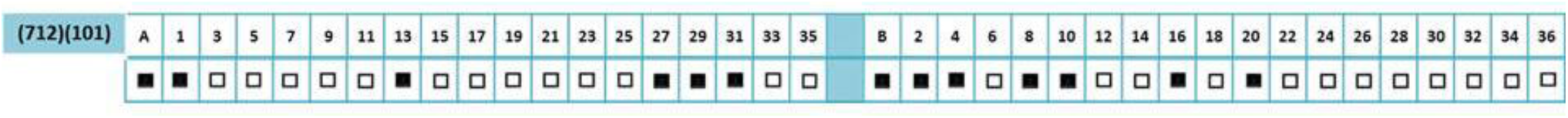
Adjustment of ARIMA model (7,1,2)(1,0,1)_11_. The model was performed with each one of the time series that are indicated above the boxes. Black boxes indicates that the model adjusts to the time series with 253 values; white boxes indicates that the model does not adjust to the time series.

### Decreasing the parameters of the ARIMA model

Since it is recommended that the most suitable ARIMA models are those with the lower number of parameters, a decrease in the numbers of the model was assayed, with variations in seasonality. Thus, the model **(1,1,1)** was assayed, combined with the seasonal parameters (P,D,Q)_11_ that were permutations of 1 and 0 (0,0,0); (0,0,1); (0,1,0); **(0,1,1)**; **(1,0,0)**; **(1,0,1)**; (1,1,0); **(1,1,1)**. Fig 35 shows only the results for 4 selected models which are highlighted in this text. All of them are able to detect the bacteria affected by the higher antibiotic concentrations. The other combinations (not highlighted) do not adjust completely to the time series.

**Fig 35.**
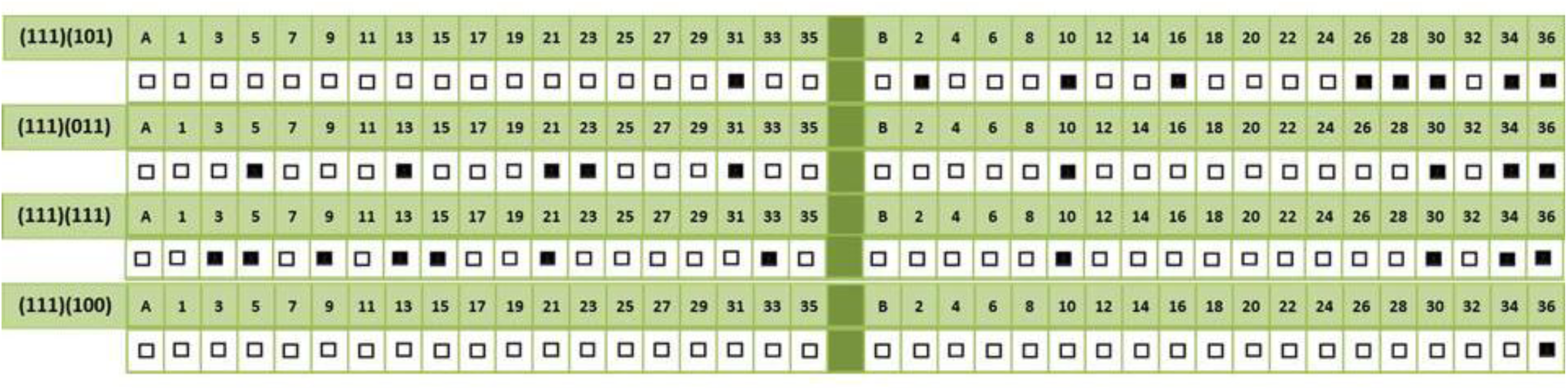
Adjustment of ARIMA models (1,1,1)(1,0,1)_11_; (1,1,1)(0,1,1)_11_; (1,1,1)(1,1,1)_11_and (1,1,1)(1,0,0)_11_. Each one of these models was performed with each one of the time series that are indicated above the boxes. Black boxes indicates that the model adjusts to the time series with 253 values; white boxes indicates that the model does not adjust to the time series.

Model (1,1,1)(1,0,0) does not fit any of the assays of culture medium without bacteria and model (1,1,1)(1,0,1) fits only one time series in this group. These models, especially the latter, are able to detect the bacteria affected by the higher antibiotic concentrations. Furthermore, model (1,1,1)(1,0,1) has a pattern that is similar to the result obtained with Shapiro-Wilk Normality test (see Fig 11). In general, the first 3 models tend to have an adjustment with the time series of assays with bacteria and the highest antibiotic concentration. In other words, these models have the tendency to distinguish the bacteria that are affected by the concentrations of Ciprofloxacin that are above the MIC that also show a normal distribution of the data.

### Antimicrobial susceptibility testing

In order to test the susceptibility of the *E*. *coli* K-12 strain HfrH to the Ciprofloxacin i.v. infusion used in this work, classic broth and agar dilution methods were performed to determine the lowest concentration of the assayed antimicrobial agent (MIC) that, under defined test conditions, inhibits the growth of the bacterium. The broth macro-dilution method showed a MIC value of 0.488μg/mL (S4 Data) which corresponds with a concentration that approaches the assays of Videos 19 and 20 (0.39 μg/mL). Furthermore, Fig 36 shows the results of the mean and SD of the growth of the bacteria as determined by its Absorbance at 625nm arranged in three groups of low, intermediate and high antibiotic concentration that are similar to those of Fig 10. The low concentration group goes from 1.53E-02 to 2.44E-01, the intermediate concentration group goes from 4.88E-01 to 7.81E00 (including the MIC) and the high concentration group goes from 1.56E+01 to 2.50E+02. The results obtained with agar dilution indicate (see S4 Data) that 5 μg of Ciprofloxacin have an inhibition diameter that is ≥21mm as required in the CLSI standards [1] for a susceptible strain. Furthermore, in this agar dilution test [28], a concentration of 0.488 μg/mL, which again is close to the assays of Videos 19 and 20 (0.39 μg/mL) of the Biospeckle technique, is the minimal concentration that promotes a precise inhibition halo which can be detected and measured under the magnifying microscope. All this indicates that Ciprofloxacin has the expected anti-microbial activity on the selected strain of *E*. *coli* K-12 and that the antimicrobial susceptibility tests show results that may be comparable to those obtained with Biospeckle under different analysis methods.

**Fig 36.**
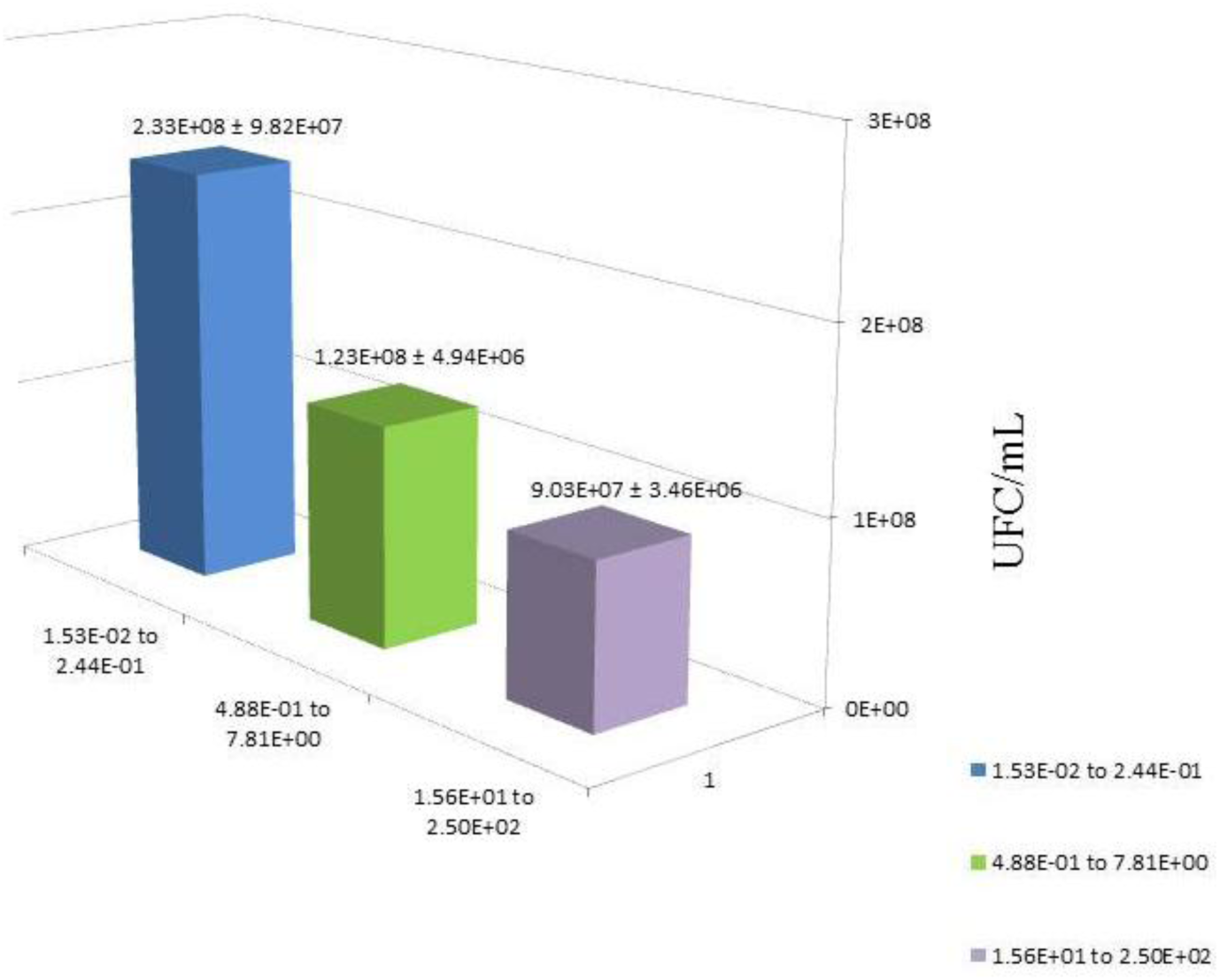
Absorbance at λ_625nm_ of the macro-broth dilution test. The macro-broth dilution test was carried out as described in Materials and Methods with bacteria and varying concentrations of Ciprofloxacin, obtaining in each case, a value of Absorbance which represents a measure of the growth of the microorganism. The y axis expresses the CFU/mL. The results of the assays with bacteria in MHB and varying antibiotic concentrations are organized in groups that are similar as the groups of Fig 10 because, due to the experimental design, variations in concentrations are obtained, however, the same groups below and above the MIC are presented, as follows, Group (1) assays with Ciprofloxacin concentration from 3.81E-03 to 3.05E-02; Group (2) assays with Ciprofloxacin concentration from 6.10E-02 to 9.77E-01; and Group (3) assays with Ciprofloxacin concentrations from 1.95E+00 to 1.25E+02.

## DISCUSSION

An assay designed to develop dynamic laser speckle or Biospeckle aims at providing a digital signal which is caught in a video. Biologic materials [29] as well as the illumination in the setup tend to cause inhomogeneities that affect the digital processing of the images. Furthermore, subsurface inhomogeneities have been reported and revealed with laser speckle [30]. These inhomogeneities could interfere if the whole photogram is used for calculations, obscuring the real effect that is to be revealed. Therefore, the selection of a ROI is an important step prior to the analysis of the results. Braga *et al* [29] have addressed the problem of inhomogeneities by selecting pixels at random, whereas Meschino *et al* have performed a Self-Organizing Map based on Artificial Neural Networks. According to those authors, care must be taken so as to select a representative area for calculation. Therefore, in the present work a quantitative approach was developed. The rational indicated that areas of the photogram with high illumination should be avoided because small changes could be obscured within the high intensity (*I*) peaks. Areas of high variation should also be avoided because the variable background could make the activity of the bacteria undetectable, due to their size, their motility and the high cell density usually required. In this context and with the idea of performing statistical analysis, a low resolution of the camera was selected. The quantitative analysis of the SD of 11 pixels in 11 photograms in terms of a video with bacteria and a video without bacteria, both with the lowest Ciprofloxacin concentration, revealed an appropriate approach to find the ROI which is located in the area of low intensity (*I*).

After having identified the ROI, three approaches were taken which focus on the evolution of intensity (*I*) as a function of the time domain. For this purpose the behavior for a length of time of either an area of 900 pixels or of the time series of individual pixels, was examined. In the first approach, after quantitatively identifying the ROI, the segment was punched out of each frame and all the extracted segments were used to edit a new video with 80 segmented frames which corresponds to a time period of over 2 seconds. The number of frames was chosen from previous studies [16] in which it was determined that 80 frames was an appropriate length for ImageDP processing. With the combination of ImageJ for video construction and ImageDP for video processing (Fig 2), it was possible to obtain a linear tendency which shows that the mean intensity (*A*) of the Biospeckle pattern of the bacteria decreases as the antibiotic concentration increases. However, since there is a high dispersion of the points, the analysis was improved organizing the results within groups above and below the MIC. For the other two approaches, a pixel with a high differential activity was chosen within the selected ROI, for the extraction of a time series with a length of over 8 seconds and then analyzing each one either through diagnostic statistical tests or by adjusting an ARIMA model for each time series.

The diagnostic statistical tests were chosen in reference to linear regressions. In other words, a linear regression model and three of its principal assumptions (normality, homoscedasticity and independence) were tested on the time series as a repeated-measures analysis. Only the time series of the least affected bacteria (low Ciprofloxacin concentration) behaved in an expected manner, being non independent and mostly non-linear, non-normal and heteroscedastic. On the other hand, the time series of the most affected bacteria (higher Ciprofloxacin concentration) are non-independent and tend to be linear, normal and heteroscedastic; and the time series of the culture medium without bacteria (all Ciprofloxacin concentrations) are non-independent and non-normal and tend to be linear and homoscedastic. Therefore, adjustment to a linear regression identifies both, the culture medium and the most affected bacteria, normality identifies the most affected bacteria, and heteroscedasticity-homoscedasticity distinguishes the presence-absence of bacteria, respectively.

Through the use of ARIMA models, we explored the relationship between the activities of the Biospeckle pattern of *E*. *coli* K-12 as it becomes increasingly affected by Ciprofloxacin. Thus, the activity of the Biospeckle pattern was modeled as a univariate ARIMA with a seasonal component which indicates that the values not only depend on their own past values and random errors, but also they depend on the values of a previous cycle. Based on the ACF and PACF we fitted several ARIMA models with varying AR and MA orders. In the fitting process, the AR and MA coefficients were estimated using the log-likelihood method. The residuals were further inspected for autocorrelation through ACF and PACF. Models with autocorrelated residuals or with coefficients >1, were discarded. ARIMA models were adjusted showing that a (1,1,1)(1,0,1)_11_ model identifies the most affected bacteria (higher antibiotic concentration); a (4,1,1)(1,1,1)_11_ model is able to identify the time series of the most affected bacteria (higher antibiotic concentration) and of the culture medium (with varying antibiotic concentrations), indicating that the bacteria, as they become affected by increasing concentrations of the antibiotic, tend to behave in the same manner as the culture medium. The time series of the least affected bacteria (lowest antibiotic concentration) are identified by a (7,1,2)(1,0,1)_11_ model. The non-linear, non-normal and heteroscedastic behavior of this group is probably responsible for its adjustment to a model with a relatively high parameter. Thus, with the three methods it is possible to distinguish the presence and absence of bacteria and also bacteria with antibiotic concentrations below and above the MIC.

Bacteria move with a run-and-tumble navigation. During runs the organism moves approximately straight and during tumbles, or reorientations at zero speed, the organism explores other directions [31]. This strategy is affected by the presence of positive and negative feedbacks. In the presence of a gradient it has been described that there is a non-normal dynamical structure of the positive feedback and a quadratic non-linearity in the speed along the gradient [13]. This is in agreement with the results of this work in the case of the time series of bacteria at low concentrations of Ciprofloxacin. Moreover, in *Bacillus subtilis*, a motile Gram positive bacterium, it has been shown that under sub-lethal antibiotic concentrations, the statistics of collectively swarming cells, experiments a transition from normal to anomalous [32]. The authors conclude that the transition is due to changes in the properties of the bacterial motion and the formation of a motility-defective sub-population that self-segregates into regions. This phenomenon suggests a new strategy bacteria employ to fight antibiotic stress. In the present work this phenomenon could represent the transient condition of sub-lethal concentrations of antibiotic ranging from 4.88E-02μg/mL to 7.81E-01 μg/mL. This group is flanked on one side by the very low sub-lethal antibiotic concentrations (7.63E-04μg/mL to 2.44E-02μg/mL) and on the other side by the high lethal antibiotic concentrations (1.56 μg/mL to 1.00E+02μg/mL). The effect of Ciprofloxacin at these concentration ranges was confirmed in the antimicrobial susceptibility tests performed on the bacterial strain by macro-broth and agar dilution assays by growth inhibition and by the size of the diameter of the inhibition halo respectively, as established by CLSI [1]. Moreover, the results of these microbiological tests show a profile that is comparable to the result obtained with the images after 15min of antibiotic action and with ImageJ-ImageDP processing. In this work, a combination of a low intensity (I) of the selected region (ROI) and a low resolution of the camera were adequate to manage the data with several methods. This approach was appropriate for this case in which the microorganism is a bacterium which has a small size, high motility and is used at a high cell density. Moreover, all the performed analysis lead to comparable results possibly due to the quantitative method used for the selection of the ROI, on which the edition of the new video and the selection of a time series were performed. This allowed both, the edition and quantification with a “flip-book animation” extracted at the ROI and the selection of a pixel for the time series construction for diagnostic statistics. The four evaluation methods used: diagnostic statistical tests, fitting of ARIMA models, ImageJ-ImageDP processing and antimicrobial susceptibility tests, show similar results, being able to distinguish among the groups of assays with bacteria and Ciprofloxacin below and above the MIC. This result opens a possible way to perform a fast method to evaluate antibiotic susceptibility in control as well as in clinical samples.

## CONCLUSIONS

A Biospeckle laser assay was designed for the evaluation of Ciprofloxacin susceptibility of a strain of *E*. *coli* K-12. Digital image processing was performed by three methods that were comparable to classical antimicrobial susceptibility tests in statistical terms. Diagnostic statistical tests, fitting of ARIMA models and ImageJ-ImageDP processing showed results that were comparable to antimicrobial susceptibility tests. This Biospeckle laser method is a possible starting point for testing body fluids and other samples and it opens a possible way to perform a fast method to evaluate antibiotic susceptibility.

## Supporting Information

**S1 DATA.** Data of the edited videos using ImageJ–ImageDP

**S2 DATA.** Diagnostic statistical tests (linear regression model, normality, homoscedasticity and independence) of the time series for each video

**S3 DATA.** Run sequence plot, ACF and PACF for the time series of Video 1 and Video 2.

**S4 DATA.** Data of the antimicrobial susceptibility tests

**S1 TEXT.** ImageDP Manual for online processing

## Acknowledgements

We thank Centro Multidisciplinario de Ciencias, Instituto Venezolano de Investigaciones Científicas(IVIC-Mérida) and Sección Biotecnología, Instituto de Investigaciones, Facultad de Farmacia y Bioanálisis, Universidad de Los Andes, Mérida, Venezuela. The advice of Francisco A. Andrades-Grassi is gratefully acknowledged.

## Supporting Information

### S1 DATA

Data of the edited videos using ImageJ–ImageDP

#### Biospeckle laser digital image processing for quantitative and statistical evaluation of the activity of Ciprofloxacin on *Escherichia coli* K-12

Hilda Cristina Grassi, Ana Velásquez, Olga Mercedes Belandria, María Lorena Lobo-Sulbarán, Jesús E. Andrades-Grassi, Humberto Cabrera, Efrén D. J. Andrades

The following table shows the values of *A* which are presented in Fig 9 as the linear regression for the values of *A* as a function of the logarithm of the concentration of Ciprofloxacin.

| Ciprofloxacin concentration<br>$\mu\text{g/mL}$ | Video number<br>(without bacteria) | $A$<br>(a.u) | Video number<br>(with bacteria) | $A$<br>(a.u.) |
| --- | --- | --- | --- | --- |
| 7.600E-04 | 1 | 0.163533 | 2 | 0.334237 |
| 1.520E-03 | 3 | 0.137181 | 4 | 0.36378 |
| 3.050E-03 | 5 | 0.16193084 | 6 | 0.25338 |
| 6.100E-03 | 7 | 0.149833 | 8 | 0.313928 |
| 1.200E-02 | 9 | 0.14169385 | 10 | 0.289908 |
| 2.400E-02 | 11 | 0.19969003 | 12 | 0.311146 |
| 4.800E-02 | 13 | 0.204874 | 14 | 0.302165 |
| 9.700E-02 | 15 | 0.17109235 | 16 | 0.208719 |
| 1.900E-01 | 17 | 0.11410259 | 18 | 0.255644 |
| 3.900E-01 | 19 | 0.17901294 | 20 | 0.232792 |
| 7.800E-01 | 21 | 0.15331705 | 22 | 0.232792 |
| 1.560E+00 | 23 | 0.12586017 | 24 | 0.244527 |
| 3.120E+00 | 25 | 0.2215253 | 26 | 0.255748 |
| 6.250E+00 | 27 | 0.19220906 | 28 | 0.206645 |
| 1.250E+01 | 29 | 0.16066046 | 30 | 0.19108 |

The following table shows the values of *A* in groups of low, intermediate and high concentration of Ciprofloxacin, with and without bacteria. It should be noted that each group has 5 videos.

| GROUP | VIDEOS | CONCENTRATION<br>RANGE<br>µg/mL | MEAN<br>of the A values | STANDARD<br>DEVIATION<br>of the A values |
| --- | --- | --- | --- | --- |
| Bacteria with low concentrations of Ciprofloxacin | Even numbered videos 2-10 | 7.600E-04 to 1.200E-02 | 0.311 | 0.042 |
| Without Bacteria with low concentrations of Ciprofloxacin | Odd numbered videos 1-9 | 7.600E-04 to 1.200E-02 | 0.151 | 0.012 |
| Bacteria with intermediate concentrations of Ciprofloxacin (including the MIC) | Even numbered videos 12-20 | 2.400E-02 to 3.900E-01 | 0.262 | 0.044 |
| Without Bacteria with intermediate concentrations of Ciprofloxacin (including the MIC) | Odd numbered videos 11-19 | 2.400E-02 to 3.900E-01 | 0.174 | 0.036 |
| Bacteria with high concentrations of Ciprofloxacin | Even numbered videos 22 to 30 | 7.800E-01 to 1.250E+01 | 0.226 | 0.027 |
| Without Bacteria with intermediate concentrations of Ciprofloxacin | Odd numbered videos 21 to 29 | 7.800E-01 to 1.250E+01 | 0.171 | 0.037 |

### S1 TEXT

ImageDP Manual for online processing

#### Biospeckle laser digital image processing for quantitative and statistical evaluation of the activity of Ciprofloxacin on *Escherichia coli* K-12

Hilda Cristina Grassi, Ana Velásquez, Olga Mercedes Belandria, María Lorena Lobo-Sulbarán, Jesús E. Andrades-Grassi, Humberto Cabrera, Efrén D. J. Andrades

### ImageDP Manual

1. Preparation of the video for processing.
  a. The video must be named with numbers (no letters are allowed).
  b. The video must be a.wmv file.
2. ImageDP processing.
  a. Open http://bigjocker.com/biospeckle/
  b. The following will open:

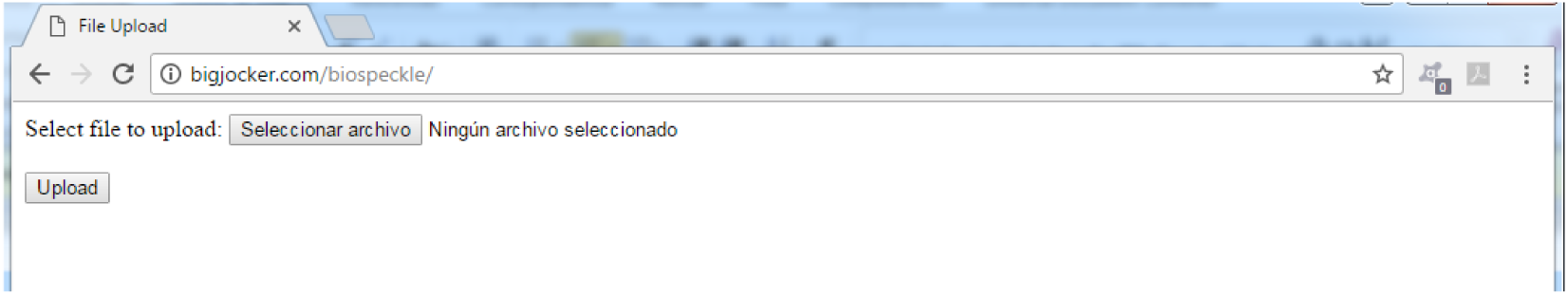
  c. “Seleccionar archivo” will guide the user through the selection of the file. The name of the file will appear substituting “Ningún archivo seleccionado”.

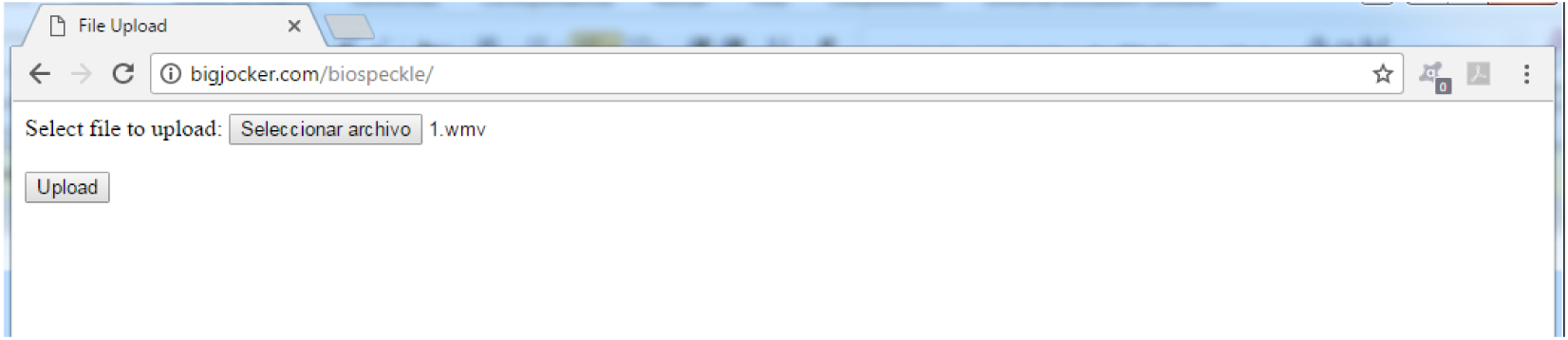
  d. “Upload” will accomplish the processing of the file.
  e. When the processing is done, the following will open:

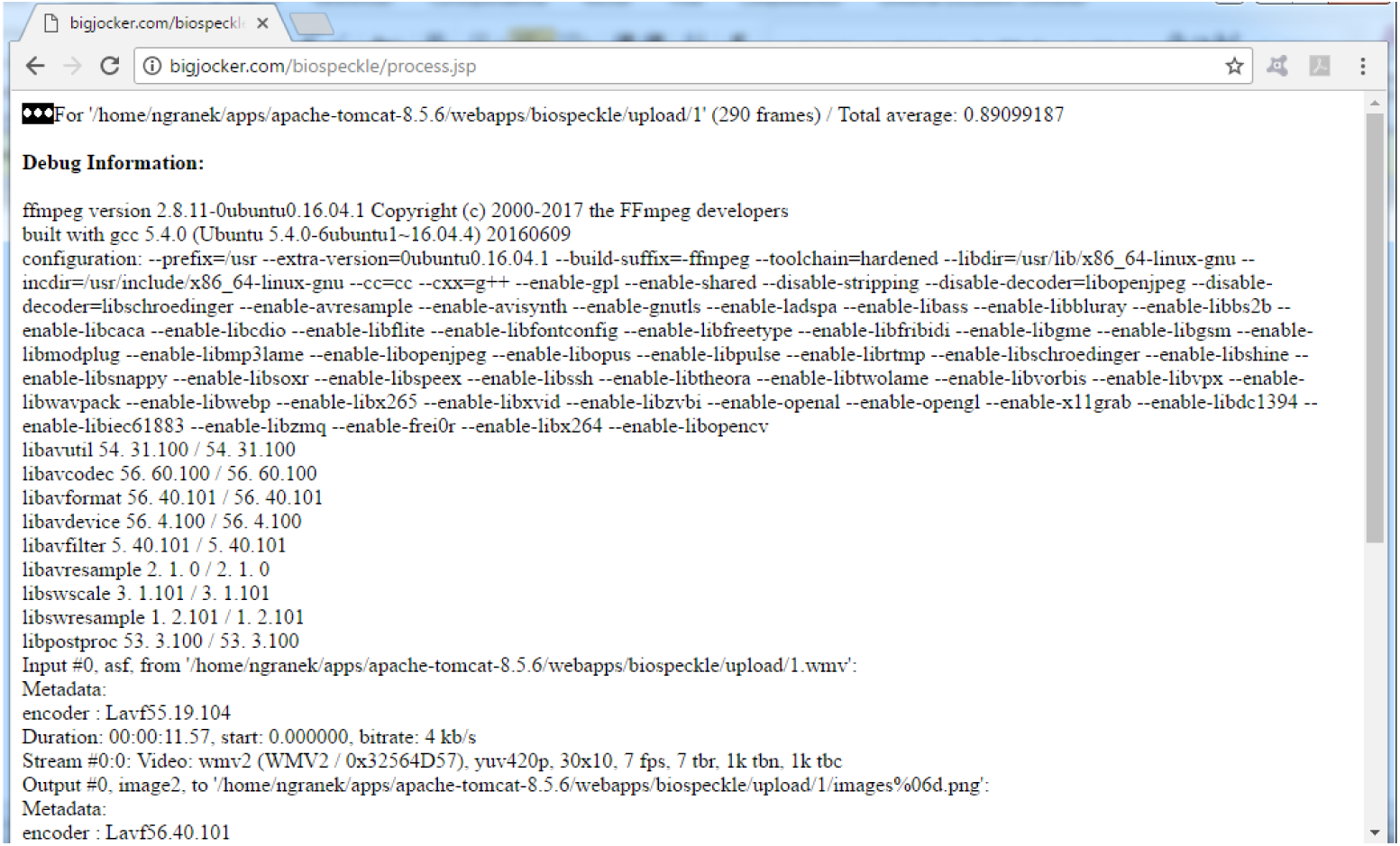
  f. The result will appear in the first line of the page as “Total average:” which in this case is 0.89099187. The rest is Debug Information which should be neglected.

### S2 DATA

Diagnostic statistical tests (linear regression model, normality, homoscedasticity and independence) of the time series for each video

#### Biospeckle laser digital image processing for quantitative and statistical evaluation of the activity of Ciprofloxacin on *Escherichia coli* K-12

Hilda Cristina Grassi, Ana Velásquez, Olga Mercedes Belandria, María Lorena Lobo-Sulbarán, Jesús E. Andrades-Grassi, Humberto Cabrera, Efrén D. J. Andrades

##### Statistical diagnostic tests applied to the time series

###### 1. Adjustment to a linear regression model

| Video | Adjustment to a Linear regression | Video | Adjustment to a Linear regression |
| --- | --- | --- | --- |
| 1 | Call:<br>lm(formula = VIDEO.1 ~ Tiempo, data = Datos)<br>Residuals:<br>Min 1Q Median 3Q Max<br>-117291 -47883 257 36010 137517<br>Coefficients:<br>Estimate Std. Error t value Pr(>\|t\|)<br>(Intercept) 870500 7206 120.807 < 2e-16 ***<br>Tiempo 5100 1490 3.421 0.000727 ***<br>--- | 2 | Call:<br>lm(formula = VIDEO.2 ~ Tiempo, data = Datos)<br>Residuals:<br>Min 1Q Median 3Q Max<br>-320023 -107175 -10803 112645 354472<br>Coefficients:<br>Estimate Std. Error t value Pr(>\|t\|)<br>(Intercept) 913123 19168 47.638 < 2e-16 ***<br>Tiempo 6060 3965 1.529 0.128<br>--- |
|  | Signif. codes: 0 '***' 0.001 '**' 0.01 '*' 0.05 '.' 0.1 ' ' 1<br>Residual standard error: 57140 on 251 degrees of freedom<br>Multiple R-squared: 0.04456, Adjusted R-squared: 0.04075<br>F-statistic: 11.71 on 1 and 251 DF, p-value: 0.000727 |  | Signif. codes: 0 '***' 0.001 '**' 0.01 '*' 0.05 '.' 0.1 ' ' 1<br>Residual standard error: 152000 on 251 degrees of freedom<br>Multiple R-squared: 0.009223, Adjusted R-squared: 0.005276<br>F-statistic: 2.337 on 1 and 251 DF, p-value: 0.1276 |
| 3 | Call:<br>lm(formula = VIDEO.3 ~ Tiempo, data = Datos)<br>Residuals:<br>Min 1Q Median 3Q Max<br>-484725 -244334 -82325 240884 683887<br>Coefficients:<br>Estimate Std. Error t value Pr(>\|t\|)<br>(Intercept) 1270022 41230 30.803 <2e-16 ***<br>Tiempo 9022 8528 1.058 0.291<br>---<br>Signif. codes: 0 '***' 0.001 '**' 0.01 '*' 0.05 '.' 0.1 ' ' 1<br>Residual standard error: 326900 on 251 degrees of freedom<br>Multiple R-squared: 0.004439, Adjusted R-squared: 0.0004723<br>F-statistic: 1.119 on 1 and 251 DF, p-value: 0.2911 | 4 | Call:<br>lm(formula = VIDEO.4 ~ Tiempo, data = Datos)<br>Residuals:<br>Min 1Q Median 3Q Max<br>-320666 -179106 -26064 153914 530574<br>Coefficients:<br>Estimate Std. Error t value Pr(>\|t\|)<br>(Intercept) 1035639 25412 40.754 <2e-16 ***<br>Tiempo 3511 5256 0.668 0.505<br>---<br>Signif. codes: 0 '***' 0.001 '**' 0.01 '*' 0.05 '.' 0.1 ' ' 1<br>Residual standard error: 201500 on 251 degrees of freedom<br>Multiple R-squared: 0.001774, Adjusted R-squared: -0.002203<br>F-statistic: 0.4461 on 1 and 251 DF, p-value: 0.5048 |
| 5 | Call:<br>lm(formula = VIDEO.5 ~ Tiempo, data = Datos)<br>Residuals:<br>Min 1Q Median 3Q Max<br>-155584 -40900 8798 40555 114043<br>Coefficients:<br>Estimate Std. Error t value Pr(>\|t\|)<br>(Intercept) 632324 7450 84.87 <2e-16 ***<br>Tiempo 1495 1541 0.97 0.333<br>---<br>Signif. codes: 0 '***' 0.001 '**' 0.01 '*' 0.05 '.' 0.1 ' ' 1<br>Residual standard error: 59080 on 251 degrees of freedom<br>Multiple R-squared: 0.003734, Adjusted R-squared: -0.000235<br>F-statistic: 0.9408 on 1 and 251 DF, p-value: 0.333 | 6 | Call:<br>lm(formula = VIDEO.6 ~ Tiempo, data = Datos)<br>Residuals:<br>Min 1Q Median 3Q Max<br>-299148 -70251 -10263 81341 515025<br>Coefficients:<br>Estimate Std. Error t value Pr(>\|t\|)<br>(Intercept) 954661 18369 51.972 <2e-16 ***<br>Tiempo 1589 3799 0.418 0.676<br>---<br>Signif. codes: 0 '***' 0.001 '**' 0.01 '*' 0.05 '.' 0.1 ' ' 1<br>Residual standard error: 145700 on 251 degrees of freedom<br>Multiple R-squared: 0.0006962, Adjusted R-squared: -0.003285<br>F-statistic: 0.1749 on 1 and 251 DF, p-value: 0.6762 |
| 7 | Call:<br>lm(formula = VIDEO.7 ~ Tiempo, data = Datos)<br>Residuals:<br>Min 1Q Median 3Q Max<br>-469139 -138508 5276 167822 476378<br>Coefficients:<br>Estimate Std. Error t value Pr(>\|t\|)<br>(Intercept) 3000091 29440 101.907 <2e-16 ***<br>Tiempo -28003 6089 -4.599 6.75e-06 ***<br>---<br>Signif. codes: 0 '***' 0.001 '**' 0.01 '*' 0.05 '.' 0.1 ' ' 1<br>Residual standard error: 233400 on 251 degrees of freedom<br>Multiple R-squared: 0.07771, Adjusted R-squared: 0.07403<br>F-statistic: 21.15 on 1 and 251 DF, p-value: 6.748e-06 | 8 | Call:<br>lm(formula = VIDEO.8 ~ Tiempo, data = Datos)<br>Residuals:<br>Min 1Q Median 3Q Max<br>-234624 -85311 -17019 81990 257430<br>Coefficients:<br>Estimate Std. Error t value Pr(>\|t\|)<br>(Intercept) 972109 14844 65.49 <2e-16 ***<br>Tiempo 3253 3070 1.06 0.29<br>---<br>Signif. codes: 0 '***' 0.001 '**' 0.01 '*' 0.05 '.' 0.1 ' ' 1<br>Residual standard error: 117700 on 251 degrees of freedom<br>Multiple R-squared: 0.004453, Adjusted R-squared: 0.0004868<br>F-statistic: 1.123 on 1 and 251 DF, p-value: 0.2904 |
| 9 | Call:<br>lm(formula = VIDEO.9 ~ Tiempo, data = Datos)<br>Residuals:<br>Min 1Q Median 3Q Max<br>-207184 -76206 -2884 89555 271102<br>Coefficients:<br>Estimate Std. Error t value Pr(>\|t\|)<br>(Intercept) 890773 13552 65.729 <2e-16 ***<br>Tiempo -26961 2803 -9.618 <2e-16 ***<br>---<br>Signif. codes: 0 '***' 0.001 '**' 0.01 '*' 0.05 '.' 0.1 ' ' 1<br>Residual standard error: 107500 on 251 degrees of freedom<br>Multiple R-squared: 0.2693, Adjusted R-squared: 0.2664<br>F-statistic: 92.5 on 1 and 251 DF, p-value: < 2.2e-16 | 10 | Call:<br>lm(formula = VIDEO.10 ~ Tiempo, data = Datos)<br>Residuals:<br>Min 1Q Median 3Q Max<br>-215151 -90130 -2618 81252 256142<br>Coefficients:<br>Estimate Std. Error t value Pr(>\|t\|)<br>(Intercept) 1030770 13566 75.982 <2e-16 ***<br>Tiempo -17638 2806 -6.286 1.43e-09 ***<br>---<br>Signif. codes: 0 '***' 0.001 '**' 0.01 '*' 0.05 '.' 0.1 ' ' 1<br>Residual standard error: 107600 on 251 degrees of freedom<br>Multiple R-squared: 0.136, Adjusted R-squared: 0.1326<br>F-statistic: 39.51 on 1 and 251 DF, p-value: 1.435e-09 |
| 11 | Call:<br>lm(formula = VIDEO.11 ~ Tiempo, data = Datos)<br>Residuals:<br>Min 1Q Median 3Q Max<br>-201703 -112503 -14062 106324 222104<br>Coefficients:<br>Estimate Std. Error t value Pr(>\|t\|)<br>(Intercept) 740001 14743 50.193 <2e-16 ***<br>Tiempo 16898 3050 5.541 7.58e-08 ***<br>--- | 12 | Call:<br>lm(formula = VIDEO.12 ~ Tiempo, data = Datos)<br>Residuals:<br>Min 1Q Median 3Q Max<br>-187122 -56716 -6673 56004 241873<br>Coefficients:<br>Estimate Std. Error t value Pr(>\|t\|)<br>(Intercept) 713284 10412 68.507 <2e-16 ***<br>Tiempo 1630 2154 0.757 0.45<br>--- |
|  | Signif. codes: 0 '***' 0.001 '**' 0.01 '*' 0.05 '.' 0.1 ' ' 1<br>Residual standard error: 116900 on 251 degrees of freedom<br>Multiple R-squared: 0.109, Adjusted R-squared: 0.1055<br>F-statistic: 30.71 on 1 and 251 DF, p-value: 7.576e-08 |  | Signif. codes: 0 '***' 0.001 '**' 0.01 '*' 0.05 '.' 0.1 ' ' 1<br>Residual standard error: 82560 on 251 degrees of freedom<br>Multiple R-squared: 0.002277, Adjusted R-squared: -0.001698<br>F-statistic: 0.5728 on 1 and 251 DF, p-value: 0.4498 |
| 13 | Call:<br>lm(formula = VIDEO.13 ~ Tiempo, data = Datos)<br>Residuals:<br>Min 1Q Median 3Q Max<br>-247192 -81074 -12705 70624 467553<br>Coefficients:<br>Estimate Std. Error t value Pr(>\|t\|)<br>(Intercept) 768335 15235 50.434 < 2e-16 ***<br>Tiempo -9168 3151 -2.909 0.00394 **<br>---<br>Signif. codes: 0 '***' 0.001 '**' 0.01 '*' 0.05 '.' 0.1 ' ' 1<br>Residual standard error: 120800 on 251 degrees of freedom<br>Multiple R-squared: 0.03263, Adjusted R-squared: 0.02877<br>F-statistic: 8.465 on 1 and 251 DF, p-value: 0.003945 | 14 | Call:<br>lm(formula = VIDEO.14 ~ Tiempo, data = Datos)<br>Residuals:<br>Min 1Q Median 3Q Max<br>-225973 -108469 -6417 53522 339881<br>Coefficients:<br>Estimate Std. Error t value Pr(>\|t\|)<br>(Intercept) 849278 16128 52.659 < 2e-16 ***<br>Tiempo 20619 3336 6.181 2.56e-09 ***<br>---<br>Signif. codes: 0 '***' 0.001 '**' 0.01 '*' 0.05 '.' 0.1 ' ' 1<br>Residual standard error: 127900 on 251 degrees of freedom<br>Multiple R-squared: 0.1321, Adjusted R-squared: 0.1286<br>F-statistic: 38.2 on 1 and 251 DF, p-value: 2.565e-09 |
| 15 | Call:<br>lm(formula = VIDEO.15 ~ Tiempo, data = Datos)<br>Residuals:<br>Min 1Q Median 3Q Max<br>-215863 -92612 -11438 99087 312536<br>Coefficients:<br>Estimate Std. Error t value Pr(>\|t\|)<br>(Intercept) 736779 16299 45.203 < 2e-16 ***<br>Tiempo 22278 3371 6.608 2.32e-10 ***<br>---<br>Signif. codes: 0 '***' 0.001 '**' 0.01 '*' 0.05 '.' 0.1 ' ' 1<br>Residual standard error: 129200 on 251 degrees of freedom<br>Multiple R-squared: 0.1482, Adjusted R-squared: 0.1448<br>F-statistic: 43.67 on 1 and 251 DF, p-value: 2.318e-10 | 16 | Call:<br>lm(formula = VIDEO.16 ~ Tiempo, data = Datos)<br>Residuals:<br>Min 1Q Median 3Q Max<br>-392116 -193878 -20345 191095 453841<br>Coefficients:<br>Estimate Std. Error t value Pr(>\|t\|)<br>(Intercept) 991507 28229 35.124 < 2e-16 ***<br>Tiempo 21500 5839 3.682 0.000283 ***<br>---<br>Signif. codes: 0 '***' 0.001 '**' 0.01 '*' 0.05 '.' 0.1 ' ' 1<br>Residual standard error: 223800 on 251 degrees of freedom<br>Multiple R-squared: 0.05125, Adjusted R-squared: 0.04747<br>F-statistic: 13.56 on 1 and 251 DF, p-value: 0.0002831 |
| 17 | Call:<br>lm(formula = VIDEO.17 ~ Tiempo, data = Datos)<br>Residuals:<br>Min 1Q Median 3Q Max<br>-281351 -95042 -34888 87388 429148<br>Coefficients:<br>Estimate Std. Error t value Pr(>\|t\|)<br>(Intercept) 2448303 18507 132.291 < 2e-16 ***<br>Tiempo 6792 3828 1.774 0.0772 .<br>---<br>Signif. codes: 0 '***' 0.001 '**' 0.01 '*' 0.05 '.' 0.1 ' ' 1<br>Residual standard error: 146700 on 251 degrees of freedom<br>Multiple R-squared: 0.01239, Adjusted R-squared: 0.008453<br>F-statistic: 3.148 on 1 and 251 DF, p-value: 0.07721 | 18 | Call:<br>lm(formula = VIDEO.18 ~ Tiempo, data = Datos)<br>Residuals:<br>Min 1Q Median 3Q Max<br>-401096 -100602 -10143 98088 324063<br>Coefficients:<br>Estimate Std. Error t value Pr(>\|t\|)<br>(Intercept) 1150475 19629 58.612 < 2e-16 ***<br>Tiempo -33620 4060 -8.281 7.35e-15 ***<br>---<br>Signif. codes: 0 '***' 0.001 '**' 0.01 '*' 0.05 '.' 0.1 ' ' 1<br>Residual standard error: 155600 on 251 degrees of freedom<br>Multiple R-squared: 0.2146, Adjusted R-squared: 0.2114<br>F-statistic: 68.57 on 1 and 251 DF, p-value: 7.347e-15 |
| 19 | Call:<br>lm(formula = VIDEO.19 ~ Tiempo, data = Datos)<br>Residuals:<br>Min 1Q Median 3Q Max<br>-203082 -104867 1109 91236 244976<br>Coefficients:<br>Estimate Std. Error t value Pr(>\|t\|)<br>(Intercept) 701284 15208 46.112 < 2e-16 ***<br>Tiempo 17069 3146 5.426 1.36e-07 ***<br>---<br>Signif. codes: 0 '***' 0.001 '**' 0.01 '*' 0.05 '.' 0.1 ' ' 1<br>Residual standard error: 120600 on 251 degrees of freedom<br>Multiple R-squared: 0.105, Adjusted R-squared: 0.1014<br>F-statistic: 29.44 on 1 and 251 DF, p-value: 1.357e-07 | 20 | Call:<br>lm(formula = VIDEO.20 ~ Tiempo, data = Datos)<br>Residuals:<br>Min 1Q Median 3Q Max<br>-307949 -93716 -41513 58144 493700<br>Coefficients:<br>Estimate Std. Error t value Pr(>\|t\|)<br>(Intercept) 1056966 20688 51.091 < 2e-16 ***<br>Tiempo 1859 4279 0.434 0.664<br>---<br>Signif. codes: 0 '***' 0.001 '**' 0.01 '*' 0.05 '.' 0.1 ' ' 1<br>Residual standard error: 164000 on 251 degrees of freedom<br>Multiple R-squared: 0.0007511, Adjusted R-squared: -0.00323<br>F-statistic: 0.1887 on 1 and 251 DF, p-value: 0.6644 |
| 21 | Call:<br>lm(formula = VIDEO.21 ~ Tiempo, data = Datos)<br>Residuals:<br>Min 1Q Median 3Q Max<br>-125832 -66668 -22066 71099 169179<br>Coefficients:<br>Estimate Std. Error t value Pr(>\|t\|)<br>(Intercept) 774871 9912 78.176 < 2e-16 ***<br>Tiempo 7472 2050 3.644 0.000326 ***<br>--- | 22 | Call:<br>lm(formula = VIDEO.22 ~ Tiempo, data = Datos)<br>Residuals:<br>Min 1Q Median 3Q Max<br>-216235 -74359 -15305 62223 267010<br>Coefficients:<br>Estimate Std. Error t value Pr(>\|t\|)<br>(Intercept) 780523 12413 62.879 < 2e-16 ***<br>Tiempo 492 2568 0.192 0.848<br>--- |
|  | Signif. codes: 0 '***' 0.001 '**' 0.01 '*' 0.05 '.' 0.1 ' ' 1<br>Residual standard error: 78600 on 251 degrees of freedom<br>Multiple R-squared: 0.05026, Adjusted R-squared: 0.04647<br>F-statistic: 13.28 on 1 and 251 DF, p-value: 0.0003257 |  | Signif. codes: 0 '***' 0.001 '**' 0.01 '*' 0.05 '.' 0.1 ' ' 1<br>Residual standard error: 98430 on 251 degrees of freedom<br>Multiple R-squared: 0.0001462, Adjusted R-squared: -0.003837<br>F-statistic: 0.03671 on 1 and 251 DF, p-value: 0.8482 |
| 23 | Call:<br>lm(formula = VIDEO.23 ~ Tiempo, data = Datos)<br>Residuals:<br>Min 1Q Median 3Q Max<br>-272821 -103461 4080 91155 288324<br>Coefficients:<br>Estimate Std. Error t value Pr(>\|t\|)<br>(Intercept) 998278.8 15025.1 66.44 <2e-16 ***<br>Tiempo 964.6 3107.8 0.31 0.757<br>---<br>Signif. codes: 0 '***' 0.001 '**' 0.01 '*' 0.05 '.' 0.1 ' ' 1<br>Residual standard error: 119100 on 251 degrees of freedom<br>Multiple R-squared: 0.0003837, Adjusted R-squared: -0.003599<br>F-statistic: 0.09633 on 1 and 251 DF, p-value: 0.7565 | 24 | Call:<br>lm(formula = VIDEO.24 ~ Tiempo, data = Datos)<br>Residuals:<br>Min 1Q Median 3Q Max<br>-325966 -132484 -2550 128486 288538<br>Coefficients:<br>Estimate Std. Error t value Pr(>\|t\|)<br>(Intercept) 1261346 17863 70.612 <2e-16 ***<br>Tiempo -6272 3695 -1.698 0.0908<br>---<br>Signif. codes: 0 '***' 0.001 '**' 0.01 '*' 0.05 '.' 0.1 ' ' 1<br>Residual standard error: 141600 on 251 degrees of freedom<br>Multiple R-squared: 0.01135, Adjusted R-squared: 0.007411<br>F-statistic: 2.882 on 1 and 251 DF, p-value: 0.09084 |
| 25 | Call:<br>lm(formula = VIDEO.25 ~ Tiempo, data = Datos)<br>Residuals:<br>Min 1Q Median 3Q Max<br>-425023 -199757 -87238 122518 535580<br>Coefficients:<br>Estimate Std. Error t value Pr(>\|t\|)<br>(Intercept) 1033636 32607 31.700 <2e-16 ***<br>Tiempo 30365 6744 4.502 1.03e-05 ***<br>---<br>Signif. codes: 0 '***' 0.001 '**' 0.01 '*' 0.05 '.' 0.1 ' ' 1<br>Residual standard error: 258600 on 251 degrees of freedom<br>Multiple R-squared: 0.07472, Adjusted R-squared: 0.07104<br>F-statistic: 20.27 on 1 and 251 DF, p-value: 1.03e-05 | 26 | Call:<br>lm(formula = VIDEO.26 ~ Tiempo, data = Datos)<br>Residuals:<br>Min 1Q Median 3Q Max<br>-235068 -120308 -7730 77496 434569<br>Coefficients:<br>Estimate Std. Error t value Pr(>\|t\|)<br>(Intercept) 1468574 18523 79.284 <2e-16 ***<br>Tiempo 10003 3831 2.611 0.00957 **<br>---<br>Signif. codes: 0 '***' 0.001 '**' 0.01 '*' 0.05 '.' 0.1 ' ' 1<br>Residual standard error: 146900 on 251 degrees of freedom<br>Multiple R-squared: 0.02644, Adjusted R-squared: 0.02256<br>F-statistic: 6.817 on 1 and 251 DF, p-value: 0.009573 |
| 27 | Call:<br>lm(formula = VIDEO.27 ~ Tiempo, data = Datos)<br>Residuals:<br>Min 1Q Median 3Q Max<br>-194954 -65088 -3376 53535 286076<br>Coefficients:<br>Estimate Std. Error t value Pr(>\|t\|)<br>(Intercept) 955504 12433 76.854 <2e-16 ***<br>Tiempo 7126 2572 2.771 0.00601 **<br>---<br>Signif. codes: 0 '***' 0.001 '**' 0.01 '*' 0.05 '.' 0.1 ' ' 1<br>Residual standard error: 98580 on 251 degrees of freedom<br>Multiple R-squared: 0.02968, Adjusted R-squared: 0.02582<br>F-statistic: 7.678 on 1 and 251 DF, p-value: 0.006009 | 28 | Call:<br>lm(formula = VIDEO.28 ~ Tiempo, data = Datos)<br>Residuals:<br>Min 1Q Median 3Q Max<br>-184035 -89483 959 82720 268953<br>Coefficients:<br>Estimate Std. Error t value Pr(>\|t\|)<br>(Intercept) 993360 13417 74.038 <2e-16 ***<br>Tiempo -12652 2775 -4.559 8.04e-06 ***<br>---<br>Signif. codes: 0 '***' 0.001 '**' 0.01 '*' 0.05 '.' 0.1 ' ' 1<br>Residual standard error: 106400 on 251 degrees of freedom<br>Multiple R-squared: 0.07647, Adjusted R-squared: 0.07279<br>F-statistic: 20.78 on 1 and 251 DF, p-value: 8.038e-06 |
| 29 | Call:<br>lm(formula = VIDEO.29 ~ Tiempo, data = Datos)<br>Residuals:<br>Min 1Q Median 3Q Max<br>-422848 -143847 30505 123099 433622<br>Coefficients:<br>Estimate Std. Error t value Pr(>\|t\|)<br>(Intercept) 1514064 26777 56.543 <2e-16 ***<br>Tiempo 6674 5539 1.205 0.229<br>---<br>Signif. codes: 0 '***' 0.001 '**' 0.01 '*' 0.05 '.' 0.1 ' ' 1<br>Residual standard error: 212300 on 251 degrees of freedom<br>Multiple R-squared: 0.005752, Adjusted R-squared: 0.001791<br>F-statistic: 1.452 on 1 and 251 DF, p-value: 0.2293 | 30 | Call:<br>lm(formula = VIDEO.30 ~ Tiempo, data = Datos)<br>Residuals:<br>Min 1Q Median 3Q Max<br>-185409 -32568 13032 41810 132576<br>Coefficients:<br>Estimate Std. Error t value Pr(>\|t\|)<br>(Intercept) 640121 7859 81.45 <2e-16 ***<br>Tiempo 1463 1626 0.90 0.369<br>---<br>Signif. codes: 0 '***' 0.001 '**' 0.01 '*' 0.05 '.' 0.1 ' ' 1<br>Residual standard error: 62310 on 251 degrees of freedom<br>Multiple R-squared: 0.003218, Adjusted R-squared: -0.0007534<br>F-statistic: 0.8103 on 1 and 251 DF, p-value: 0.3689 |
| 31 | Call:<br>lm(formula = VIDEO.31 ~ Tiempo, data = Datos)<br>Residuals:<br>Min 1Q Median 3Q Max<br>-258323 -98362 -25144 91199 358395<br>Coefficients:<br>Estimate Std. Error t value Pr(>\|t\|)<br>(Intercept) 1013436 17534 57.800 <2e-16 ***<br>Tiempo 22032 3627 6.075 4.57e-09 ***<br>--- | 32 | Call:<br>lm(formula = VIDEO.32 ~ Tiempo, data = Datos)<br>Residuals:<br>Min 1Q Median 3Q Max<br>-547680 -268713 -101728 273029 788075<br>Coefficients:<br>Estimate Std. Error t value Pr(>\|t\|)<br>(Intercept) 1111654 44768 24.832 <2e-16 ***<br>Tiempo 24859 9260 2.685 0.00775 **<br>--- |
|  | Signif. codes: 0 '***' 0.001 '**' 0.01 '*' 0.05 '.' 0.1 ' ' 1<br>Residual standard error: 139000 on 251 degrees of freedom<br>Multiple R-squared: 0.1282, Adjusted R-squared: 0.1247<br>F-statistic: 36.9 on 1 and 251 DF, p-value: 4.573e-09 |  | Signif. codes: 0 '***' 0.001 '**' 0.01 '*' 0.05 '.' 0.1 ' ' 1<br>Residual standard error: 355000 on 251 degrees of freedom<br>Multiple R-squared: 0.02791, Adjusted R-squared: 0.02404<br>F-statistic: 7.207 on 1 and 251 DF, p-value: 0.007746 |
| 33 | Call:<br>lm(formula = VIDEO.33 ~ Tiempo, data = Datos)<br>Residuals:<br>Min 1Q Median 3Q Max<br>-418928 -233410 66114 212111 328689<br>Coefficients:<br>Estimate Std. Error t value Pr(>\|t\|)<br>(Intercept) 1118498 29850 37.471 <2e-16 ***<br>Tiempo 4144 6174 0.671 0.503<br>---<br>Signif. codes: 0 '***' 0.001 '**' 0.01 '*' 0.05 '.' 0.1 ' ' 1<br>Residual standard error: 236700 on 251 degrees of freedom<br>Multiple R-squared: 0.001791, Adjusted R-squared: -0.002186<br>F-statistic: 0.4504 on 1 and 251 DF, p-value: 0.5027 | 34 | Call:<br>lm(formula = VIDEO.34 ~ Tiempo, data = Datos)<br>Residuals:<br>Min 1Q Median 3Q Max<br>-353018 -151476 2586 127746 470894<br>Coefficients:<br>Estimate Std. Error t value Pr(>\|t\|)<br>(Intercept) 903587 23931 37.757 <2e-16 ***<br>Tiempo 46212 4950 9.336 <2e-16 ***<br>---<br>Signif. codes: 0 '***' 0.001 '**' 0.01 '*' 0.05 '.' 0.1 ' ' 1<br>Residual standard error: 189800 on 251 degrees of freedom<br>Multiple R-squared: 0.2577, Adjusted R-squared: 0.2548<br>F-statistic: 87.15 on 1 and 251 DF, p-value: < 2.2e-16 |
| 35 | Call:<br>lm(formula = VIDEO.35 ~ Tiempo, data = Datos)<br>Residuals:<br>Min 1Q Median 3Q Max<br>-214728 -119736 -13382 106921 286451<br>Coefficients:<br>Estimate Std. Error t value Pr(>\|t\|)<br>(Intercept) 799448 16606 48.142 < 2e-16 ***<br>Tiempo 13476 3435 3.923 0.000113 ***<br>---<br>Signif. codes: 0 '***' 0.001 '**' 0.01 '*' 0.05 '.' 0.1 ' ' 1<br>Residual standard error: 131700 on 251 degrees of freedom<br>Multiple R-squared: 0.05778, Adjusted R-squared: 0.05403<br>F-statistic: 15.39 on 1 and 251 DF, p-value: 0.0001128 | 36 | Call:<br>lm(formula = VIDEO.36 ~ Tiempo, data = Datos)<br>Residuals:<br>Min 1Q Median 3Q Max<br>-325899 -119607 -33302 101673 467356<br>Coefficients:<br>Estimate Std. Error t value Pr(>\|t\|)<br>(Intercept) 909358 20550 44.251 < 2e-16 ***<br>Tiempo 17755 4251 4.177 4.08e-05 ***<br>---<br>Signif. codes: 0 '***' 0.001 '**' 0.01 '*' 0.05 '.' 0.1 ' ' 1<br>Residual standard error: 162900 on 251 degrees of freedom<br>Multiple R-squared: 0.065, Adjusted R-squared: 0.06127<br>F-statistic: 17.45 on 1 and 251 DF, p-value: 4.077e-05 |
| Control A | Call:<br>lm(formula = CONT.2 ~ Tiempo, data = Datos)<br>Residuals:<br>Min 1Q Median 3Q Max<br>-290321 -92546 -7409 79482 314977<br>Coefficients:<br>Estimate Std. Error t value Pr(>\|t\|)<br>(Intercept) 863406 15526 55.611 <2e-16 ***<br>Tiempo 6739 3211 2.098 0.0369 *<br>---<br>Signif. codes: 0 '***' 0.001 '**' 0.01 '*' 0.05 '.' 0.1 ' ' 1<br>Residual standard error: 123100 on 251 degrees of freedom<br>Multiple R-squared: 0.01724, Adjusted R-squared: 0.01333<br>F-statistic: 4.403 on 1 and 251 DF, p-value: 0.03687 | Control B | Call:<br>lm(formula = CONT.4 ~ Tiempo, data = Datos)<br>Residuals:<br>Min 1Q Median 3Q Max<br>-133866 -40715 -9433 30951 210560<br>Coefficients:<br>Estimate Std. Error t value Pr(>\|t\|)<br>(Intercept) 705176 9367 75.285 <2e-16 ***<br>Tiempo 1688 1937 0.871 0.385<br>---<br>Signif. codes: 0 '***' 0.001 '**' 0.01 '*' 0.05 '.' 0.1 ' ' 1<br>Residual standard error: 74270 on 251 degrees of freedom<br>Multiple R-squared: 0.003013, Adjusted R-squared: -0.0009587<br>F-statistic: 0.7586 on 1 and 251 DF, p-value: 0.3846 |
| Control C | Call:<br>lm(formula = CONT.1 ~ Tiempo, data = Datos)<br>Residuals:<br>Min 1Q Median 3Q Max<br>-354556 -229332 72075 178169 336171<br>Coefficients:<br>Estimate Std. Error t value Pr(>\|t\|)<br>(Intercept) 1030659 26170 39.383 < 2e-16 ***<br>Tiempo 26644 5413 4.922 1.55e-06 ***<br>---<br>Signif. codes: 0 '***' 0.001 '**' 0.01 '*' 0.05 '.' 0.1 ' ' 1<br>Residual standard error: 207500 on 251 degrees of freedom<br>Multiple R-squared: 0.08803, Adjusted R-squared: 0.08439<br>F-statistic: 24.23 on 1 and 251 DF, p-value: 1.552e-06 | Control D | Call:<br>lm(formula = CONT.3 ~ Tiempo, data = Datos)<br>Residuals:<br>Min 1Q Median 3Q Max<br>-256615 -60802 9944 74501 202321<br>Coefficients:<br>Estimate Std. Error t value Pr(>\|t\|)<br>(Intercept) 878516 12726 69.032 < 2e-16 ***<br>Tiempo -11988 2632 -4.554 8.21e-06 ***<br>---<br>Signif. codes: 0 '***' 0.001 '**' 0.01 '*' 0.05 '.' 0.1 ' ' 1<br>Residual standard error: 100900 on 251 degrees of freedom<br>Multiple R-squared: 0.07632, Adjusted R-squared: 0.07264<br>F-statistic: 20.74 on 1 and 251 DF, p-value: 8.214e-06 |

###### 2. Shapiro-Wilk normality test

| Video | Shapiro-Wilk normality test | Video | Shapiro-Wilk normality test |
| --- | --- | --- | --- |
| 1 | Shapiro-Wilk normality test<br>data: Datos\$VIDEO.1<br>W = 0.8603, p-value = 2.978e-05 | 2 | Shapiro-Wilk normality test<br>data: Datos\$VIDEO.2<br>W = 0.8812, p-value = 0.0001217 |
| 3 | Shapiro-Wilk normality test<br>data: Datos\$VIDEO.2<br>W = 0.8812, p-value = 0.0001217 | 4 | Shapiro-Wilk normality test<br>data: Datos\$VIDEO.4<br>W = 0.9209, p-value = 0.002532 |
| 5 | Shapiro-Wilk normality test<br>data: Datos\$VIDEO.5<br>W = 0.9147, p-value = 0.001529 | 6 | Shapiro-Wilk normality test<br>data: Datos\$VIDEO.6<br>W = 0.9103, p-value = 0.001075 |
| 7 | Shapiro-Wilk normality test<br>data: Datos\$VIDEO.7<br>W = 0.9069, p-value = 0.0008211 | 8 | Shapiro-Wilk normality test<br>data: Datos\$VIDEO.8<br>W = 0.9743, p-value = 0.3432 |
| 9 | Shapiro-Wilk normality test<br>data: Datos\$VIDEO.9<br>W = 0.7441, p-value = 5.607e-08 | 10 | Shapiro-Wilk normality test<br>data: Datos\$VIDEO.10<br>W = 0.8447, p-value = 1.113e-05 |
| 11 | Shapiro-Wilk normality test<br>data: Datos\$VIDEO.11<br>W = 0.9157, p-value = 0.00165 | 12 | Shapiro-Wilk normality test<br>data: Datos\$VIDEO.12<br>W = 0.9495, p-value = 0.03252 |
| 13 | Shapiro-Wilk normality test<br>data: Datos\$VIDEO.13<br>W = 0.8085, p-value = 1.381e-06 | 14 | Shapiro-Wilk normality test<br>data: Datos\$VIDEO.14<br>W = 0.9151, p-value = 0.001572 |
| 15 | Shapiro-Wilk normality test<br>data: Datos\$VIDEO.15<br>W = 0.9288, p-value = 0.004983 | 16 | Shapiro-Wilk normality test<br>data: Datos\$VIDEO.16<br>W = 0.9396, p-value = 0.01294 |
| 17 | Shapiro-Wilk normality test<br>data: Datos\$VIDEO.17<br>W = 0.857, p-value = 2.406e-05 | 18 | Shapiro-Wilk normality test<br>data: Datos\$VIDEO.18<br>W = 0.8948, p-value = 0.0003252 |
| 19 | Shapiro-Wilk normality test<br>data: Datos\$VIDEO.19<br>W = 0.9362, p-value = 0.009521 | 20 | Shapiro-Wilk normality test<br>data: Datos\$VIDEO.20<br>W = 0.8847, p-value = 0.0001557 |
| 21 | Shapiro-Wilk normality test<br>data: Datos\$VIDEO.21<br>W = 0.8719, p-value = 6.391e-05 | 22 | Shapiro-Wilk normality test<br>data: Datos\$VIDEO.22<br>W = 0.773, p-value = 2.211e-07 |
| 23 | Shapiro-Wilk normality test<br>data: Datos\$VIDEO.23<br>W = 0.91, p-value = 0.001043 | 24 | Shapiro-Wilk normality test<br>data: Datos\$VIDEO.24<br>W = 0.9576, p-value = 0.07036 |
| 25 | Shapiro-Wilk normality test<br>data: Datos\$VIDEO.25<br>W = 0.8596, p-value = 2.849e-05 | 26 | Shapiro-Wilk normality test<br>data: Datos\$VIDEO.26<br>W = 0.9077, p-value = 0.0008721 |
| 27 | Shapiro-Wilk normality test<br>data: Datos\$VIDEO.27<br>W = 0.9373, p-value = 0.01058 | 28 | Shapiro-Wilk normality test<br>data: Datos\$VIDEO.28<br>W = 0.9547, p-value = 0.05754 |
| 29 | Shapiro-Wilk normality test<br>data: Datos\$VIDEO.29<br>W = 0.9241, p-value = 0.003318 | 30 | Shapiro-Wilk normality test<br>data: Datos\$VIDEO.30<br>W = 0.9555, p-value = 0.0577 |
| 31 | Shapiro-Wilk normality test<br>data: Datos\$VIDEO.31<br>W = 0.9055, p-value = 0.0007347 | 32 | Shapiro-Wilk normality test<br>data: Datos\$VIDEO.32<br>W = 0.9703, p-value = 0.2385 |
| 33 | Shapiro-Wilk normality test<br>data: Datos\$VIDEO.33<br>W = 0.8161, p-value = 2.106e-06 | 34 | Shapiro-Wilk normality test<br>data: Datos\$VIDEO.34<br>W = 0.9633, p-value = 0.1218 |
| 35 | Shapiro-Wilk normality test<br>data: Datos\$VIDEO.35<br>W = 0.8904, p-value = 0.0002354 | 36 | Shapiro-Wilk normality test<br>data: Datos\$VIDEO.36<br>W = 0.7825, p-value = 3.544e-07 |
| Control A | Shapiro-Wilk normality test<br>data: Datos\$CONT.2<br>W = 0.8752, p-value = 8.012e-05 | Control B | Shapiro-Wilk normality test<br>data: Datos\$CONT.4<br>W = 0.9746, p-value = 0.3528 |
| Control<br>C | Shapiro-Wilk normality test<br>data: Datos\$CONT.1<br>W = 0.8618, p-value = 3.282e-05 | Control<br>D | Shapiro-Wilk normality test<br>data: Datos\$CONT.3<br>W = 0.9066, p-value = 0.0007993 |

###### 3. Breusch-Pagan homoscedasticity/heteroscedasticity test

**3. Breusch-Pagan homoscedasticity/heteroscedasticity test.**
| Video | Breusch-Pagan test | Video | Breusch-Pagan test |
| --- | --- | --- | --- |
| 1 | Breusch-Pagan test<br>data: VIDEO.1 ~ Tiempo<br>BP = 0.4847, df = 1, p-value = 0.4863 | 2 | Breusch-Pagan test<br>data: VIDEO.2 ~ Tiempo<br>BP = 27.007, df = 1, p-value = 2.027e-07 |
| 3 | Breusch-Pagan test<br>data: VIDEO.3 ~ Tiempo<br>BP = 4.5653, df = 1, p-value = 0.03263 H0 | 4 | Breusch-Pagan test<br>data: VIDEO.4 ~ Tiempo<br>BP = 6.3092, df = 1, p-value = 0.01201 |
| 5 | Breusch-Pagan test<br>data: VIDEO.5 ~ Tiempo<br>BP = 0.0042, df = 1, p-value = 0.9484 H0 | 6 | Breusch-Pagan test<br>data: VIDEO.6 ~ Tiempo<br>BP = 4.8933, df = 1, p-value = 0.02696 |
| 7 | Breusch-Pagan test<br>data: VIDEO.7 ~ Tiempo<br>BP = 0.4104, df = 1, p-value = 0.5217 H0 | 8 | Breusch-Pagan test<br>data: VIDEO.8 ~ Tiempo<br>BP = 1.3417, df = 1, p-value = 0.2467 |
| 9 | Breusch-Pagan test<br>data: VIDEO.9 ~ Tiempo<br>BP = 2.7262, df = 1, p-value = 0.09871 H0 | 10 | Breusch-Pagan test<br>data: VIDEO.10 ~ Tiempo<br>BP = 9.5423, df = 1, p-value = 0.002008 |
| 11 | Breusch-Pagan test<br>data: VIDEO.11 ~ Tiempo<br>BP = 0.556, df = 1, p-value = 0.4559 H0 | 12 | Breusch-Pagan test<br>data: VIDEO.12 ~ Tiempo<br>BP = 4.4931, df = 1, p-value = 0.03403 |
| 13 | Breusch-Pagan test<br>data: VIDEO.13 ~ Tiempo<br>BP = 49.5292, df = 1, p-value = 1.954e-12 | 14 | Breusch-Pagan test<br>data: VIDEO.14 ~ Tiempo<br>BP = 0.3135, df = 1, p-value = 0.5755 H0 |
| 15 | Breusch-Pagan test<br>data: VIDEO.15 ~ Tiempo<br>BP = 2.4645, df = 1, p-value = 0.1164 H0 | 16 | Breusch-Pagan test<br>data: VIDEO.16 ~ Tiempo<br>BP = 25.9506, df = 1, p-value = 3.503e-07 |
| 17 | Breusch-Pagan test<br>data: VIDEO.17 ~ Tiempo<br>BP = 17.1702, df = 1, p-value = 3.417e-05 | 18 | Breusch-Pagan test<br>data: VIDEO.18 ~ Tiempo<br>BP = 39.6826, df = 1, p-value = 2.988e-10 |
| 19 | Breusch-Pagan test<br>data: VIDEO.19 ~ Tiempo | 20 | Breusch-Pagan test<br>data: VIDEO.20 ~ Tiempo |
|  | BP = 12.5146, df = 1, p-value = 0.0004038 |  | BP = 1.8524, df = 1, p-value = 0.1735 H0 |
| 21 | Breusch-Pagan test<br>data: VIDEO.21 ~ Tiempo<br>BP = 0.8997, df = 1, p-value = 0.3429 H0 | 22 | Breusch-Pagan test<br>data: VIDEO.22 ~ Tiempo<br>BP = 6.0733, df = 1, p-value = 0.01372 |
| 23 | Breusch-Pagan test<br>data: VIDEO.23 ~ Tiempo<br>BP = 0.6228, df = 1, p-value = 0.43 H0 | 24 | Breusch-Pagan test<br>data: VIDEO.24 ~ Tiempo<br>BP = 0.0621, df = 1, p-value = 0.8032 H0 |
| 25 | Breusch-Pagan test<br>data: VIDEO.25 ~ Tiempo<br>BP = 0.0257, df = 1, p-value = 0.8726 H0 | 26 | Breusch-Pagan test<br>data: VIDEO.26 ~ Tiempo<br>BP = 6.7467, df = 1, p-value = 0.009392 |
| 27 | Breusch-Pagan test<br>data: VIDEO.27 ~ Tiempo<br>BP = 0.3094, df = 1, p-value = 0.578 H0 | 28 | Breusch-Pagan test<br>data: VIDEO.28 ~ Tiempo<br>BP = 12.6903, df = 1, p-value = 0.0003676 |
| 29 | Breusch-Pagan test<br>data: VIDEO.29 ~ Tiempo<br>BP = 2.4715, df = 1, p-value = 0.1159 H0 | 30 | Breusch-Pagan test<br>data: VIDEO.30 ~ Tiempo<br>BP = 4.1197, df = 1, p-value = 0.04239 |
| 31 | Breusch-Pagan test<br>data: VIDEO.31 ~ Tiempo<br>BP = 5.373, df = 1, p-value = 0.02045 | 32 | Breusch-Pagan test<br>data: VIDEO.32 ~ Tiempo<br>BP = 15.0819, df = 1, p-value = 0.0001029 |
| 33 | Breusch-Pagan test<br>data: VIDEO.33 ~ Tiempo<br>BP = 0.0696, df = 1, p-value = 0.7919 H0 | 34 | Breusch-Pagan test<br>data: VIDEO.34 ~ Tiempo<br>BP = 0.187, df = 1, p-value = 0.6654 H0 |
| 35 | Breusch-Pagan test<br>data: VIDEO.35 ~ Tiempo<br>BP = 0.8886, df = 1, p-value = 0.3459 H0 | 36 | Breusch-Pagan test<br>data: VIDEO.36 ~ Tiempo<br>BP = 22.0248, df = 1, p-value = 2.691e-06 |
| Contr<br>ol A | Breusch-Pagan test<br>data: CONT.2 ~ Tiempo<br>BP = 1.2233, df = 1, p-value = 0.2687 H0 | Control<br>B | Breusch-Pagan test<br>data: CONT.4 ~ Tiempo<br>BP = 10.7095, df = 1, p-value = 0.001066 |
| Contr<br>ol C | Breusch-Pagan test<br>data: CONT.1 ~ Tiempo<br>BP = 5.2008, df = 1, p-value = 0.02258 | Control<br>D | Breusch-Pagan test<br>data: CONT.3 ~ Tiempo<br>BP = 30.2329, df = 1, p-value = 3.831e-08 |

###### 4. Durbin-Watson independence test

4. Durbin-Watson independence test.
| Video | Durbin-Watson test | Video | Durbin-Watson test |
| --- | --- | --- | --- |
| 1 | Durbin-Watson test<br>data: VIDEO.1 ~ Tiempo<br>DW = 0.399, p-value < 2.2e-16 | 2 | Durbin-Watson test<br>data: VIDEO.2 ~ Tiempo<br>DW = 0.435, p-value < 2.2e-16 |
|  | alternative hypothesis: true autocorrelation is greater than 0 |  | alternative hypothesis: true autocorrelation is greater than 0 |
| 3 | Durbin-Watson test<br>data: VIDEO.3 ~ Tiempo<br>DW = 0.2724, p-value < 2.2e-16<br>alternative hypothesis: true autocorrelation is greater than 0 | 4 | Durbin-Watson test<br>data: VIDEO.4 ~ Tiempo<br>DW = 0.3471, p-value < 2.2e-16<br>alternative hypothesis: true autocorrelation is greater than 0 |
| 5 | Durbin-Watson test<br>data: VIDEO.5 ~ Tiempo<br>DW = 0.418, p-value < 2.2e-16<br>alternative hypothesis: true autocorrelation is greater than 0 | 6 | Durbin-Watson test<br>data: VIDEO.6 ~ Tiempo<br>DW = 0.5456, p-value < 2.2e-16<br>alternative hypothesis: true autocorrelation is greater than 0 |
| 7 | Durbin-Watson test<br>data: VIDEO.7 ~ Tiempo<br>DW = 0.3796, p-value < 2.2e-16<br>alternative hypothesis: true autocorrelation is greater than 0 | 8 | Durbin-Watson test<br>data: VIDEO.8 ~ Tiempo<br>DW = 0.604, p-value < 2.2e-16<br>alternative hypothesis: true autocorrelation is greater than 0 |
| 9 | Durbin-Watson test<br>data: VIDEO.9 ~ Tiempo<br>DW = 0.4568, p-value < 2.2e-16<br>alternative hypothesis: true autocorrelation is greater than 0 | 10 | Durbin-Watson test<br>data: VIDEO.10 ~ Tiempo<br>DW = 0.3467, p-value < 2.2e-16<br>alternative hypothesis: true autocorrelation is greater than 0 |
| 11 | Durbin-Watson test<br>data: VIDEO.11 ~ Tiempo<br>DW = 0.5297, p-value < 2.2e-16<br>alternative hypothesis: true autocorrelation is greater than 0 | 12 | Durbin-Watson test<br>data: VIDEO.12 ~ Tiempo<br>DW = 0.4708, p-value < 2.2e-16<br>alternative hypothesis: true autocorrelation is greater than 0 |
| 13 | Durbin-Watson test<br>data: VIDEO.13 ~ Tiempo<br>DW = 0.5147, p-value < 2.2e-16<br>alternative hypothesis: true autocorrelation is greater than 0 | 14 | Durbin-Watson test<br>data: VIDEO.14 ~ Tiempo<br>DW = 0.4507, p-value < 2.2e-16<br>alternative hypothesis: true autocorrelation is greater than 0 |
| 15 | Durbin-Watson test<br>data: VIDEO.15 ~ Tiempo<br>DW = 0.4962, p-value < 2.2e-16<br>alternative hypothesis: true autocorrelation is greater than 0 | 16 | Durbin-Watson test<br>data: VIDEO.16 ~ Tiempo<br>DW = 0.1401, p-value < 2.2e-16<br>alternative hypothesis: true autocorrelation is greater than 0 |
| 17 | Durbin-Watson test<br>data: VIDEO.17 ~ Tiempo<br>DW = 0.6099, p-value < 2.2e-16<br>alternative hypothesis: true autocorrelation is greater than 0 | 18 | Durbin-Watson test<br>data: VIDEO.18 ~ Tiempo<br>DW = 0.7294, p-value < 2.2e-16<br>alternative hypothesis: true autocorrelation is greater than 0 |
| 19 | Durbin-Watson test<br>data: VIDEO.19 ~ Tiempo<br>DW = 0.7425, p-value < 2.2e-16<br>alternative hypothesis: true autocorrelation is greater than 0 | 20 | Durbin-Watson test<br>data: VIDEO.20 ~ Tiempo<br>DW = 0.5276, p-value < 2.2e-16<br>alternative hypothesis: true autocorrelation is greater than 0 |
| 21 | Durbin-Watson test<br>data: VIDEO.21 ~ Tiempo<br>DW = 0.3901, p-value < 2.2e-16<br>alternative hypothesis: true autocorrelation is greater than 0 | 22 | Durbin-Watson test<br>data: VIDEO.22 ~ Tiempo<br>DW = 0.5172, p-value < 2.2e-16<br>alternative hypothesis: true autocorrelation is greater than 0 |
| 23 | Durbin-Watson test<br>data: VIDEO.23 ~ Tiempo<br>DW = 0.3401, p-value < 2.2e-16<br>alternative hypothesis: true autocorrelation is greater than 0 | 24 | Durbin-Watson test<br>data: VIDEO.24 ~ Tiempo<br>DW = 0.504, p-value < 2.2e-16<br>alternative hypothesis: true autocorrelation is greater than 0 |
| 25 | Durbin-Watson test<br>data: VIDEO.25 ~ Tiempo<br>DW = 0.7982, p-value < 2.2e-16<br>alternative hypothesis: true autocorrelation is greater than 0 | 26 | Durbin-Watson test<br>data: VIDEO.26 ~ Tiempo<br>DW = 0.4709, p-value < 2.2e-16<br>alternative hypothesis: true autocorrelation is greater than 0 |
| 27 | Durbin-Watson test<br>data: VIDEO.27 ~ Tiempo<br>DW = 0.5026, p-value < 2.2e-16<br>alternative hypothesis: true autocorrelation is greater than 0 | 28 | Durbin-Watson test<br>data: VIDEO.28 ~ Tiempo<br>DW = 0.3671, p-value < 2.2e-16<br>alternative hypothesis: true autocorrelation is greater than 0 |
| 29 | Durbin-Watson test<br>data: VIDEO.29 ~ Tiempo<br>DW = 0.2727, p-value < 2.2e-16<br>alternative hypothesis: true autocorrelation is greater than 0 | 30 | Durbin-Watson test<br>data: VIDEO.30 ~ Tiempo<br>DW = 0.4378, p-value < 2.2e-16<br>alternative hypothesis: true autocorrelation is greater than 0 |
| 31 | Durbin-Watson test<br>data: VIDEO.31 ~ Tiempo<br>DW = 0.3137, p-value < 2.2e-16<br>alternative hypothesis: true autocorrelation is greater than 0 | 32 | Durbin-Watson test<br>data: VIDEO.32 ~ Tiempo<br>DW = 0.4782, p-value < 2.2e-16<br>alternative hypothesis: true autocorrelation is greater than 0 |
| 33 | Durbin-Watson test<br>data: VIDEO.33 ~ Tiempo<br>DW = 0.2124, p-value < 2.2e-16<br>alternative hypothesis: true autocorrelation is greater than 0 | 34 | Durbin-Watson test<br>data: VIDEO.34 ~ Tiempo<br>DW = 0.4572, p-value < 2.2e-16<br>alternative hypothesis: true autocorrelation is greater than 0 |
| 35 | Durbin-Watson test<br>data: VIDEO.35 ~ Tiempo | 36 | Durbin-Watson test<br>data: VIDEO.36 ~ Tiempo |
|  | DW = 0.2382, p-value < 2.2e-16<br>alternative hypothesis: true autocorrelation is greater than 0 |  | DW = 0.3834, p-value < 2.2e-16<br>alternative hypothesis: true autocorrelation is greater than 0 |
| Control A | Durbin-Watson test<br>data: CONT.2 ~ Tiempo<br>DW = 0.6094, p-value < 2.2e-16<br>alternative hypothesis: true autocorrelation is greater than 0 | Control B | Durbin-Watson test<br>data: CONT.4 ~ Tiempo<br>DW = 0.378, p-value < 2.2e-16<br>alternative hypothesis: true autocorrelation is greater than 0 |
| Control C | Durbin-Watson test<br>data: CONT.1 ~ Tiempo<br>DW = 0.4028, p-value < 2.2e-16<br>alternative hypothesis: true autocorrelation is greater than 0 | Control D | Durbin-Watson test<br>data: CONT.3 ~ Tiempo<br>DW = 0.3015, p-value < 2.2e-16<br>alternative hypothesis: true autocorrelation is greater than 0 |

### S3 DATA

Run sequence plot, ACF and PACF for the time series of Video 1 and Video 2

#### Biospeckle laser digital image processing for quantitative and statistical evaluation of the activity of Ciprofloxacin on *Escherichia coli* K-12

Hilda Cristina Grassi, Ana Velásquez, Olga Mercedes Belandria, María Lorena Lobo-Sulbarán, Jesús E. Andrades-Grassi, Humberto Cabrera, Efrén D. J. Andrades

**VIDEO 1. Culture medium with the lowest added Ciprofloxacin concentration (7.63E-04 μg/mL in well), without bacteria. ACF: Autocorrelation function; PACF: Partial autocorrelation function.**

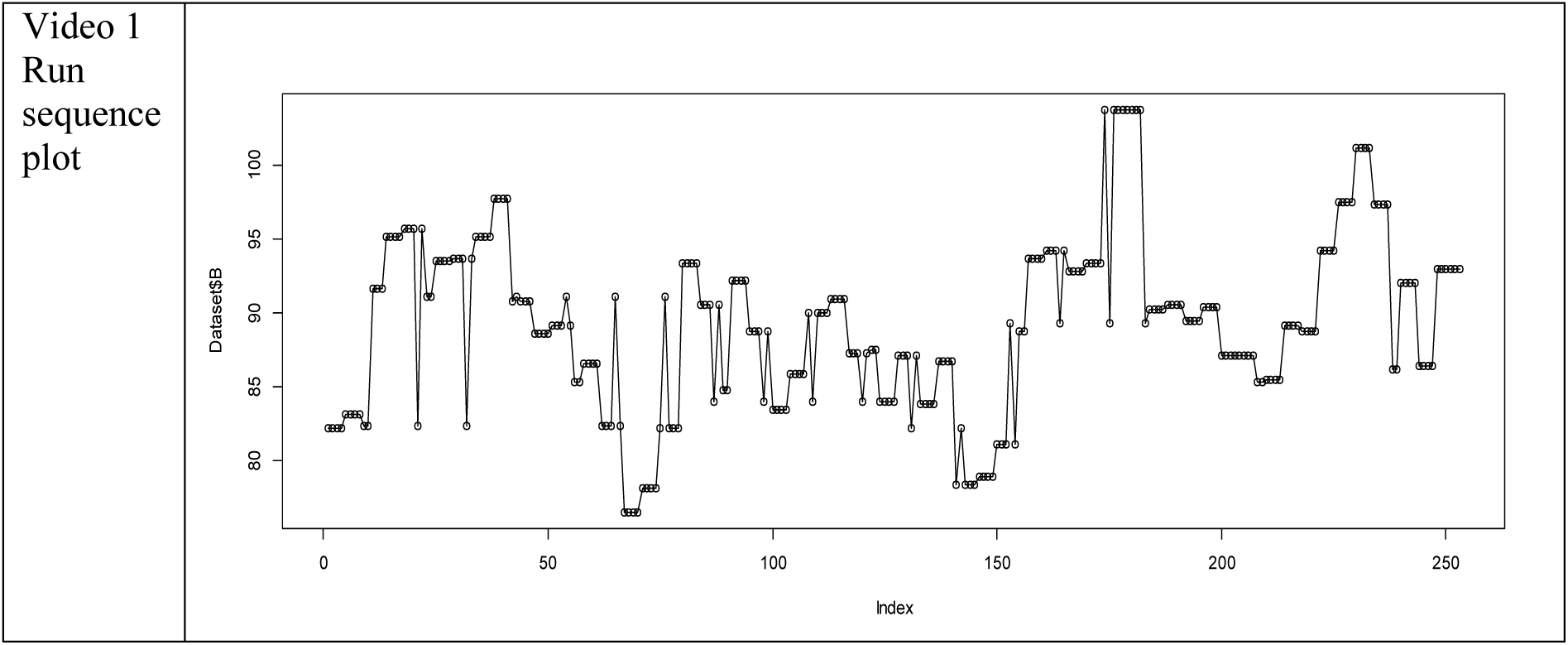
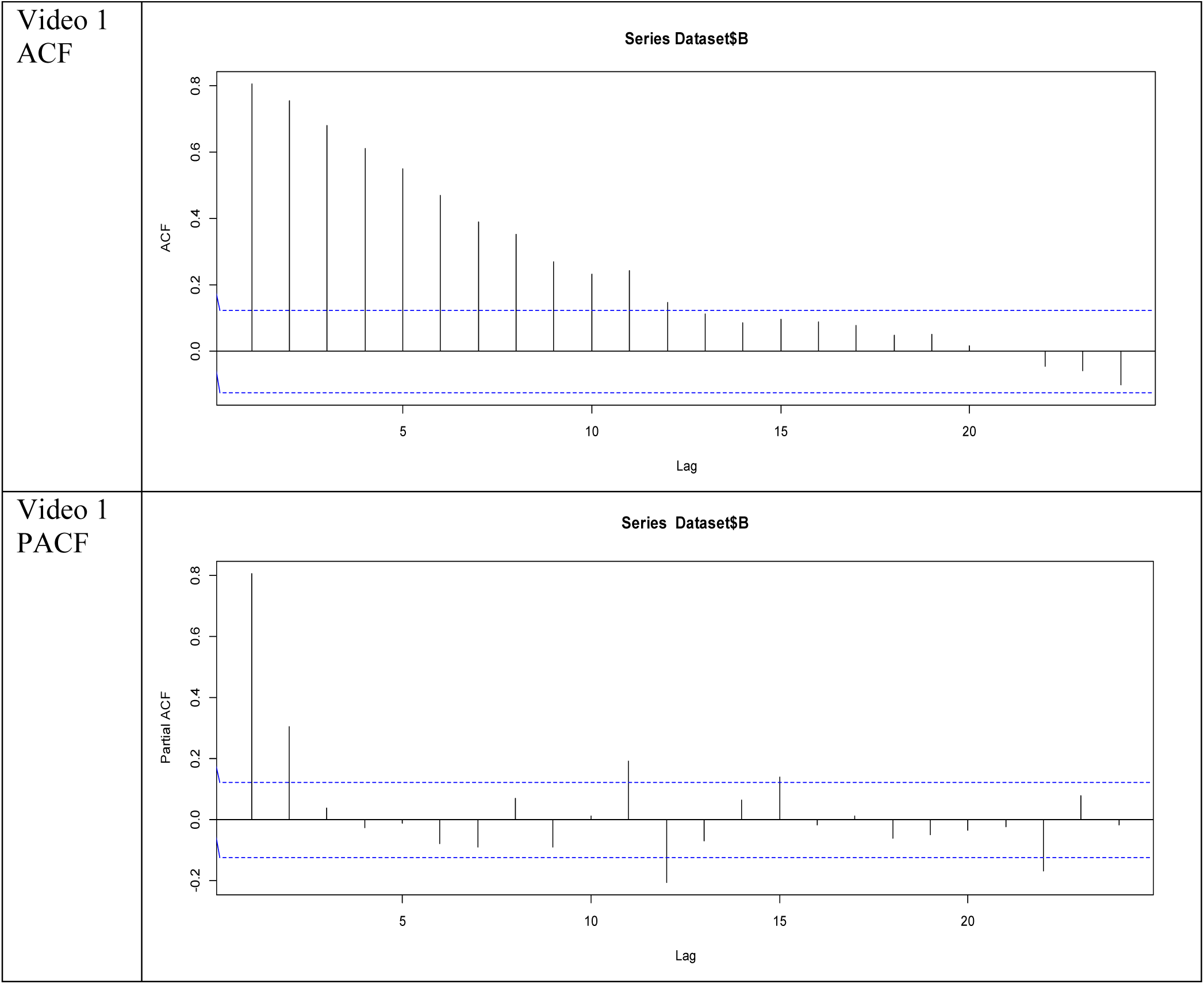

**VIDEO 2. Culture medium with the lowest added Ciprofloxacin concentration (7.63E-04 μg/mL in well), with bacteria. ACF: Autocorrelation function; PACF: Partial autocorrelation function.**

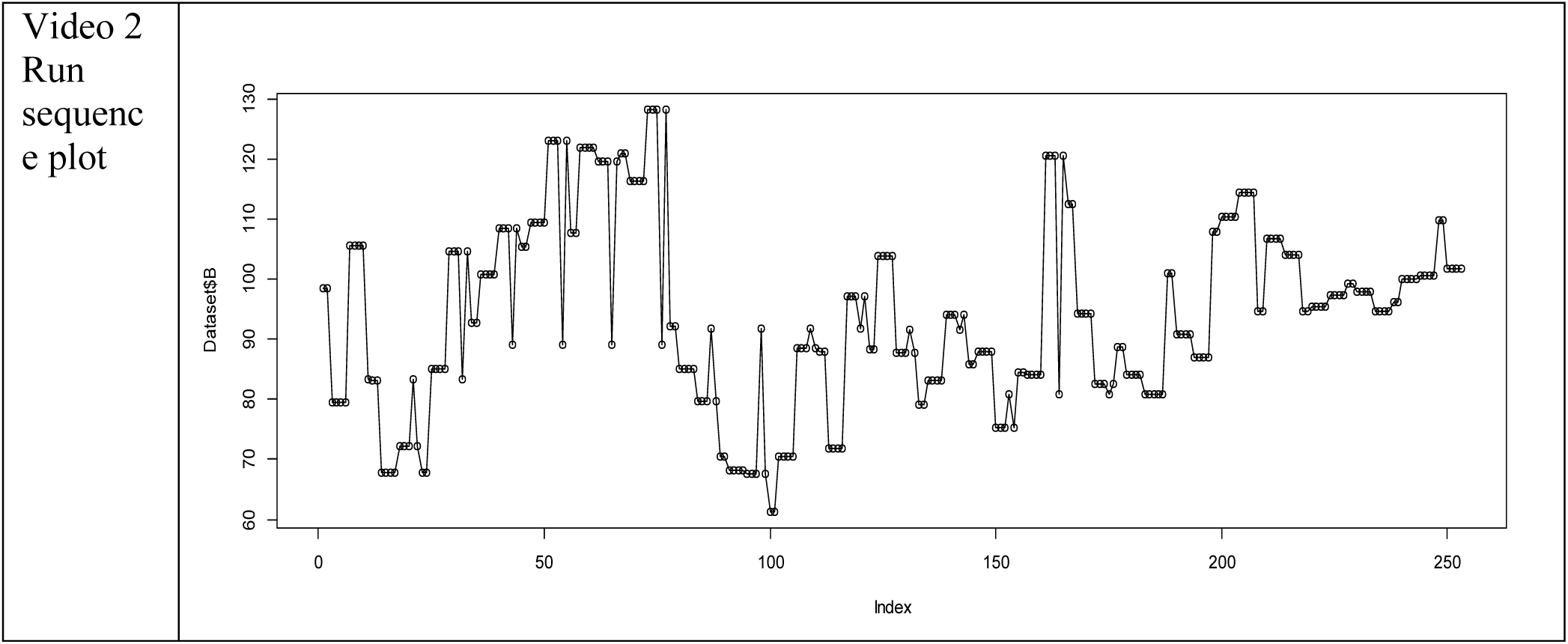
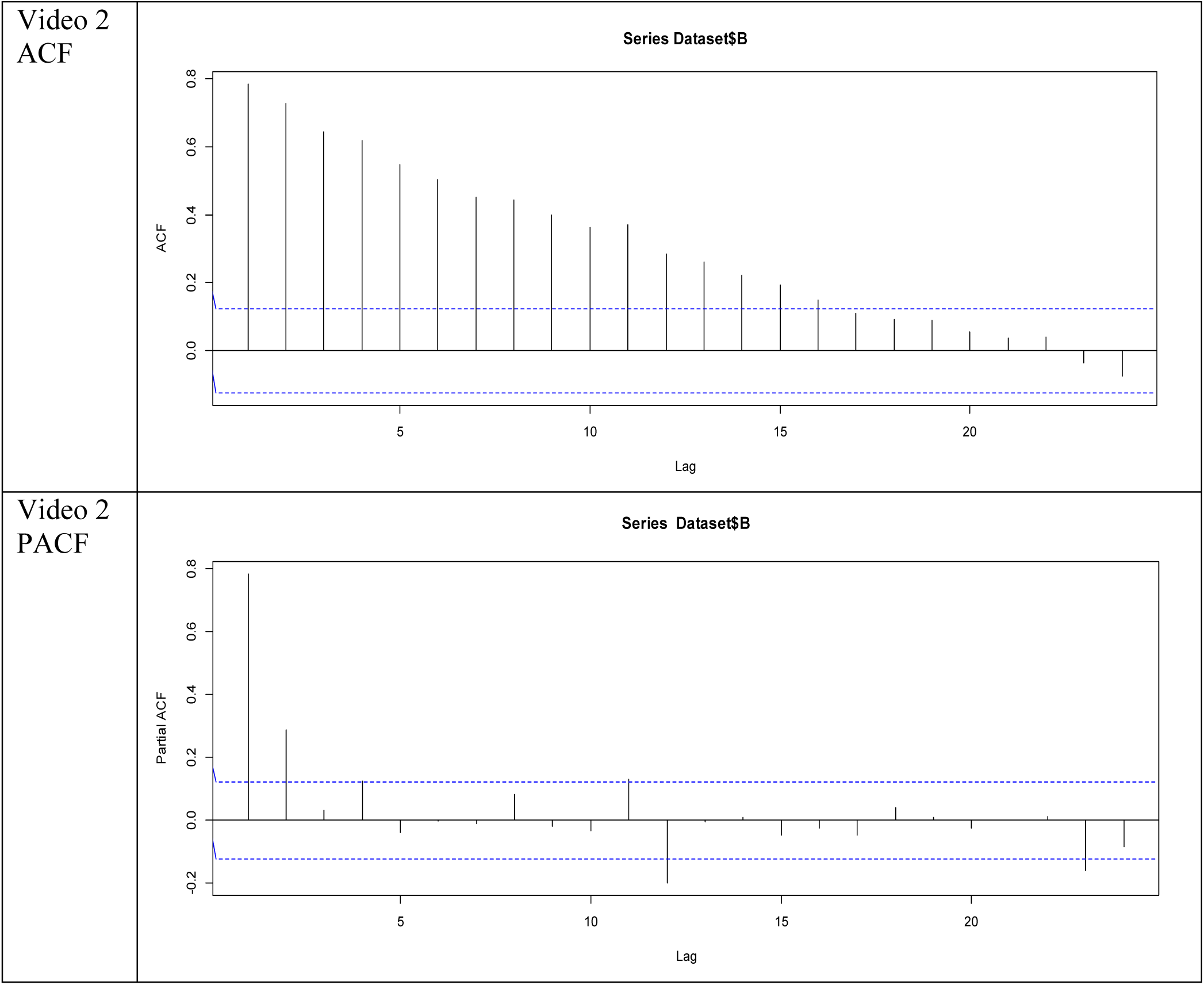

### S4 DATA

Antimicrobial Susceptibility Testing

#### Biospeckle laser digital image processing for quantitative and statistical evaluation of the activity of Ciprofloxacin on *Escherichia coli* K-12

Hilda Cristina Grassi, Ana Velásquez, Olga Mercedes Belandria, María Lorena Lobo-Sulbarán, Jesús E. Andrades-Grassi, Humberto Cabrera, Efrén D. J. Andrades

##### 1. Broth macrodilution test for microbial susceptibility. Individual values

**1. Broth macrodilution test for microbial susceptibility. Individual values.**
| Ciprofloxacin concentration | Absorbance (625nm) | UFC/mL |
| --- | --- | --- |
| 2.50E+02 | 0 | 0 |
| 1.25E+02 | 0.066 | 1.10E+08 |
| 6.25E+01 | 0.067 | 1.12E+08 |
| 3.125E+01 | 0.068 | 1.13E+08 |
| 1.563E+01 | 0.070 | 1.17E+08 |
| 7.813E+00 | 0.071 | 1.18E+08 |
| 3.906E+00 | 0.072 | 1.20E+08 |
| 1.953E+00 | 0.074 | 1.23E+08 |
| 9.766E-01 | 0.075 | 1.25E+08 |
| 4.883E-01 | 0.076 | 1.27E+08 |
| 2.441E-01 | 0.08 | 1.33E+08 |
| 1.221E-01 | 0.101 | 1.68E+08 |
| 6.104E-02 | 0.106 | 1.77E+08 |
| 3.052E-02 | 0.201 | 3.35E+08 |
| 1.526E-02 | 0.210 | 3.50E+08 |

###### 2. Broth macrodilution test for microbial susceptibility. Values organized in groups

**2. Broth macrodilution test for microbial susceptibility. Values organized in groups.**
| GROUP | CIPROFLOXACIN CONCENTRATION RANGE mg/mL | MEAN OF BACTERIAL GROWTH (UFC/mL) | STANDARD DEVIATION OF BACTERIAL GROWTH (UFC/mL) |
| --- | --- | --- | --- |
| Low concentration of Ciprofloxacin | 1.53E-02 to 2.44E-01 | 9.03E+07 | 3.46E+06 |
| Intermediate concentration of Ciprofloxacin | 4.88E-01 to 7.81E+00 | 1.23E+08 | 4.94E+06 |
| High concentration of Ciprofloxacin | 1.56E+01 to 2.50E+02 | 2.33E+08 | 9.82E+07 |

###### 3. Agar diffusion test for microbial susceptibility

The results are expressed as the inhibition diameter and the % growth of the bacteria. 100% growth (0% inhibition) corresponds with a diameter of 50mm (internal diameter of a small petri dish). A Ciprofloxacin concentration of 2.44E-03 μg/5uL corresponds with 4.88E-01 μg/mL is the minimal concentration that promotes a precise inhibition halo which can be detected and measured under the magnifying microscope.

| Group according to Ciprofloxacin concentration | Ciprofloxacin concentration µg/5uL | Inhibition diameter (mm) | Growth (%) | Mean (%) | Standard Deviation (%) |
| --- | --- | --- | --- | --- | --- |
| Low | 7.63E-05 | 0 | 100 | 100 | 0 |
| Low | 1.53E-04 | 0 | 100 |  |  |
| Low | 3.05E-04 | 0 | 100 |  |  |
| Low | 6.10E-04 | 0 | 100 |  |  |
| Low | 1.22E-03 | 0 | 100 |  |  |
| Intermediate | 2.44E-03 | 3 | 94 | 81.2 | 10.64 |
| Intermediate | 4.88E-03 | 5 | 90 |  |  |
| Intermediate | 9.77E-03 | 10 | 80 |  |  |
| Intermediate | 1.95E-02 | 14 | 72 |  |  |
| Intermediate | 3.91E-02 | 15 | 70 |  |  |
| High | 7.81E-02 | 16 | 68 | 56 | 9.27 |
| High | 1.56E-01 | 19 | 62 |  |  |
| High | 3.13E-01 | 23 | 54 |  |  |
| High | 6.25E-01 | 24 | 52 |  |  |
| High | 1.25E00 | 28 | 44 |  |  |
| Very high | 2.50E00 | 30 | 40 | NCIC | NCIC |
| Very high | 5.00E00 | 30 | 40 | NCIC | NCIC |
| Very high | 10.00E00 | 31 | 38 | NCIC | NCIC |

NCIC: Data not considered in calculation, in order to maintain 5 samples per group.

## References

1. CLSI. Performance Standards for Antimicrobial Susceptibility Testing; Twenty-Fifth Informational Supplement. CLSI document M100-S25. Wayne, PA: Clinical and Laboratory Standards Institute; 2015

2. Waldeisen JR, Wang T, Mitra D, Lee LP (2011) A Real-Time PCR Antibiogram for Drug-Resistant Sepsis. PLoS ONE 6(12): e28528. https://doi.org/10.1371/journal.pone.0028528

3. Hansen WL, Beuving J, Verbon A, Wolffs PF (2012) One-day Workflow Scheme for Bacterial Pathogen Detection and Antimicrobial Resistance Testing from Blood Cultures. J. Vis. Exp. (65), e3254, DOI:10.3791/3254

4. Liu T, Lu Y, Gau V, Liao JC, Wong PK (2014) Rapid Antimicrobial Susceptibility Testing with Electrokinetics Enhanced Biosensors for Diagnosis of Acute Bacterial Infections. Annals of Biomedical Engineering 42(11) 2314–2321. doi:10.1007/s10439-014-1040-.

5. Mohan R, Mukherjee A, Sevgen SE, Sanpitakseree Ch, Lee J, Schroeder ChM, Kenis PA (2013) A multiplexed microfluidic platform for rapid antibiotic susceptibility testing. Biosensors and Bioelectronics 49, 118–125. http://dx.doi.org/10.1016/j.bios.2013.04.046

6. Liul Ch-Y, Han Y-Y, Shih P-H, Lian W-N, Wang H-H, Lin Ch-H, Hsueh P-R, Wang J-K, Wang Y-L (2016) Rapid bacterial antibiotic susceptibility test based on simple surface-enhanced Raman spectroscopic biomarkers. Scientific Reports | 6:23375 | doi:10.1038/srep23375. www.nature.com/scientificreports.

7. Martí-López L, Cabrera H, Martínez-Celorio RA, González-Peña R (2010) Temporal difference method for processing dynamic speckle patterns. Optics Communications 283, 4972–4977.

8. Sendra H, Murialdo S, Passoni L (2007) Dynamic laser speckle to detect motile bacterial response of Pseudomonas aeruginosa. Journal of Physics: Conference Series 90 (2007) 012064 doi:10.1088/1742-6596/90/1/01206.

9. González-Peña RJ, Braga Jr. RA, Cibrián RM, Salvador-Palmer R, Gil-Benso R, San Miguel T (2014) Monitoring of the action of drugs in melanoma cells by dynamic laser speckle. Journal of Biomedical Optics 19(5), 057008. doi:10.1117/1.JBO.19.5.05700.

10. Ramírez-Miquet EE, Cabrera H, Grassi HC, Andrades EdJ, Otero I, Rodríguez D, Darias JG (2017) Digital imaging information technology for biospeckle activity assessment relative to bacteria and parasites. Lasers Med Sci doi:10.1007/s10103-017-2256-.

11. Cojoc D, Finaurini S, Livshits P, Gur, E, Shapira A, Mico V, Zalevsky Z (2012) Toward fast malaria detection by secondary speckle sensing microscopy. Biomedical Optics Express 3(5)993.

12. Sendra GH, DaiPra AL, Passoni LI, Arizaga R, Rabal HJ, Trivi M. (2010) Biospeckle descriptors: a performance comparison. Proc. of SPIE 7387, 73871K · doi:10.1117/12.87068.

13. Long J, Zucker SW, Emonet T (2017) Feedback between motion and sensation provides nonlinear boost in run-and-tumble navigation. PLoS Comput Biol 13(3): e1005429. doi:10.1371/journal.pcbi.100542.

14. Drlica K, Zhao X (1997) DNA Gyrase, Topoisomerase IV, and the 4-Quinolones. Microbiology and Molecular Biology Reviews 61(3)377–392

15. Pommier Y, Leo E, Zhang HL, Marchand C (2010) DNA Topoisomerases and Their Poisoning by Anticancer and Antibacterial Drugs. Chemistry & Biology 170. DOI 10.1016/j.chembiol.2010.04.012

16. Grassi HC, García LC, Lobo-Sulbarán ML, Velásquez A, Andrades-Grassi FA, Cabrera H, Andrades-Grassi JE, Andrades EDJ (2016) Quantitative Laser Biospeckle Method for the Evaluation of the Activity of Trypanosoma cruzi Using VDRL Plates and Digital Analysis. PLoS Negl Trop Dis 10(12): e0005169. doi:10.1371/journal. pntd.000516.

17. Wooldridge JM (2013) Econometrics, A Modern Approach, 5^th^ Edition. South-Western, Cengage Learning, USA. Library of Congress Control Number: 2012945120, ISBN-13: 978-1-111-53104-1, ISBN-10: 1-111-53104-8

18. Meschino G, Murialdo S, Passoni L, Rabal H, Trivi M (2010) Biospeckle Image Stack Process based on Artificial Neural Networks. Annual International Conference of the IEEE Engineering in Medicine and Biology Society. IEEE Engineering in Medicine and Biology Society. Conference. doi:10.1109/IEMBS.2010.562762.

19. Passoni LI, Dai Pra A, Rabal H, Arizaga R (2005) Dynamic speckle processing using wavelets based entropy. Optics Communications 246(1-3):219–228. doi:10.1016/j.optcom.2004.10.05.

20. Zdunek A, Adamiak A, Pieczywek PM, Kurenda A (2014) The biospeckle method for the investigation of agricultural crops: A review. Optics and Lasers in Engineering 52(2014)276–285. http://dx.doi.org/10.1016/j.optlaseng.2013.06.017

21. Hayes W (1953) The mechanism of genetic recombination in Escherichia coli. Cold Spring Harb. Symp. Quant. Biol. 18, 75–93. doi:10.1101/SQB.1953.018.01.01.

22. Isenberg HD (1992) Clinical Microbiology Procedures Handbook. American Society for Microbiology, USA. ISBN 1555810381, 9781555810382

23. Shapiro SS, Wilk MB (1965) An analysis of variance test for normality (complete samples). Biometrika 52(3–4)591–611. doi:10.1093/biomet/52.3-4.59.

24. Breusch, T.S. and Pagan, A.R. (1979) A Simple Test for Heteroscedasticity and Random Coefficient Variation. Econometrica, 47(5)1287–1294. http://dx.doi.org/10.2307/1911963

25. Durbin, J.; Watson, G. S. (1950). “Testing for Serial Correlation in Least Squares Regression, I”. Biometrika. 37 (3–4): 409–428. doi:10.1093/biomet/37.3-4.40.

26. Durbin, J.; Watson, G. S. (1951). “Testing for Serial Correlation in Least Squares Regression, II”. Biometrika. 38 (1–2): 159–179. doi:10.1093/biomet/38.1-2.15.

27. Box GEP, Jenkins G (1976). Time Series Analysis: Forecasting and Control. Holden-Day, USA.

28. Balouiri M, Sadiki M, Ibnsouda SK (2016) Methods for in vitro evaluating antimicrobial activity: A review. Journal of Pharmaceutical Analysis 6 (2016)71–79. http://dx.doi.org/10.1016/j.jpha.2015.11.005

29. Braga Jr. RA, González-Peña RJ, Campos Viana D, Pujaico Rivera F (2017) Dynamic laser speckle analyzed considering inhomogeneities in the biological sample. Journal of Biomedical Optics 22(4), 045010. 1083-3668/2017

30. Nothdurft R, Yao G (2005) Imaging obscured subsurface inhomogeneity using laser speckle. Optics Express 13(25)10034.

31. Berg HC, Brown DA (1972) Chemotaxis in Escherichia coli analysed by three-dimensional tracking. Nature 239(5374)500 – 504. doi:10.1038/239500a.

32. Benisty S, Ben-Jacob E, Ariel G, Be’er A (2015) Antibiotic-Induced Anomalous Statistics of Collective Bacterial Swarming. Phys. Rev. Lett. 114, 018105. doi:10.1103/PhysRevLett.114.01810.

